# Climate change drives northward range shifts in invasive forest pests and pathogens

**DOI:** 10.64898/2026.11.06.764783

**Authors:** Haoran Wu, Jiqi Gu, Chengzhang Zheng, Yadvinder Malhi, Cecilia A. L. Dahlsjö

**Author notes:** Corresponding author: Haoran Wu. **Data and code availability:** Data and analysing code for the manuscript is available at zenodo https://zenodo.org/records/22843856.

## Abstract

Invasive tree pests and pathogens are increasingly threatening forests worldwide. Understanding how a warming climate and changing precipitation patterns will affect the future distribution ranges of these pathogenic species requires systematic analysis across a variety of host trees and pathogenic species. We analysed 31 major tree pests and pathogens that have caused widespread and extensive damage globally, predicting their current and future distribution under various climate change scenarios using species distribution modelling. Our results showed that tree pests and pathogens will consistently shift northward, threatening northern temperate and boreal forests between the years 2041 and 2080. On the other hand, some warmer regions, including southern Europe, were expected to experience fewer pest and pathogen outbreaks in the future. Annual mean temperature, mean temperature of the warmest quarter, precipitation of the warmest quarter, and precipitation of the driest month contributed to the species distribution models, indicating that warming temperatures and moderate precipitation may promote the survival, reproduction, and spread of the pathogenic agents. Our research highlights the need for enhanced monitoring, early pest and pathogen detection, stricter biosecurity measures, and improved silvicultural strategies in temperate and boreal forests in the northern hemisphere. Moreover, forest assessments, including those focused on carbon budgets, biodiversity, and ecosystem services, should account for the effects of tree pests and pathogen outbreaks, as they may drastically alter future functioning of forests globally.

**Statement of authorship:** HW designed and implemented the project and wrote the manuscript with significant contributions and advice from JG and CALD. CZ implemented the visualisations of the model. The project is supervised by CALD and YM.

## 2. Methods

## 1. Introduction

Tree pests and pathogens are natural components that regulate host tree population dynamics, forest nutrient cycles, and overall forest health. However, due to increasing global change and human trade, some pests and pathogens have moved beyond their native ranges, where they cause severe tree diseases, tree mortality events, and subsequent carbon loss, resulting in substantial economic losses and damage to ecosystem services (Boyd et al. 2013; Fei et al. 2019; Gougherty and Davies 2022).

Currently, new records of emerging forest diseases are doubling approximately every 11 years (Gougherty 2023), with e.g., Ash Dieback and Dutch Elm Disease continuing to raise concerns (Copeland et al. 2023; Carroll 2024).

Despite their global prevalence, numerous studies suggest that tree pests and pathogens appear to be particularly destructive to northern temperate and boreal forests (Fei et al. 2019; Venäläinen et al. 2020; Forzieri et al. 2023; Wang et al. 2023). Boreal forests account for approximately 32% of the carbon sequestered in the world’s forests (Pan et al. 2011), and their biomass is predicted to increase due to global warming, highlighting their importance in global timber production, forest management, and carbon policies (Xu et al. 2024). However, these forest ecosystems are relatively simple, often dominated by only a few species, which makes them especially susceptible to large-scale disease outbreaks. Tropical forests, on the other hand, have higher biodiversity which has been shown to reduce the effects of tree pests and pathogens (Ciesla 2001).

Warming trends and milder winters enable pests and pathogens to expand their ranges, reproduce more frequently, and survive in regions previously protected by cold winters (Goberville et al. 2016; Goodsman et al. 2018; Szyniszewska et al. 2024). Furthermore, forests in northern temperate and boreal zones are major hubs for timber production and international trade (Hashiramoto et al. 2004), which further promotes the spread of pests and pathogens (Ramsfield et al. 2016). It is therefore vital to assess how tree pests and pathogens are likely to spread in northern regions and to identify the most vulnerable regions.

Although studies have projected the future distribution of individual forest pests or pathogens (e.g. Watt et al. 2009; Goberville et al. 2016; Aierken et al. 2024), the cumulative climate-driven redistribution of multiple high-impact species remains poorly understood. Such a synthesis is essential as simultaneous range expansions across taxa may generate emergent disturbance regimes that disproportionately threaten carbon-rich northern temperate and boreal forests.

Here, we analyse the geographic distributions of 31 high-impact tree pests and pathogens responsible for widespread tree mortality. Using ensemble climate projections under multiple Shared Socioeconomic Pathways (SSPs), we (1) estimate their current potential distributions, and (2) quantify projected shifts in cumulative pest and pathogen pressure through the mid- and late-21st century. By identifying regions where multiple agents are likely to converge, we aim to assess the vulnerability of global forests to future biologically driven disturbance under climate change.

### 2.1. Pest and pathogen selection

In this study, 31 tree-infecting pests and pathogens were evaluated (see Supplementary Table S1 for details). The compilation of this pest or pathogen list followed dual criteria: (1) the organism is documented to cause extensive, widespread, or economically important damage in at least one country or region, based on peer-reviewed literature, forest health surveys, or official reports from national forestry agencies, and (2) the organism has been formally recognised or prioritised by relevant authorities, being designated as a regulated pest or pathogen, quarantine organism, notifiable pathogen, or listed species on official forest health databases. Selection was done through a systematic review of pest and pathogen inventories from authoritative sources across major regions, including:

- The “Pest and Disease Resources” registry by the UK Forest Research
- The “Key Tree Pests and Diseases” registry by the UK Woodland Trust
- The “Terrestrial Pathogens and Diseases” registry by the U.S. Department of Agriculture (USDA)
- The “Top Forest Insects and Diseases” registry by the Government of Canada
- The “Forest Damage Caused by Pests and Diseases” registry by the Japanese Forestry Agency
- The “National Survey of Forestry Harmful Organisms” by China’s National Forestry and Grassland Administration

From these sources, we selected organisms that met our predefined criteria. While the consulted sources do not cover all countries globally, many of the documented pests and pathogens extend their damage beyond the countries hosting the source websites, thereby ensuring regional or global representativeness. For instance, the Pine wood nematode (*Bursaphelenchus xylophilus*) appears on the UK Forest Research list, although its most severe impacts have been documented in East Asia. In addition, though this list may miss organisms that cause damages in many Global South countries, it still covers major pests and pathogens in temperate and boreal forests of the northern hemisphere, and therefore, it allows us to accurately estimate the geographic shift of pests and pathogens in this region, which is the key focus of this paper.

The complete list of pests and pathogens included in the analysis is provided in Supplementary Table S1.

### 2.2. Pest and Pathogen Distribution Data

To model the geographic areas affected by these pests and pathogens, we compiled presence data from a range of sources, including peer-reviewed literature, government agency databases (e.g., the U.S. Alien Forest Pest Explorer), the Global Biodiversity Information Facility (GBIF), and the EPPO Global Database. Priority was given to literature that synthesises multiple data sources to generate comprehensive pest and pathogen presence datasets (see Supplementary Table S1 for details on these literary sources), with GBIF occurrence points used to supplement these primary datasets. For pests and pathogens lacking robust literature sources, GBIF occurrence data served as the primary presence dataset. All presence points were subsequently cross-validated against state- or province-level records of pest and pathogen presence from the EPPO database. A detailed account of the data collection process for each pest or pathogen is provided in Supplementary Table S1.

Some locations with recorded pest or pathogen presence may experience only minimal forest damage. For instance, the emerald ash borer (*Agrilus planipennis*) is highly destructive to several ash species (e.g. white ash (*Fraxinus americana*), black ash (*Fraxinus nigra*), green ash (*Fraxinus pennsylvanica*)) in North America and common ash (*Fraxinus excelsior*) in Europe, but its impact in East Asia, its native area, is limited due to long-term co-evolution with its host tree species there. In this paper, we excluded presence points where the pest or pathogen was documented to co-evolve with its native hosts and do not cause destructive epidemics, evidenced by peer-reviewed research papers, official websites, and EPPO database.

### 2.3. Climatic Data

We used the WorldClim version 2 bioclimatic variables, with a spatial resolution of 10 arc-minutes. The dataset represents average values for the period 1970–2000. The definitions of WorldClim bioclimatic variables are displayed in Supplementary Table S2.

To project the future range of pests and pathogens, WorldClim future climate data was used. The dataset consists of downscaled future climate projections from the Coupled Model Intercomparison Project Phase 6 (CMIP6). We focused on 10 arc-minute resolution bioclimatic variables for two time periods: 2041–2060 and 2061–2080.

Projection datasets were downloaded for four Shared Socio-economic Pathways (SSPs): SSP126 (sustainable and green pathway, with an anthropogenic radiative forcing of 2.6 W/m^2^), SSP245 (the “middle of the road” pathway with moderate global population growth, a certain degradation of ecosystems, and an anthropogenic radiative forcing of 4.5 W/m^2^), SSP370 (regional rivalry pathway, with rising inequality and an anthropogenic radiative forcing of 7 W/m^2^), and SSP585 (fossil-fuelled development, with an anthropogenic radiative forcing of 8.5 W/m^2^), and 11 global climate models (GCMs): ACCESS-CM2, BCC-CSM2-MR, CMCC-ESM2, EC-Earth3-Veg, GISS-E2-1-G, INM-CM5-0, IPSL-CM6A-LR, MIROC6, MPI-ESM1-2-HR, MRI-ESM2-0, and UKESM1-0-LL.

### 2.4. Modelling Procedure

We used the Maximum Entropy (MaxEnt) algorithm to estimate both current and future distributions of pest- or pathogen-damaged areas. The MaxEnt software version 3.4.4 (Phillips, Anderson & Schapire 2006) was used to run the algorithm. For each pest or pathogen, we first ran MaxEnt using an initial set of bioclimatic variables that could potentially best describe a pest’s or a pathogen’s ecological tolerance: bio_1 (Annual Mean Temperature), bio_5 (Max Temperature of Warmest Month), bio_6 (Min Temperature of Coldest Month), bio_12 (Annual Precipitation), bio_16 (Precipitation of Wettest Quarter), bio_17 (Precipitation of Driest Quarter). The annual mean temperature and annual precipitation are approximates of the pests’ and pathogens’ suitable ecological niche to thrive. The maximum temperature of warmest month and precipitation of driest quarter represent heat and drought stress that could reduce the sporulation of pathogen species, reproduction success of pest species, but may also increase host tree vulnerability. The minimum temperature of coldest month represents cold stress in the winter, which defines the northern boundary for some pests and pathogens to survive. Precipitation of wettest quarter represents the moisture condition, which is essential for some pathogens to sporulate, spread, and infect new trees.

With the result from the initial MaxEnt model available, we then selected bioclimatic variables from them for a refined model based on four criteria:

1. Variables must have contributed at least 5% to the initial MaxEnt model incorporating six bioclimatic variables.
2. Selected variables must exhibit a Pearson correlation coefficient below 0.8 to avoid collinearity.
3. Each model must include no more than four variables, as additional variables do not significantly improve model accuracy (Goberville et al. 2016).
4. Selected variables should generally be supported by evidence from existing literature demonstrating that they constrain the biological performance of the pest or the pathogen. When such evidence is unavailable, variable contribution and permutation importance in the initial MaxEnt model are used as the primary selection criteria.

Using the filtered bioclimatic variables, we re-ran a refined MaxEnt model for each pest or pathogen to predict its current distribution range. We then projected the effects of climate change on future pest and pathogen ranges for the periods 2041–2060 and 2061–2080 under the four SSPs and 11 GCMs, utilising the WorldClim future climate dataset. For each SSP, the future pest or pathogen range was determined by averaging prediction values across the 11 GCMs.

To address uneven sampling efforts in GBIF and other occurrence datasets, we used a bias-corrected sampling approach to generate background points for the MaxEnt model. Specifically, we sampled 10,000 background points from a bias raster representing spatial variation in sampling effort, using probability-weighted random sampling. The bias raster was created based on synthesised data from the literature. In our previous work, we synthesised plot-based monitoring data of tree pests and pathogens from 472 peer-reviewed papers, selected from an initial pool of 16,355 (Chapter 4). 12,933 field survey records, spanning 1953 to 2024 and a wide range of forest pests and pathogens are used to reflect the scientific sampling intensity across the globe.

To evaluate the model performance, the presence data were split into two subsets: 75% of the data were used to build the model (training data); and the remaining 25% served to evaluate model performance (testing data). The Area Under the Curve (AUC) values on both training and testing data were calculated. The AUC values quantify the performance of MaxEnt models, with values ranging from 0 to 1. Higher values indicate better model accuracy. The calculated values for each pest’s or pathogen’s MaxEnt model are provided in Supplementary Table S3.

For the output format, the cloglog value (complementary log-log) was used to estimate the probability of pest or pathogen occurrence. This value represents suitability of a habitat for a pest or pathogen species.

Next, a threshold provided by MaxEnt that balances training omission and predicted area was applied to generate a binary distribution map for each pest or pathogen. These individual distribution maps were then summed to produce the final species richness map.

### 2.5. Calculate Pest and Pathogen Richness Map

To estimate pest and pathogen richness under both current and future conditions, we first applied spatial masks to each pest’s or pathogen’s prediction map to restrict calculations to relevant regions. This step was necessary because some areas predicted as suitable may correspond to regions where the pest is present but does not cause significant damage, often due to host adaptation or resistance.

Based on available information on pest- or pathogen-damaged ranges from EPPO, we restricted projections to regions within the documented invaded range to avoid unrealistic extrapolation beyond dispersal constraints. Details of the invaded range for each pest or pathogen are provided in Supplementary Table S1.

## 3. Results

The current global distributions of 31 major tree pests and pathogens that threaten European forests were successfully mapped (Figure 1, see individual distributions in Figure S1). North America (22 species) and Europe (17 species) were found to be hotspots, while East Asia harboured a sizeable number of pathogenic species (10 species) (Figure 1, see Figure S2 for the standard deviation of model projections across 11 GCMs).

**Figure 1.**
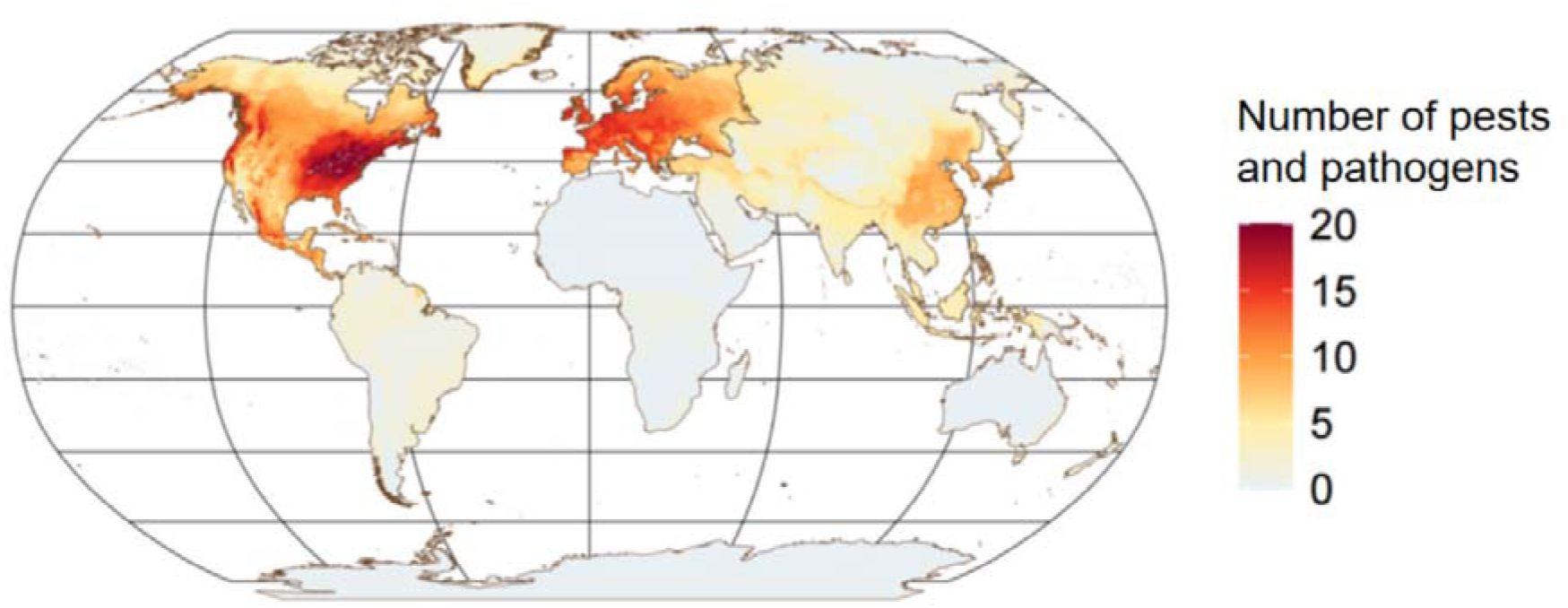
Modelled pest and pathogen species richness under current conditions. Presence-only data for 31 major forest pests and pathogens were used as input for the MaxEnt program. Species richness was calculated by summing the predicted distribution map for each pest or pathogen, where predicted presence regions are places with larger predicted probability of occurrence than a selected threshold. This balanced training omissions and predicted areas in the MaxEnt model.

Between 2041 – 2080 we found that pests and pathogens are expected to expand into new areas at a rate of 16,747 km^2^/year (SD = 9,562 km^2^/year for 11 GCMs) under the green emissions (SSP126) scenario and 59,304 km^2^/year (SD = 16,179 km^2^/year for 11 GCMs) under the fossil fuel (SSP585) scenario. Regardless of the climate scenario, eastern parts of North America are expected to experience the highest increase in the number of novel pests and pathogens from 2041 (Figure 2). The expansion in Europe is expected to take place in eastern and northern regions, while the current southeastern hotspot is expected to experience little increase (Figure 2). While the expansion of pests and pathogens in East Asia is projected to be limited in the near term (2041–2060), under a high emissions scenario (SSP585), we expect to see an expansion of the modelled species into large parts of north Asia during 2061–2080 (Figure 2). The predicted pest and pathogen richness is given in Supplementary Figure S3, and the changes in individual pest or pathogen distributions are shown in Supplementary Figure S4. When summarising the changes in pest and pathogen richness across the globe, we found that most regions will experience an increase in the number of pests and pathogens (Supplementary Figure S5).

**Figure 2.**
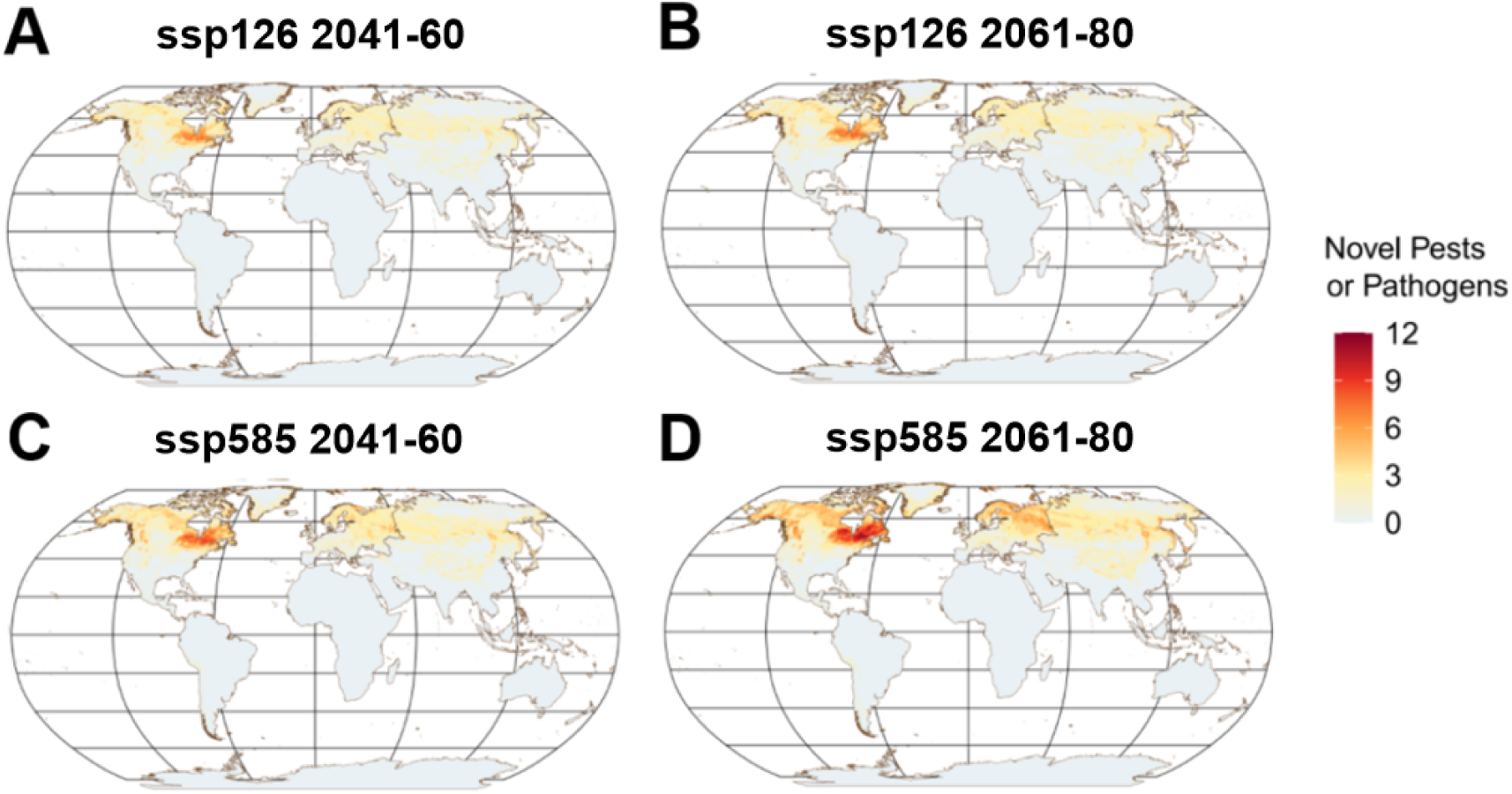
Number of newly emerged pests and pathogens (novel pests and pathogens) for the periods 2041– 2060 and 2061–2080 under the lowest (SSP126) and highest (SSP585) emission scenarios.

The six most damaging forest pests and pathogens included in our model (*Hymenoscyphus fraxineus*, *Anoplophora glabripennis*, *Ophiostoma ulmi*/*O. novo-ulmi*, *Agrilus planipennis*, *Bursaphelenchus xylophilus*, and *Cryphonectria parasitica*) all show a consistent northward expansion in suitable habitat between 2041 and 2080 (Figure 3). Across taxa, occurrence probabilities increase in northern temperate and boreal regions, including northern Europe, Canada, and parts of western Siberia, while declining in some currently affected southern regions (Figure 4). This pattern is particularly pronounced in Europe and North America. For example, *H. fraxineus* is projected to expand into northern Europe and Scotland while retreating from southeastern Europe and southern England (Supplementary Figure S4). Similarly, *A. planipennis* and *Ophiostoma* spp. show increased suitability across Canada, Alaska, and northern Europe, indicating an elevated risk to ash and elm populations in these regions. In Asia and Eurasia, *B. xylophilus* is projected to expand into Europe and western Siberia, while *C. parasitica* shows increased suitability across northern and western Europe and the United Kingdom (Supplementary Figure S4).

**Figure 3.**
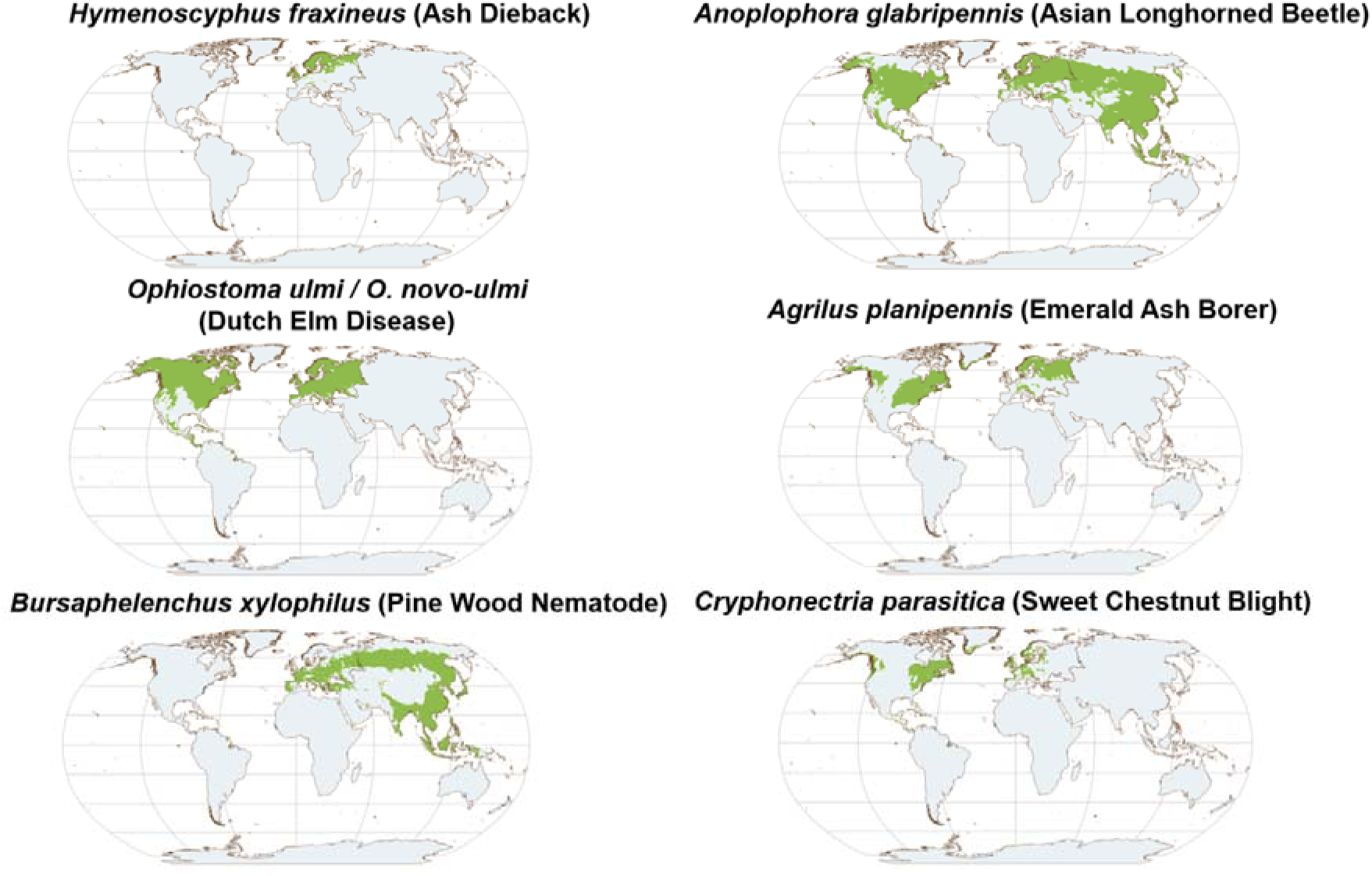
Projected distributions of six important pests or pathogens (*Hymenoscyphus fraxineus* (Ash Dieback), *Anoplophora glabripennis* (Asian Longhorned Beetle), *Ophiostoma ulmi* and *Ophiostoma novo-ulmi* (Dutch Elm Disease), *Agrilus planipennis* (Emerald Ash Borer), *Bursaphelenchus xylophilus* (Pine Wood Nematode), C*ryphonectria parasitica* (Sweet Chestnut Blight)) for 2061–2080 under scenario SSP585.

**Figure 4.**
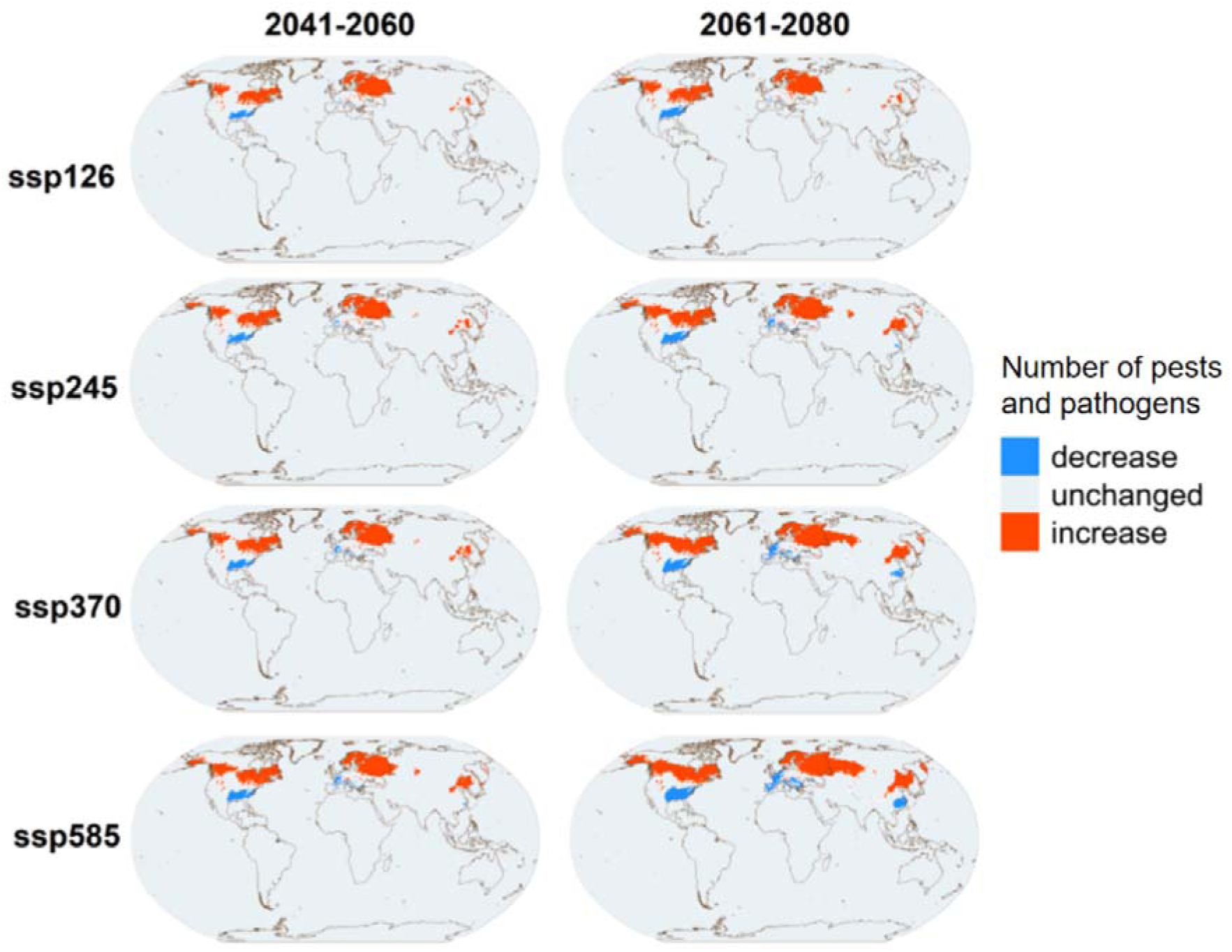
Projections of changes in the number of pests or pathogens for 2041–2060 and 2061–2080 under four Shared Socio-economic Pathways (SSPs): SSP126 (the most climate-positive scenario, characterised by low greenhouse gas emissions), SSP245, SSP370, and SSP585 (the most high-emission, climate-negative scenario). The number of pests and pathogens was calculated by summing the binary distribution maps of individual pest or pathogen species. Each binary distribution map was generated by applying a threshold that balances training omission and predicted area. A pest or pathogen was considered present at a given location if its predicted occurrence probability exceeded the corresponding threshold.

## 4. Discussion

### 4.1. Temperature and Northward Range Shifts

Across the provided 31 tree pests and pathogens, we detect a consistent poleward intensification of climatic suitability, suggesting a systemic redistribution of forest disturbance pressure toward northern temperate and boreal forests. Warming temperatures appear to be a key driver of this trend.

For insect pests, the rate of development increases with temperature, but declines rapidly beyond an upper thermal limits (Jactel et al. 2019). Warmer temperatures both shorten generation times and increase the number of offspring, facilitating insect outbreaks (Jactel et al. 2019). For fungal pathogens, warmer temperatures may favour sporulation and dispersal for some species (La Porta et al. 2008). However, temperatures above 30°C can strongly limit pathogen germination and growth. At the same time, extreme heat and drought can weaken tree health, making them more vulnerable to pest and pathogen attacks (Jactel et al. 2019; Chen et al. 2024; Tabassum et al. 2024). These patterns align with our results, which show that almost all pests and pathogens analysed have increased occurrence probabilities in northern temperate and boreal regions, while probabilities decrease in the southeastern United States, southwestern Europe, and eastern China. Although the effects of temperature on the geographical range shifts of pests and pathogens are influenced by many complex factors (Jactel et al. 2019; Singh et al. 2023), we can still predict their net effect using niche-based species distribution models.

These mechanisms are consistent with the thermal biology of many tree pests and pathogens. For instance, both insect pests such as the Asian Longhorned Beetle and fungal pathogens, such as those causing Dutch elm disease, exhibit optimal performance at moderate temperatures (Zhou et al. 1984, Zhou et al. 1984, D’Arcy 2000, Straw et al. 2015), suggesting that warming conditions will enhance their spread into northern regions. In addition, warming temperature may increase drought events, thus having an indirect effect on intensifying the disease outbreaks through weaking the host trees. For example, plantations of poplars with a shortage of water were found to be attacked by Asian Longhorned Beetle, causing serious epidemics (Luo et al. 2003). These provide a mechanistic basis for the northward shifts observed in our models.

### 4.2. Impact of Precipitation on Pest and Pathogen Range

Many fungal pathogens generally benefit from increased rainfall and high moisture, which support spore production and transmission (Milici et al. 2020). *Hymenoscyphus fraxineus*, the pathogen responsible for Ash Dieback, exemplifies this, as it requires moist spring and summer conditions for sporulation and spread across host trees (Marçais et al. 2023). Similarly, *Cryphonectria parasitica*, the causal agent of Sweet Chestnut Blight, also depends on humid conditions for growth, spore germination, and dispersal. Elevated precipitation increases splashing and water flow, potentially enhancing the efficiency of fungal spore dispersal (Metreveli et al. 2024).

For insect pests, the effects of precipitation are complex. Moderate precipitation may support pest survival, whereas both excessive dryness and excessive wetness can negatively affect their life cycles. These hydrological extremes may also damage tree health, making host trees more vulnerable to infestation (Kolb et al. 2016; Dahlsjo and Malhi 2024). Generally, pests that primarily feed on woody tissues (e.g., moths boring into pine shoots or stems) are adversely affected by drought, whereas bark beetles, leaf chewers, miners, gall makers, and sap feeders often benefit from drier conditions (Jactel et al. 2019). Our results indicates that many pests (e.g., emerald ash borer) require a moderate level of precipitation for egg hatching and larval survival. The Asian Longhorned Beetle, on the other hand, showed exceptionally high occurrence under extreme drought. This may have been due to a weakened host trees affected by water stress, although such a conclusion must be supported by empirical evidence.

### 4.3. Management Considerations

The projected northward range shifts highlight the need for strengthened forest management strategies in northern temperate and boreal regions. These include enhanced monitoring and early detection, stricter biosecurity controls on potentially infected plant material, and silvicultural practices that increase forest resilience, such as promoting tree species diversity and reducing connectivity among susceptible hosts (Dahlsjo 2023). Given that many of these pests and pathogens target economically important species, mixed-species plantations may help balance productivity with resistance to disease outbreaks.

Global forest resource assessments should also account for the effects of pests and pathogens, particularly in northern temperate and boreal zones. For example, an outbreak of spruce budworm in Canada was estimated to cause a carbon sink loss of approximately 2.87 t C ha ¹ yr ¹, shifting eastern Canadian boreal forests from a net carbon sink to a carbon source (Zhang et al. 2014). In a study of ∼92,000 field plots, mortality caused by 15 major non-native forest pests and pathogens in the United States was estimated at ∼5.53 Tg C per year (Fei et al. 2019). Similarly, pine wilt disease, a major threat to Chinese pine forests, is estimated to cause 483.23 Tg C loss even under current climate conditions (Hu et al. 2024). To accurately assess forest carbon sinks, carbon budget models should incorporate the effects of pests and pathogens to project future forest productivity (Volney et al. 2000); however, this is still a major gap in global forest carbon accounting.

Biodiversity would also be adversely affected by pest and pathogen outbreaks due to large-scale tree mortality, which alters species competition and habitat structure (Mitchell et al. 2021). For instance, in the United Kingdom, 953 species were identified as associated with *Fraxinus excelsior* (common ash), including 1 fungus, 29 invertebrates, and 4 lichens that are obligately dependent on ash trees. Thus, substantial declines of ash trees has wider implications than tree mortality alone (Mitchell et al. 2014). Assessments of other ecosystem services, such as water retention, nitrogen cycling, flood risk reduction, and cultural values, in northern temperate and boreal forests, should also be reconsidered, particularly in high pest- or pathogen-risk areas.

### 4.4. Limitations

One important limitation in this study is the temporal mismatch between bioclimatic data and the species occurrence data used for MaxEnt modelling. When the baseline periods of these datasets do not align, the reliability of species distribution models may be reduced (<u>Roubicek et al. 2010</u>). For example, <u>D’Amen et al. (2024)</u> found that the temporal resolution of input data substantially influenced estimates of thermal niche overlap between Lessepsian fish in their native and invaded ranges. In the present study, the historical WorldClim bioclimatic data represent the period 1970– 2000, whereas many of the epidemics considered in this study have caused substantial damage primarily since 2000. Examples include Ash Dieback, Beech Leaf Disease, Emerald Ash Borer, Laurel Wilt, and Spotted Lanternfly. Because historical temperatures were generally lower than those experienced in recent decades, using these historical climatic data may lead to an overestimation of the cold tolerance of pests and pathogens that emerged or became prominent after 2000. Consequently, their climatic niches may be estimated to extend into colder environments than they can actually tolerate. This could, in turn, lead to overestimation of both the magnitude and rate of their projected northward range shifts.

Other methodological constraints include: Firstly, the models rely on presence-only data, which may reflect uneven sampling effort despite bias correction. Geographic bias on the bias raster that is determined by synthesised tree pest and pathogen research data (Chapter 4) can propagate to species distribution modelling. Secondly, projections of species climatic suitability rely on the assumption that tree pests and pathogens have occupied all geographically and environmentally suitable areas. This equilbrium assumption can be violated for newly emerged pests and pathogens. Thirdly, the model does not account for host availability, which is essential for survival and spread of a pest or a pathogen. Areas that are suitable for pests and pathogens to invade should be areas where host trees exist. Fourthly, model projections are restricted in documented invaded continents, which mean that continents where a pest or a pathogen arrives but is not documented would be missed. Finally, the AUC values reported in Supplementary Table S3 should be interpreted with caution. The MaxEnt models in this study were evaluated using random partitioning of occurrence records into training and testing subsets rather than spatially blocked cross-validation. This approach may inflate estimates of model performance because of spatial autocorrelation among geographically clustered occurrence records. Accordingly, greater confidence should be placed in the multi-species consensus pattern than in the predicted distributional changes of individual taxa. As shown in Supplementary Figure 4, the predicted distributional changes of all individual pest and pathogen species exhibited a consistent signal of northward expansion, regardless of model performance (i.e. AUC values). This indicates that, even if we exclude or down-weighted models with low AUC values, the principal patterns displayed in Figure 4 remains unchanged.

Overall, our results indicate a consistent poleward redistribution of tree pests and pathogens, with increasing convergence of multiple high-impact species in northern temperate and boreal regions. This emerging pattern has important implications for forest resilience, as these ecosystems play a critical role in global carbon storage, biodiversity, and ecosystem services. Incorporating biotic disturbance agents into future forest assessments will be essential for accurately predicting ecosystem responses to climate change. Without such integration, projections of forest function and stability are likely to underestimate the risks posed by biologically driven disturbances in a warming world.

## Acknowledgements

This work was supported by the Scottish Forestry Trust/Forest Commission Joint Bursary Award Scheme (grant number: P24-320B) for the DPhil programme in Geography and the Environment at the University of Oxford. We thank Dr. Kiran Jonathan Horrocks, Dr. Eric Goberville who shared pest occurrence data and provided advice.

