## Supplementary Materials for "Climate change drives northward range shifts in invasive forest pests and pathogens"

**Table S1. Data Collection**

**Supplementary Table S1.** Description of data sources used to collect occurrence data for forest pests and pathogens that cause substantial damage to global forests.

| **Pest/pathogen** | **Scientific name** | **Data source** | **Potentially affected area** | **Description** |
| --- | --- | --- | --- | --- |
| Ash Dieback | *Hymenoscyphus fraxineus* | [1] Goberville et al. (2016); [2] Forest Comission - Chalara Map; [3] GBIF; [4] EPPO | Europe (except Iceland); Russia: Northwestern; Russia: Volga; Russia: Central | [1] (provided by authors), [2] and [3] as the main data sources, cross validated by [4]. Points in East Asia, Oceania, and North America in the GBIF dataset was removed, as the pathogen does not cause damage on those areas (including its native range) |
| Asian Longhorned Beetle | *Anoplophora glabripennis* | [1] GBIF; [2] Zhou et al. (2021); [3] Zhang et al. (2022); [4] EPPO | North America; Europe (except Iceland); Asia | [1] as the main data source, cross checked by [2]. Pest distribution in China is supplemented by [3] (points in Figure 1). While in dataset [1] there are distribution points in California and Ontario, the pest seems no longer exist in these regions anymore, according to [4]. |
| Beech Bark Disease | *Neonectria faginata* | [1] GBIF; [2] Cale et al. (2017); [3] US Alien Forest Pest Explorer | North America | [1] as the main data source, cross checked by [2] and [3]. The disease specifically cause damage in North America. Therefore, points fall in the Europe in the dataset [2] were deleted |
| Beech Leaf Disease | *Litylenchus crenatae mccannii* | [1] GBIF; [2] Cleveland Metroparks (2023); [3] US Alien Forest Pest Explorer; [4] EPPO | North America | [1], [2], and [3] as main data sources, cross checked by [4]. The disease specifically cause damage in North America and, therefore, data points outside of the North America in the dataset [1] were removed |
| Bronze Birch Borer | *Agrilus anxius* | [1] GBIF; [2] Muilenburg and Herms (2012); [3] EPPO | North America | [1] as the main data source, cross checked by [2] and [3]. It should however be cautious that several states or provinces, such as Alaska, does not have records in database [1], but the pest presents in these areas, according to [2] and [3]. Unfortunately no fine-scale distribution data could be found except from [1]. A point located in the Atlantic in database [1] is invalid and was removed. Points located in Russia and East Asia were removed, as no evidence suggests that the pest causes extensive damage in those regions. |
| Brown Spruce Longhorn Beetle | *Tetropium fuscum* | [1] GBIF | Europe (except Iceland); Russia: Northwestern; Russia: Volga; Russia: Central; Russia: Urals; North America | [1] as the only source. Data points in Mexico and western Canada were deleted, as no evidence suggests that the pest causes damage in those regions |
| Butternut Canker | *Sirococcus clavigignenti-juglandacearum* | [1] Government of Canada (2017); [2] EPPO | North America | [1] as the main data source, cross checked by [2] |
| Dothistroma Needle Blight | *Dothistroma septosporum* | [1] Watt et al. (2009); [2] GBIF; [3] EPPO | None | [1] (points in Figure 2-7) and [2] as the main data sources, cross checked by [3]. Though source [3] reports pathogen presence in some Asian administrative units, no detailed records could be found in [1] and [2]. Also, no significant damage was reported from literature in those regions. Therefore, pathogen presence in those regions in dataset [3] was ignored. |
| Dutch Elm Disease | Ophiostoma ulmi and Ophiostoma novo-ulmi | [1] GBIF; [2] US Alien Forest Pest Explorer | Europe (except Iceland); Russia: Northwestern; Russia: Volga; Russia: Central; North America | European distribution is approximated by source [1]. Distribution in the United States is supplemented by [2]. Distributions in Europe and Canada may not be accurate, but unfortunately no fine-scale data sources could be found |
| Eight-Toothed Spruce Bark Beetle; European Spruce Bark Beetle; Larger Eight-toothed European Spruce Bark Beetle | *Ips typographus* | [1] GBIF; [2] EPPO | Europe (except Iceland), Asia | [1] as the main data source, cross checked by [2]. Data points in North America and Africa in [1] were removed, as the pest only causes significant damage in Europe and East Asia. For the same reason, data records outside of the Europe and East Asia in [2] were ignored |
| Emerald Ash Borer | *Agrilus planipennis* | [1] GBIF; [2] Meshkova et al. (2023); [3] EPPO | Europe (except Iceland); Russia: Northwestern; Russia: Volga; Russia: Central; North America | [1] as the main data source, cross checked by [2] (Table S1) and [3]. Data outside of North America and Europe was removed, as the pest cause limited damage in its native range, i.e. East Asia |
| Fall Webworm; American White Moth | *Hyphantria cunea* | [1] Ning et al. (2022); [2] GBIF; [3] EPPO | Europe (except Iceland); Asia | [1] and [2] as the main data source, cross checked by [3]. The pest causes much lower level of damage in its native range - North America. Thus, data points in its native range were removed |
| Forest Tent Caterpillar | *Malacosoma disstria* | [1] GBIF; [2] EPPO | North America | [1] as the main data source, cross checked by [2]. Removed an Indian data point in the dataset [1], as the pest does not appear to present in Eurasia according to [2] |
| Gypsy Moth; Spongy Moth | *Lymantria dispar* | [1] GBIF; [2] Song et al. (2022); [3] EPPO | Europe (except Iceland); Asia; North America | [1] as the main data sources, cross checked by [2] and [3] |
| Hemlock Woolly Adelgid | *Adelges tsugae* | [1] GBIF; [2] US Alien Forest Pest Explorer | North America | [1] as the main data source, cross checked by [2]. Removed points in Eurasia in dataset [2], as the pest does not appear to cause significant damage in its native area |
| Japanese Oak Wilt | *Raffaelea quercivora* | [1] Kuroda et al. (2010); [2] EPPO | Japan | [1] as the main data source, cross checked by [2] |
| Laurel Wilt | *Raffaelea lauricola* | [1] US Alien Forest Pest Explorer; [2] EPPO | North America | [1] as the main data source, cross checked by [2]. Although the pathogen is native to East Asia, it causes damage primarily in the United States |
| Mountain Pine Beetle | *Dendroctonus ponderosae* | [1] GBIF; [2] Logan and Powell (2001); [3] EPPO | North America | [1] as the main data source, cross checked by [2] and [3]. Removed a data point in Europe in the database [1], as the pest does not present in Europe |
| Oak Decline | *Brenneria goodwinii; Gibbsiella quercinecans* | [1] Kowsari and Karimi (2023); [2] EPPO | North America; Europe (except Iceland); Asia | [1] as the main data source, cross checked by [2]. The disease caused damage in North America, Europe, and Asia. |
| Oak Wilt | *Bretziella fagacearum* | [1] Pedlar et al. (2020); [2] EPPO | North America | [1] (Figure 1a) as the main data source, cross checked by [2] |
| Oriental Chestnut Gall Wasp; Chestnut Gall Wasp | *Dryocosmus kuriphilus* | [1] GBIF; [2] EPPO | Europe (except Iceland); Asia, North America | [1] as the main data source, cross checked by [2]. |
| Pine Processionary Moth; Pine Caterpillar; Pine processionary Moth Caterpillar | *Thaumetopea pityocampa* | [1] GBIF; [2] Kerdelhué et al. (2009); [3] EPPO | Europe (except Iceland); Russia: Northwestern; Russia: Volga; Russia: Central | [1] as the main data source, cross checked by [2] and [3] |
| Pine Wood Nematode; Pine Wilt Disease | *Bursaphelenchus xylophilus* | [1] Aierken et al. (2024); [2] EPPO | Europe (except Iceland); Asia | [1] (Figure 1) as the main data source, cross checked by [2] |
| Red Turpentine Beetle | *Dendroctonus valens* | [1] Zhou et al. (2024); [2] EPPO | China | [1] as the main data source. Since the pest does not cause great damage in its native range, North America, those data points were removed |
| Red-necked Longhorn Beetle | *Aromia bungii* | [1] GBIF; [2] Horrocks et al. (2024) | Russian: Far Eastern; China; Japan; North Korea; South Korea | [1] and [2] as the data sources |
| Spotted Lanternfly | *Lycorma delicatula* | [1] GBIF; [2] EPPO; [3] Wakie et al. (2020) | Russia: Far Eastern; China; Japan; North Korea; South Korea; North America | [1] as the main data source, cross checked by [2] and [3] |
| Spruce Budworm; Eastern Spruce Budworm | *Choristoneura fumiferana* | [1] GBIF; [2] Nealis (2015); [3] EPPO | North America | [1] as the main data source, cross checked by [2] and [3] |
| Sudden Oak Death; Ramorum Disease; Ramorum Blight, Ramorum Dieback | *Phytophthora ramorum* | [1] Ireland et al. (2013); [2] EPPO; [3] US Alien Forest Pest Explorer | Europe (except Iceland and Russia); North America | [1] as the main data source (point data in Figure 2 for the United States and Norway), cross checked by [2] and [3]. Since the pathogen the Europe, but no European distribution data points (except Norway) were available, projection should be careful |
| Sweet Chestnut Blight; Chestnut Blight | *Cryphonectria parasitica* | [1] GBIF; [2] EPPO | North America; Europe (except Iceland and Russia) | [1] as the main data source, cross checked by [2]. Distribution points in East Asia were removed, as the pathogen causes relatively low damage in its native range |
| White Pine Blister Rust | *Cronartium ribicola* | [1] GBIF; [2] EPPO | North America; Europe (except Iceland); Russia: Northwestern; Russia: Volga; Russia: Central | [1] as the main data source, cross checked by [2]. Data points outside of Europe and North America in the dataset [1] was removed, as the pathogen causes significantly lower level of damage in East Asia, where host trees show partial resistance |
| Xylella | *Xylella fastidiosa* | [1] Koufakis et al. (2016); [2] EPPO | North America; South America; Europe (except Iceland); Iraq; Iran; Lebanon; Israel; Taiwan, China | [1] as the main data source, cross checked by [2] |

**Table S2. WorldClim Bioclimatic Variables\**

**Supplementary Table S3.** Explanation of 19 WorldClim bioclimatic variables.

| **Variable name** | **Meaning** |
| --- | --- |
| bio_1 | Annual Mean Temperature |
| bio_2 | Mean Diurnal Range (Mean of monthly (max temp - min temp)) |
| bio_3 | Isothermality (bio_2/bio_7) (×100) |
| bio_4 | Temperature Seasonality (standard deviation ×100) |
| bio_5 | Max Temperature of Warmest Month |
| bio_6 | Min Temperature of Coldest Month |
| bio_7 | Temperature Annual Range (bio_5-bio_6) |
| bio_8 | Mean Temperature of Wettest Quarter |
| bio_9 | Mean Temperature of Driest Quarter |
| bio_10 | Mean Temperature of Warmest Quarter |
| bio_11 | Mean Temperature of Coldest Quarter |
| bio_12 | Annual Precipitation |
| bio_13 | Precipitation of Wettest Month |
| bio_14 | Precipitation of Driest Month |
| bio_15 | Precipitation Seasonality (Coefficient of Variation) |
| bio_16 | Precipitation of Wettest Quarter |
| bio_17 | Precipitation of Driest Quarter |
| bio_18 | Precipitation of Warmest Quarter |
| bio_19 | Precipitation of Coldest Quarter |

**Table S3. AUC**

**Supplementary Table S3.** Area Under the Curve (AUC) values for training and testing data of MaxEnt models predicting the distribution of tree-infecting pests and pathogens. AUC (Area Under the Curve) values quantify the performance of MaxEnt models, with values ranging from 0 to 1 (higher values indicate better ability of the model to discriminate true presence). For each MaxEnt model, species presence data were split into two subsets: 75% of the data were used to train and build the model (training data), and the remaining 25% were reserved to evaluate model performance (testing data). Training data AUC reflects model performance on the data used to build the model, while testing data AUC indicates performance on independent data not used in model training. Values are provided for each pest or pathogen evaluated in the study.

| **Pest/pathogen** | **AUC on training data** | **AUC on testing data** |
| --- | --- | --- |
| Ash Dieback | 0.8552 | 0.8621 |
| Asian Longhorned Beetle | 0.8715 | 0.8459 |
| Beech Bark Disease | 0.9479 | 0.9501 |
| Beech Leaf Disease | 0.9375 | 0.9441 |
| Bronze Birch Borer | 0.8495 | 0.8459 |
| Brown Spruce Longhorn Beetle | 0.8391 | 0.8348 |
| Butternut Canker | 0.8275 | 0.8289 |
| Dothistroma Needle Blight | 0.7975 | 0.7887 |
| Dutch Elm Disease | 0.6643 | 0.662 |
| Eight-Toothed Spruce Bark Beetle | 0.7726 | 0.7784 |
| Emerald Ash Borer | 0.8274 | 0.7784 |
| Fall Webworm | 0.845 | 0.8547 |
| Forest Tent Caterpillar | 0.7453 | 0.7343 |
| Gypsy Moth; Spongy Moth | 0.6206 | 0.6117 |
| Hemlock Woolly Adelgid | 0.8868 | 0.8817 |
| Japanese Oak Wilt | 0.9897 | 0.9904 |
| Laurel Wilt | 0.9552 | 0.955 |
| Mountain Pine Beetle | 0.8277 | 0.8394 |
| Oak Decline | 0.6787 | 0.6436 |
| Oak Wilt | 0.8861 | 0.8764 |
| Oriental Chestnut Gall Wasp | 0.7997 | 0.8128 |
| Pine Processionary Moth | 0.7871 | 0.7947 |
| Pine Wood Nematode | 0.8267 | 0.8334 |
| Red Turpentine Beetle | 0.9754 | 0.9717 |
| Red-necked Longhorn Beetle | 0.9916 | 0.9903 |
| Spotted Lanternfly | 0.8822 | 0.8687 |
| Spruce Budworm; Eastern Spruce Budworm | 0.8703 | 0.8715 |
| Sudden Oak Death | 0.9283 | 0.9202 |
| Sweet Chestnut Blight | 0.8589 | 0.8658 |
| White Pine Blister Rust | 0.7771 | 0.7793 |
| Xylella | 0.861 | 0.8536 |

**Figure S1. Individual Pest or Pathogen Range**

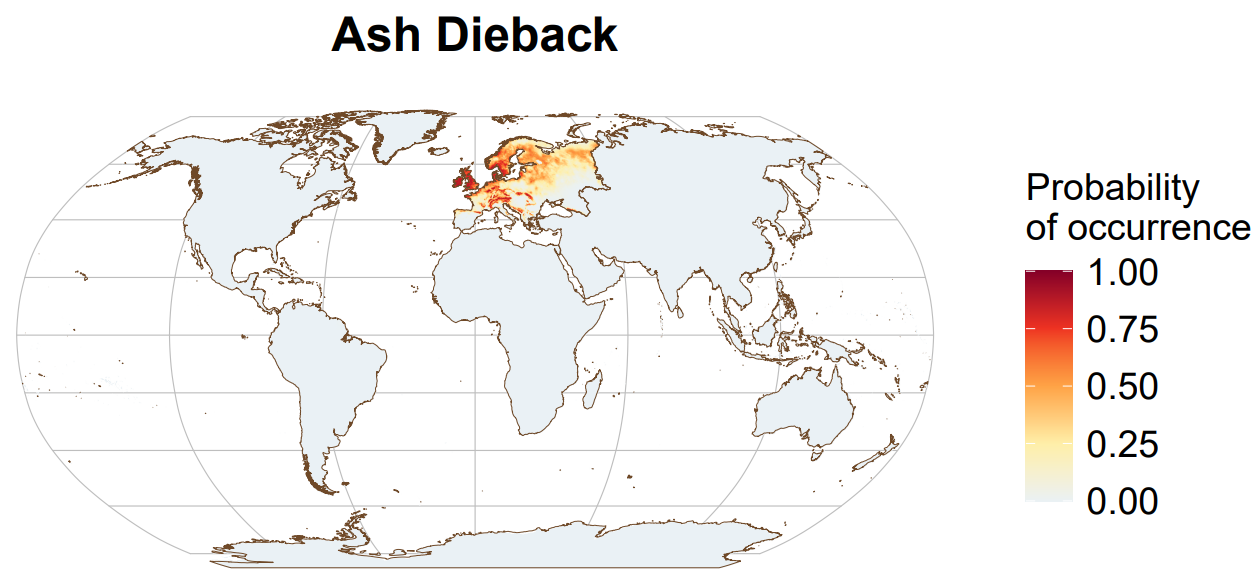

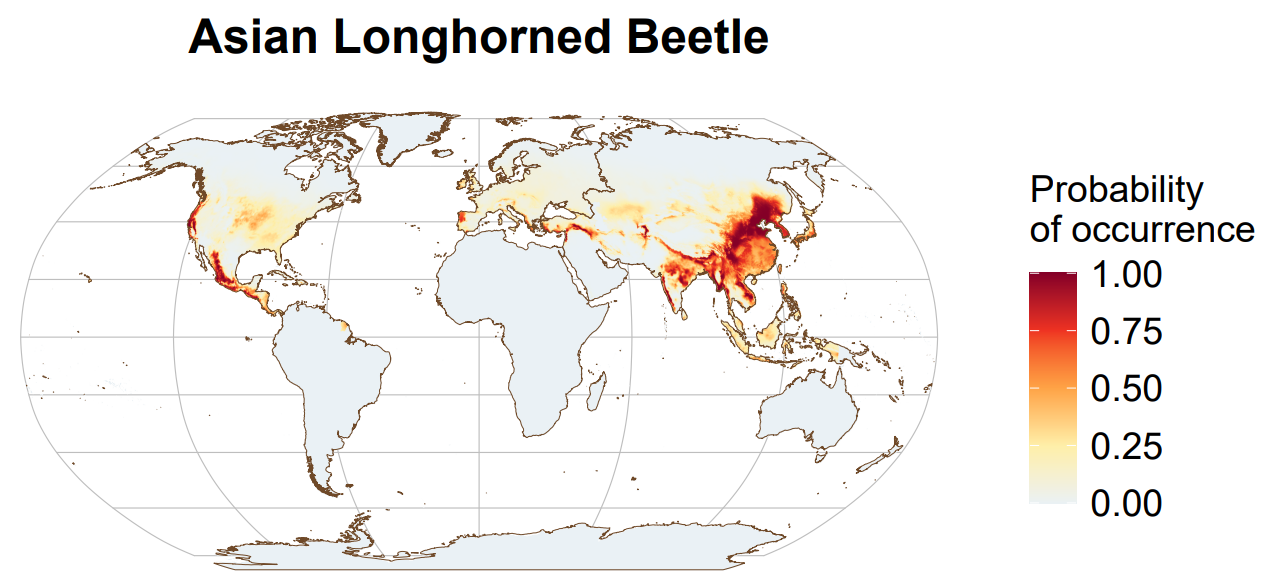

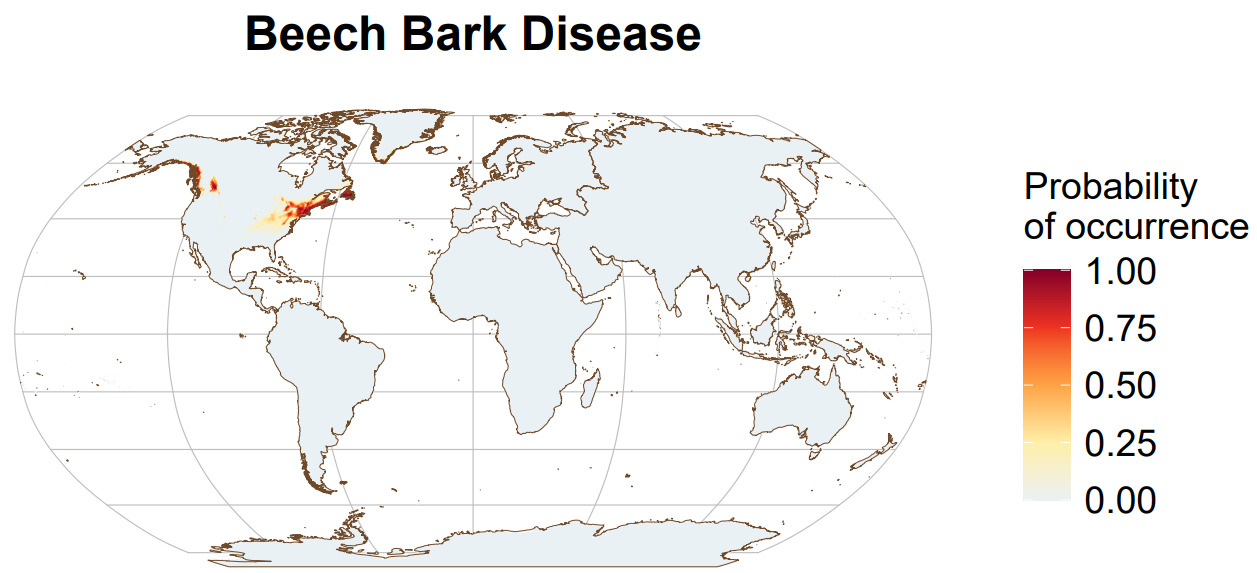

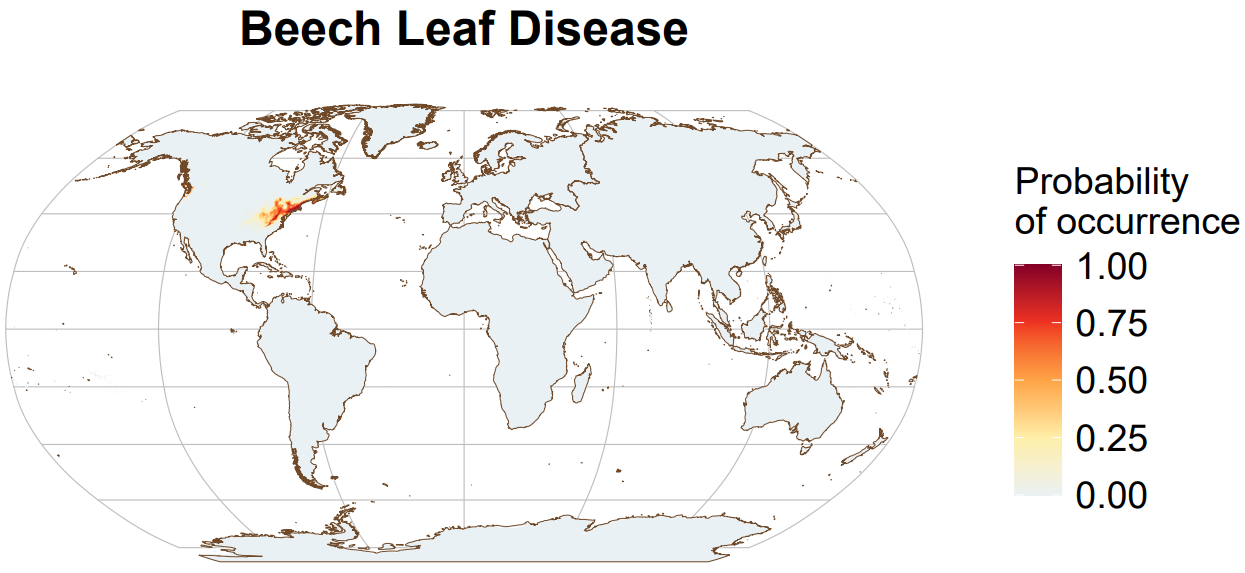

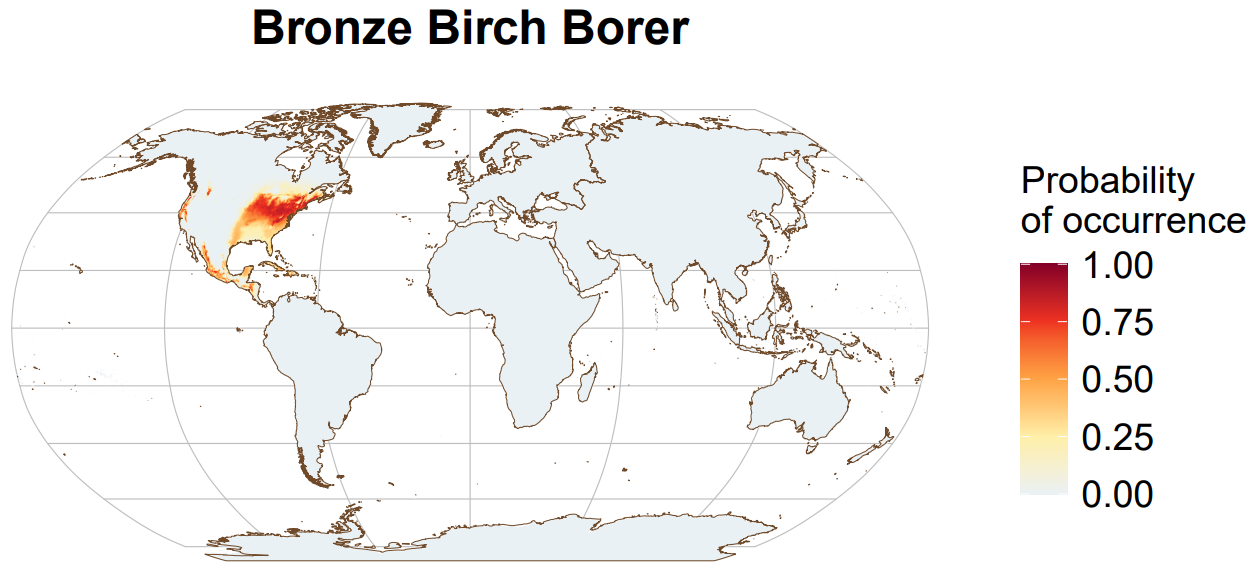

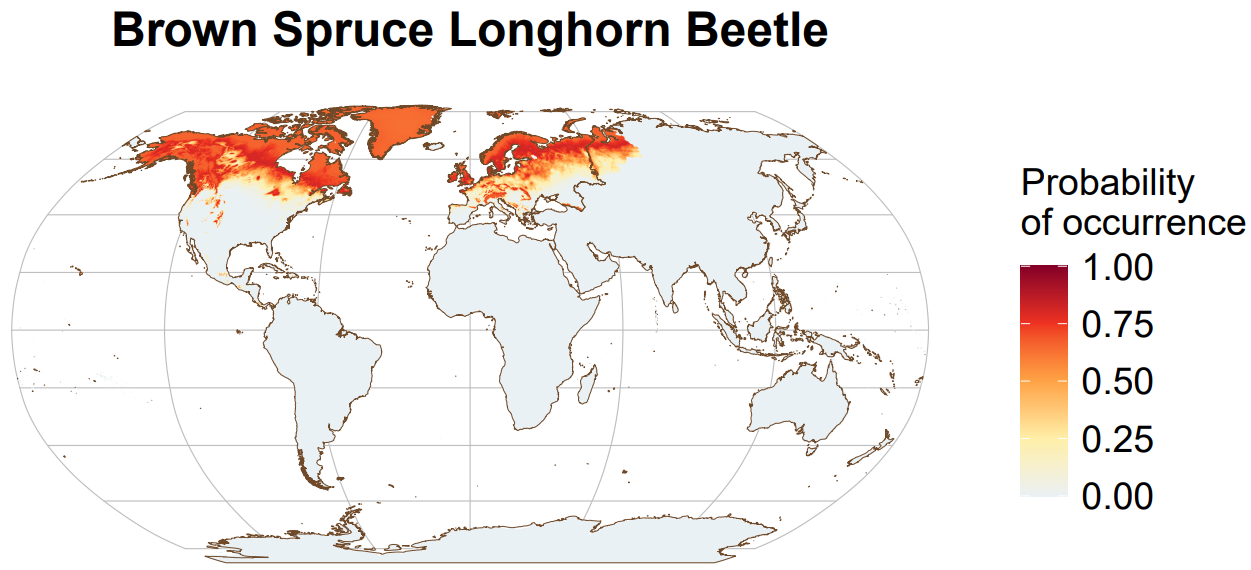

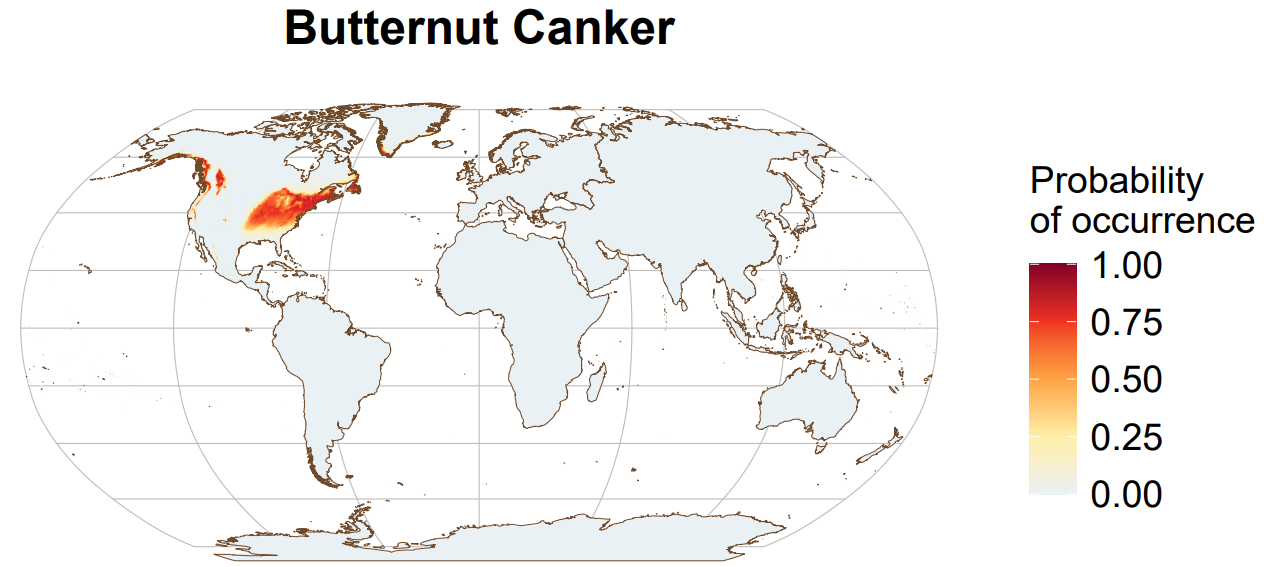

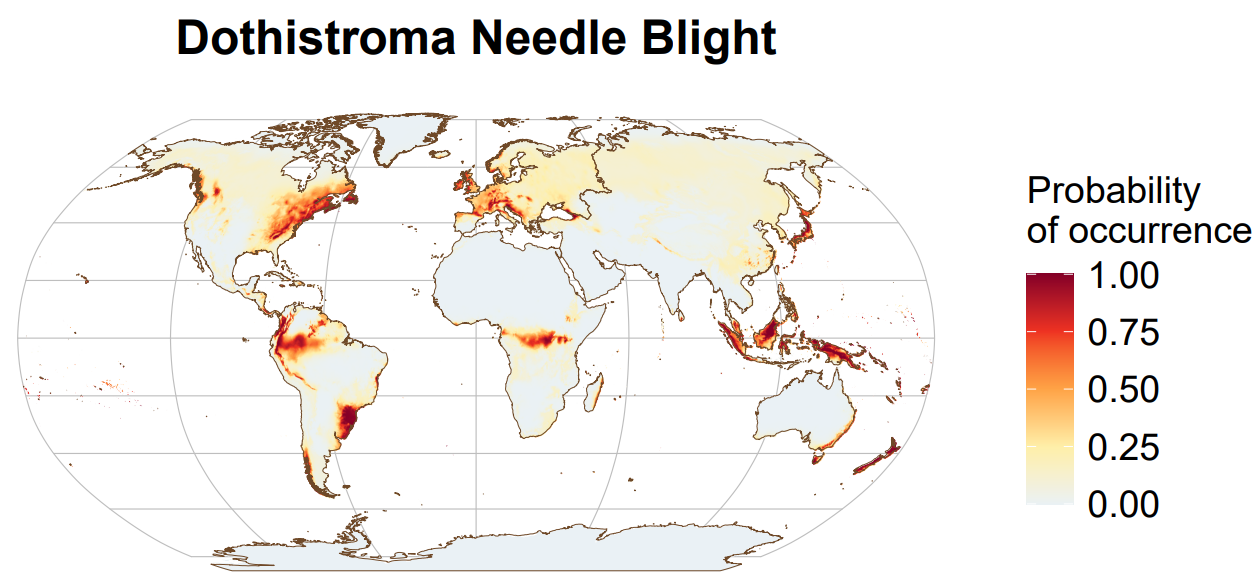

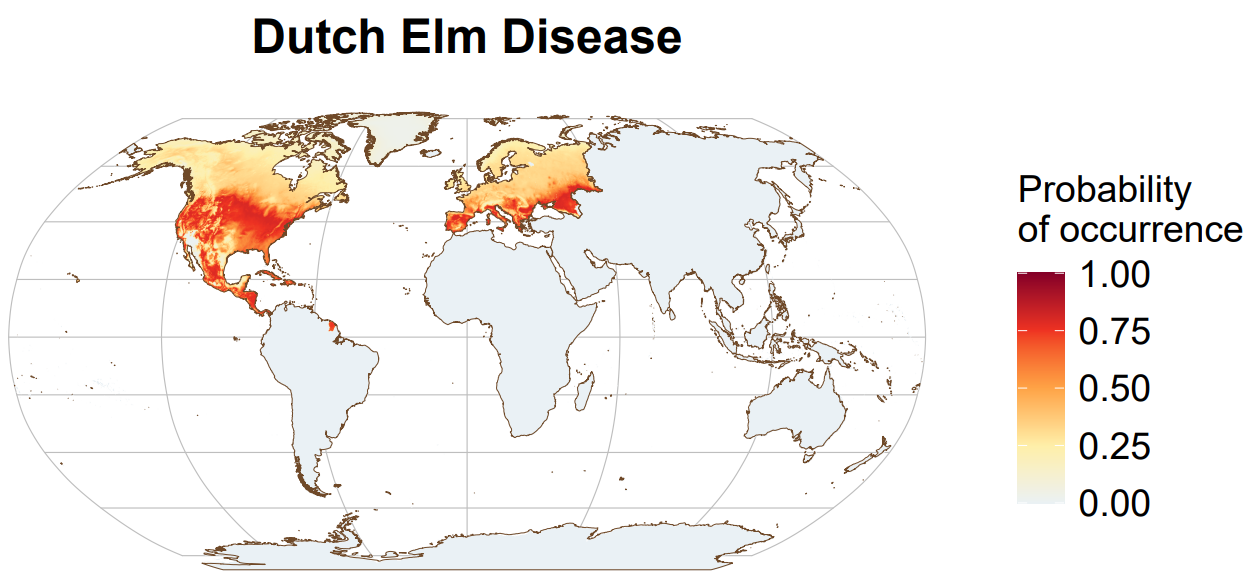

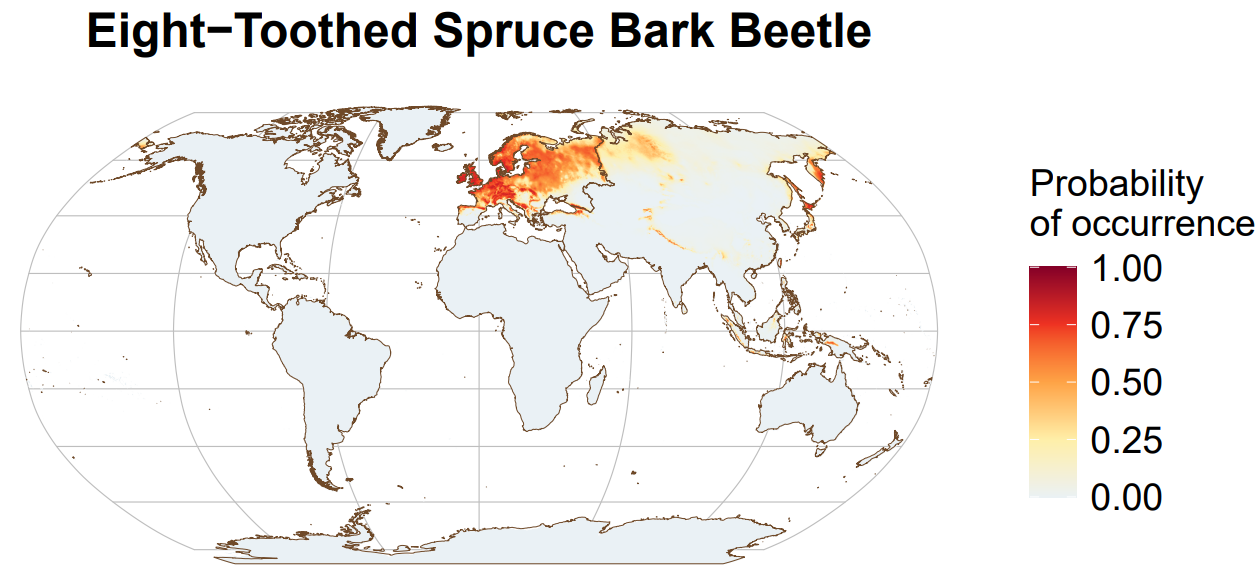

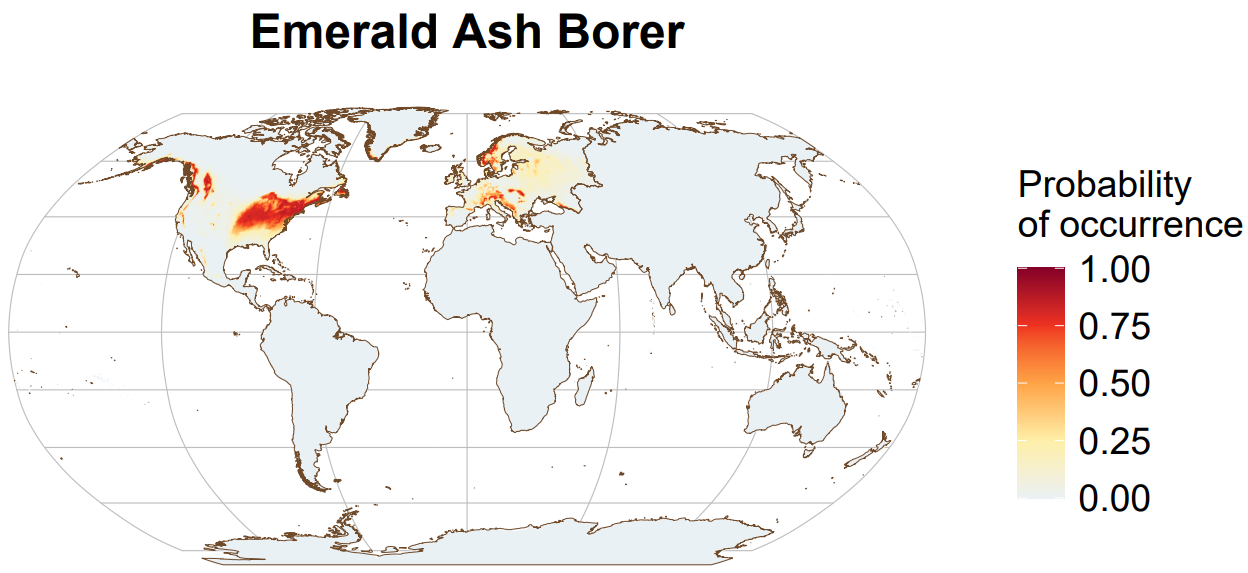

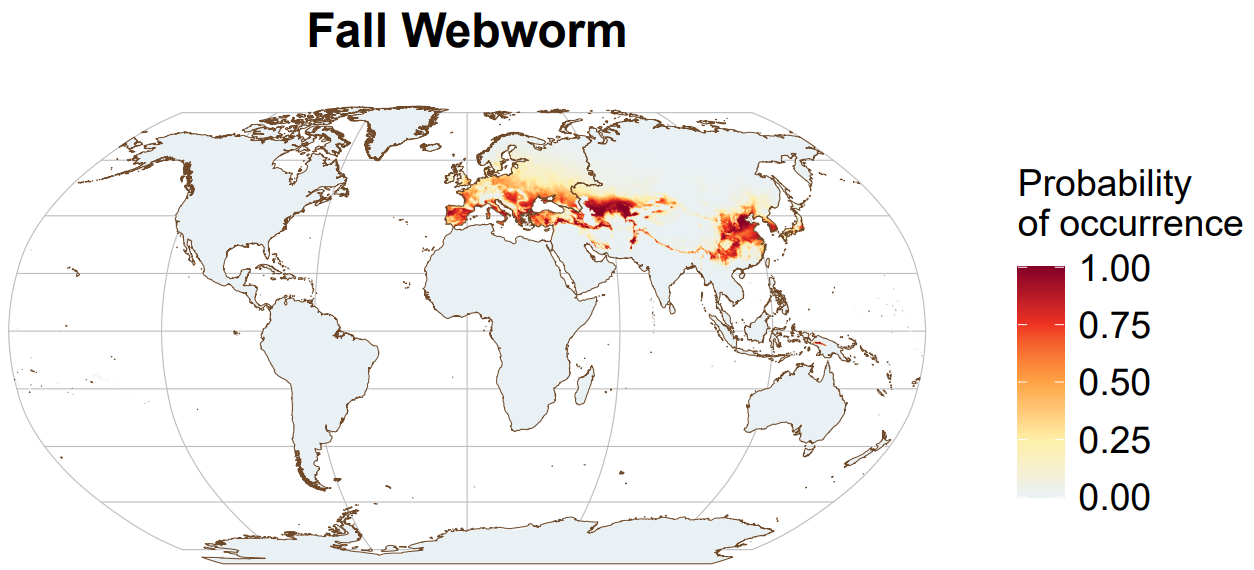

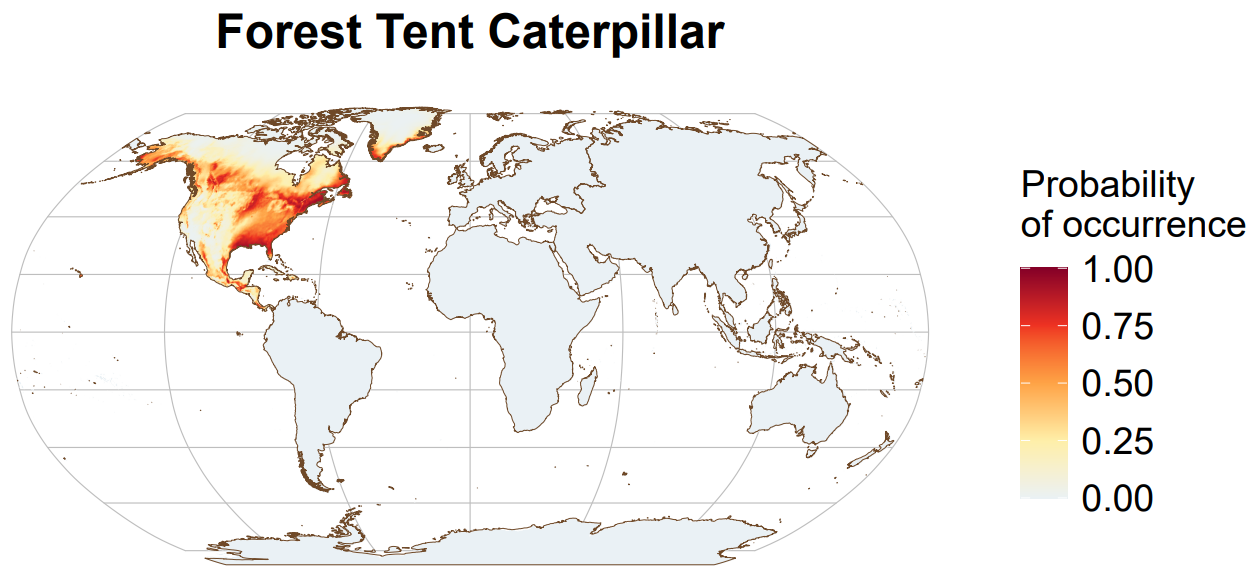

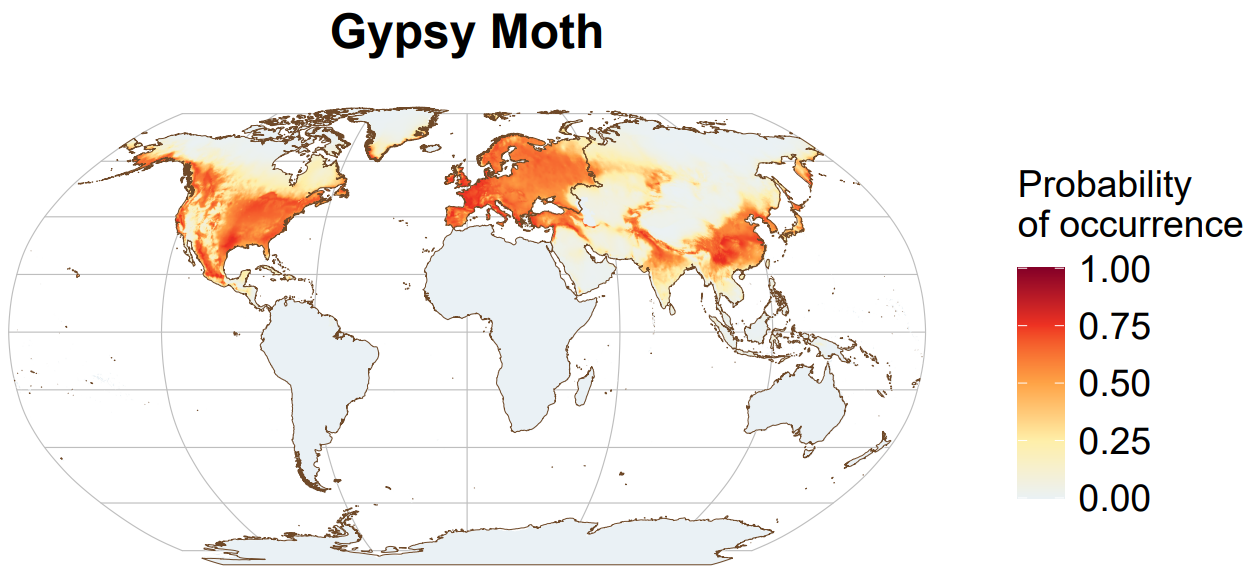

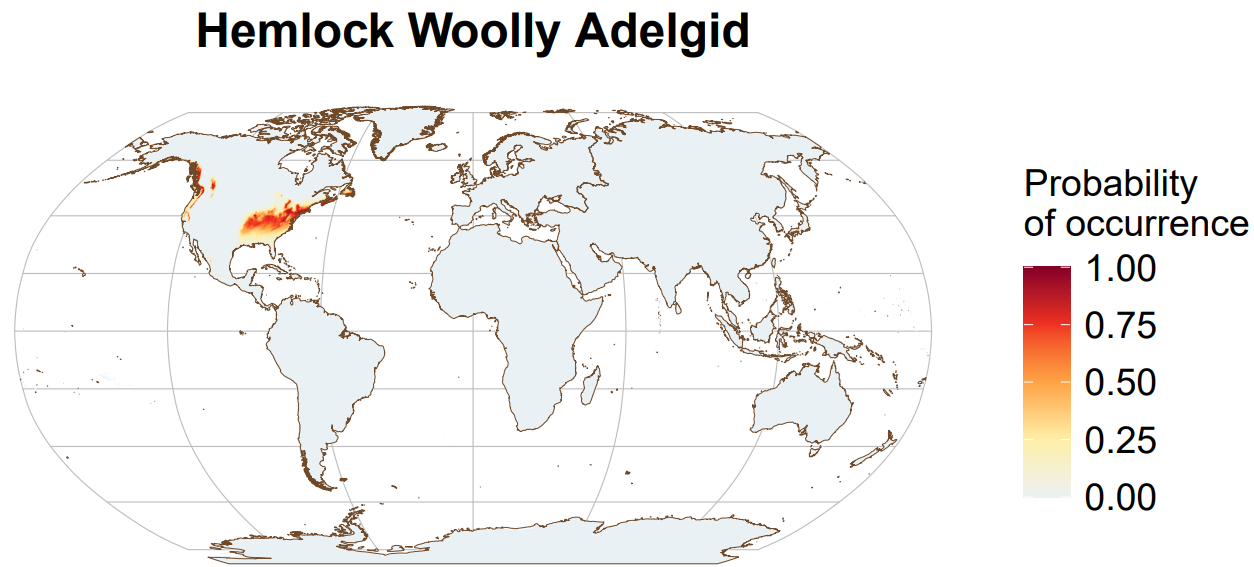

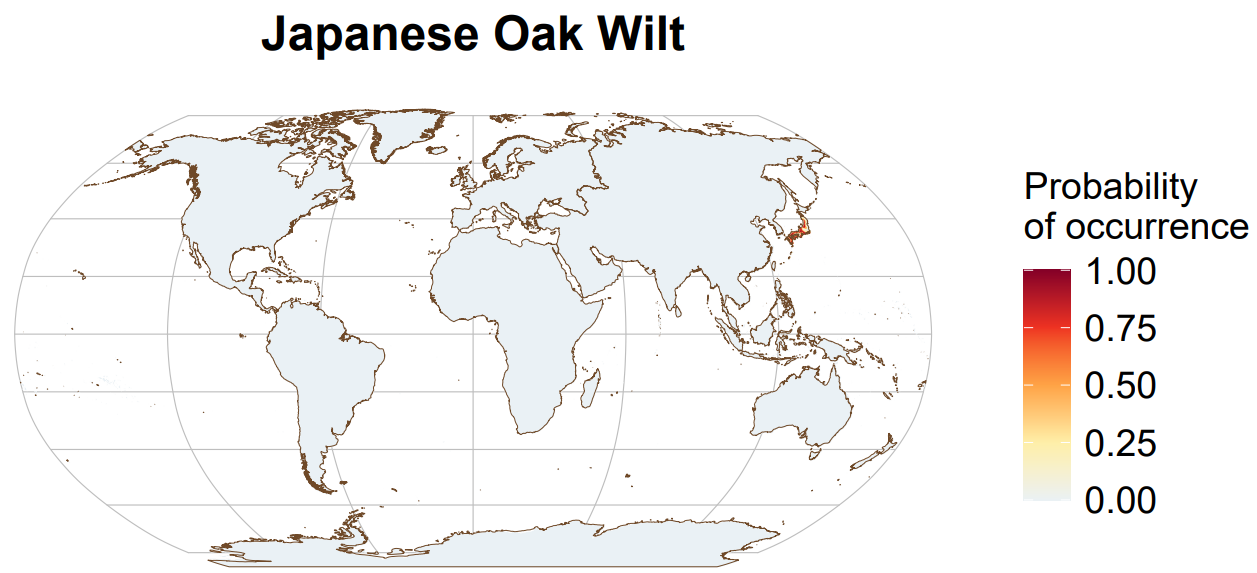

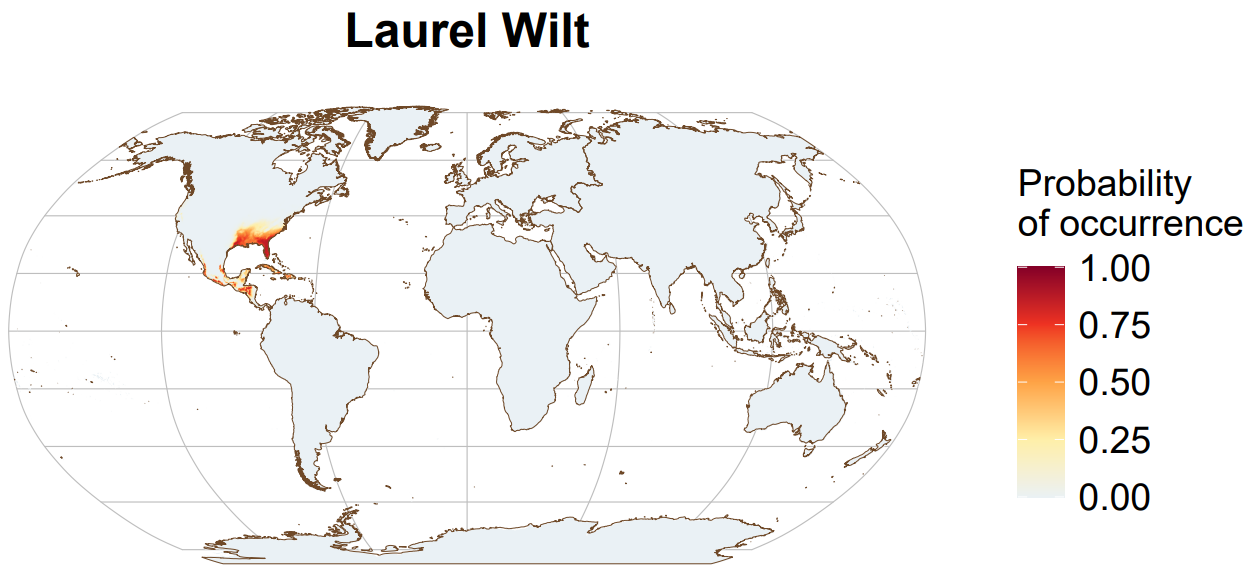

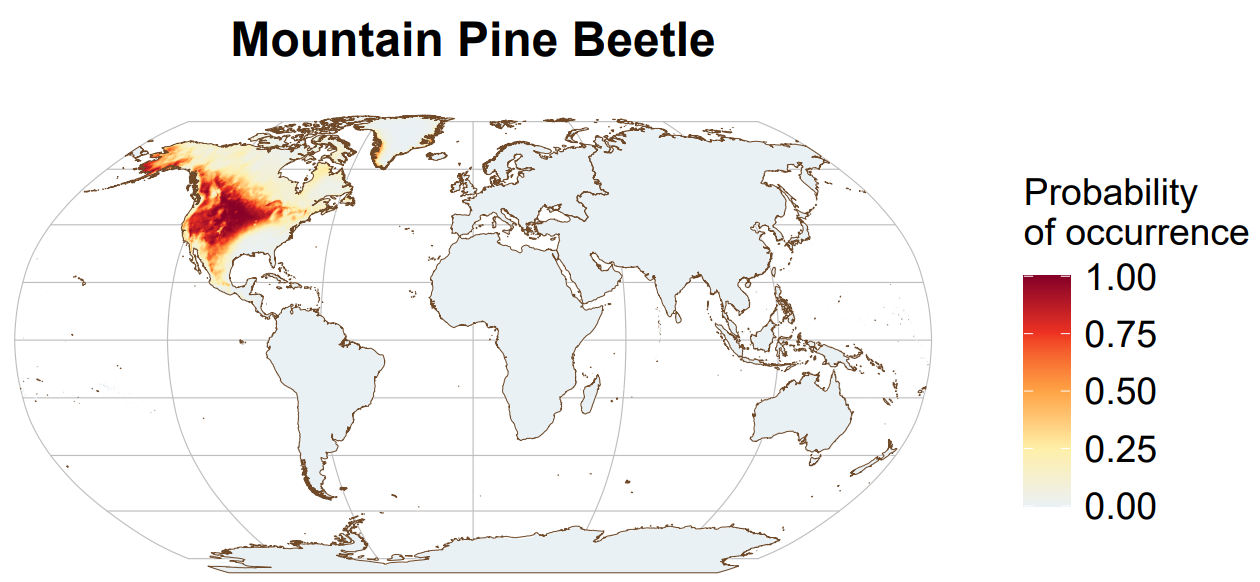

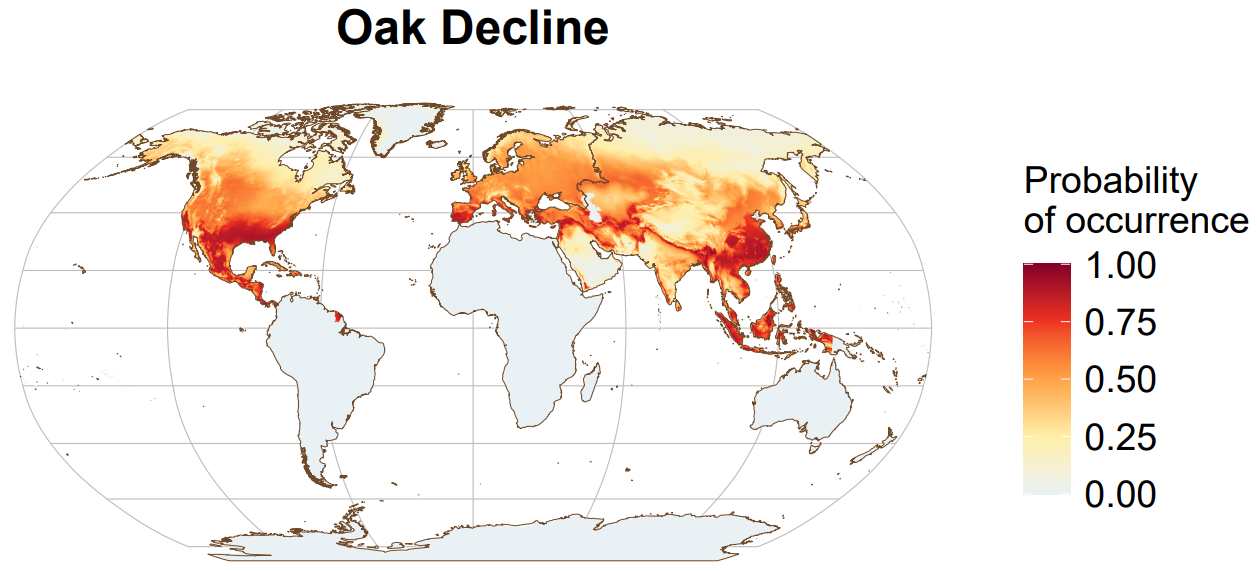

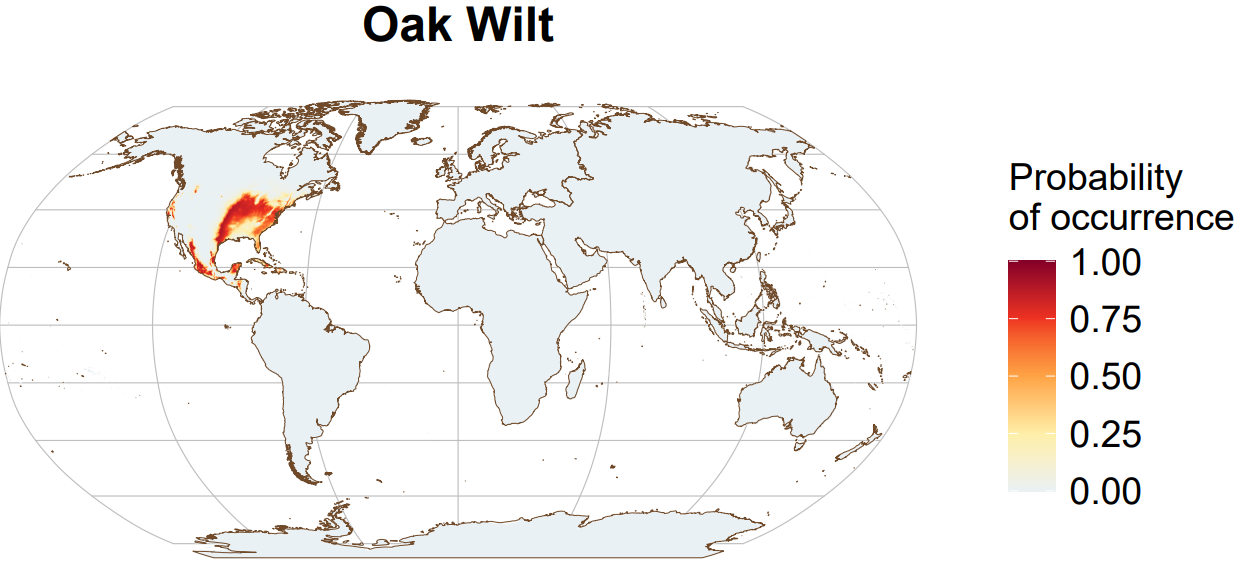

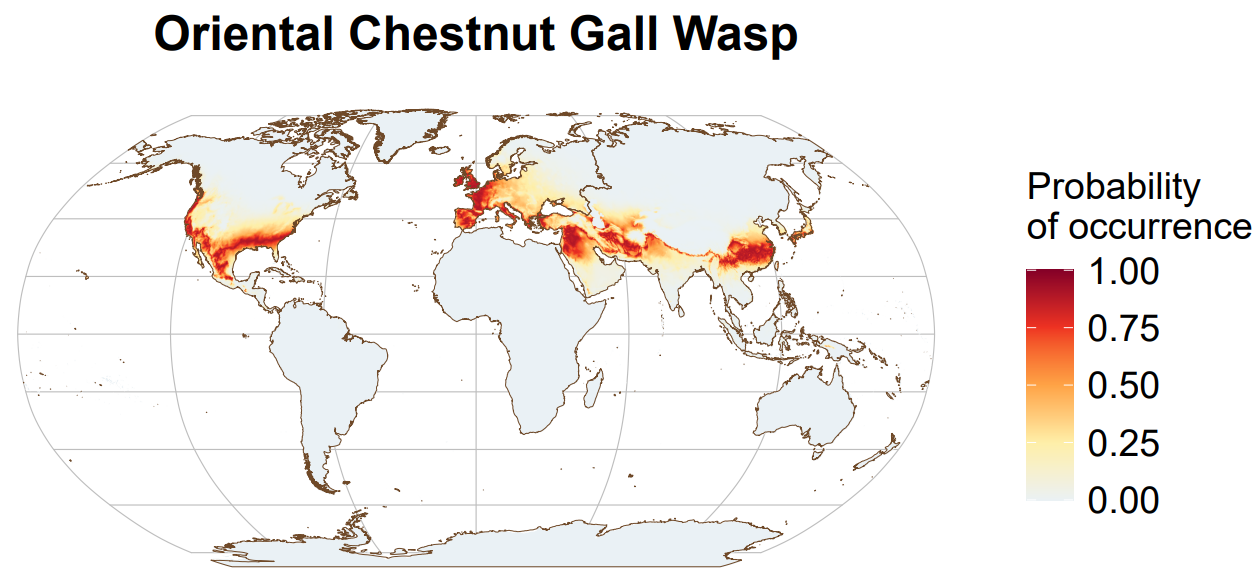

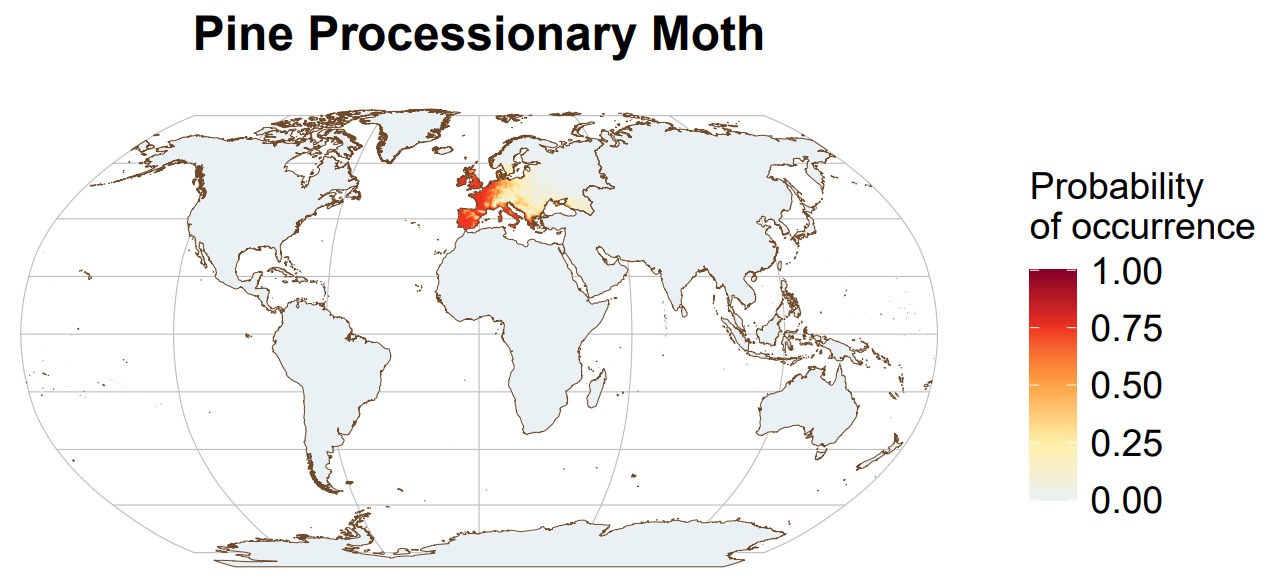

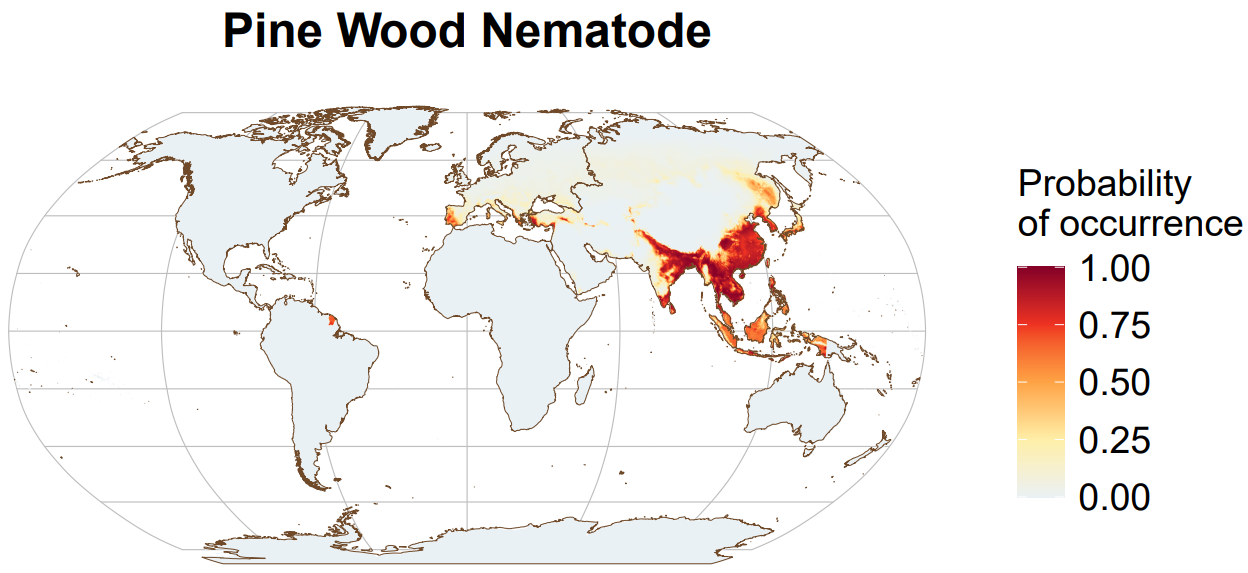

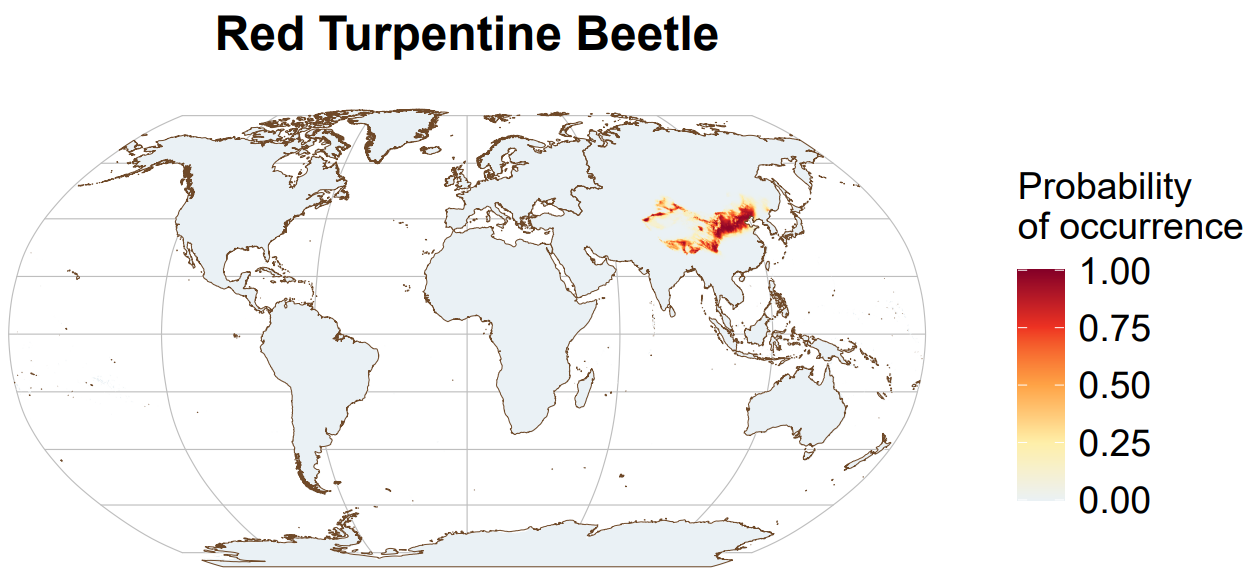

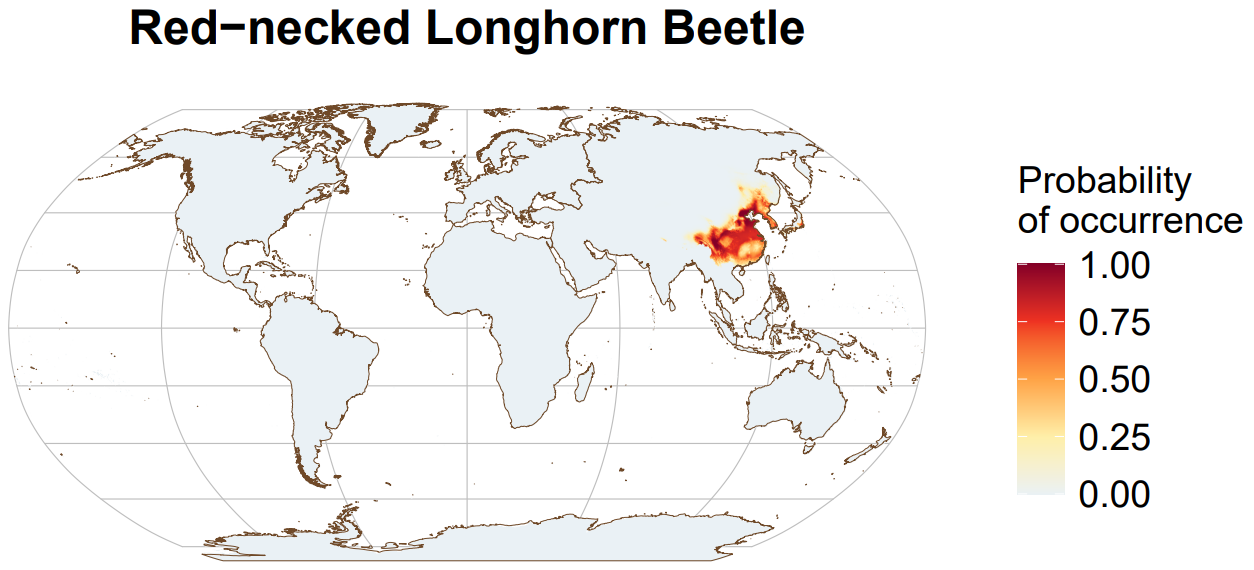

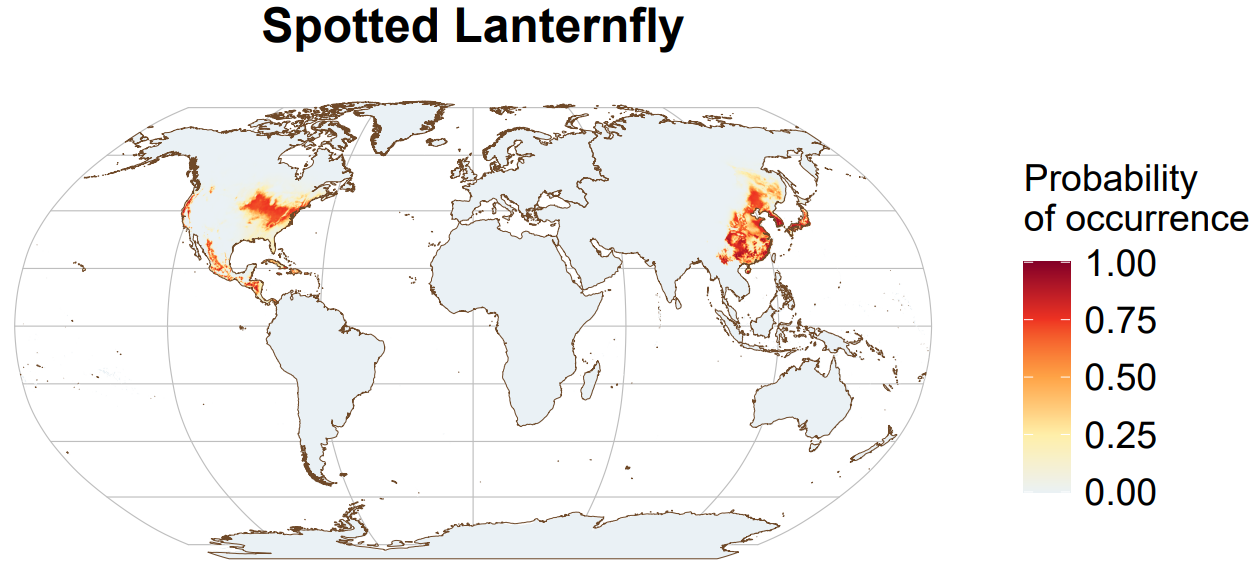

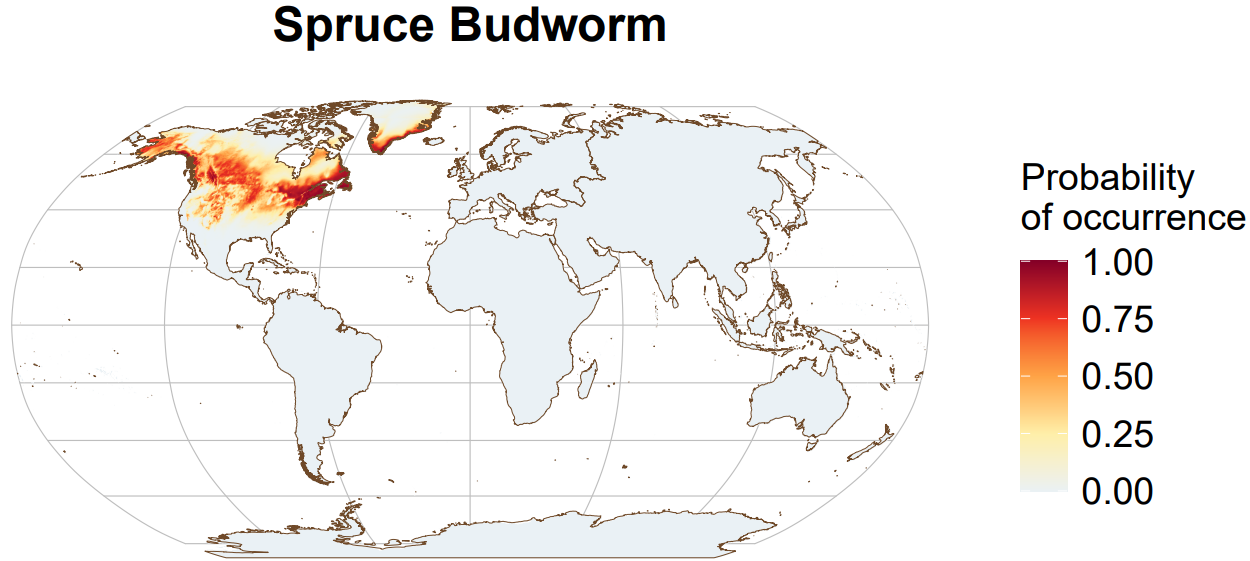

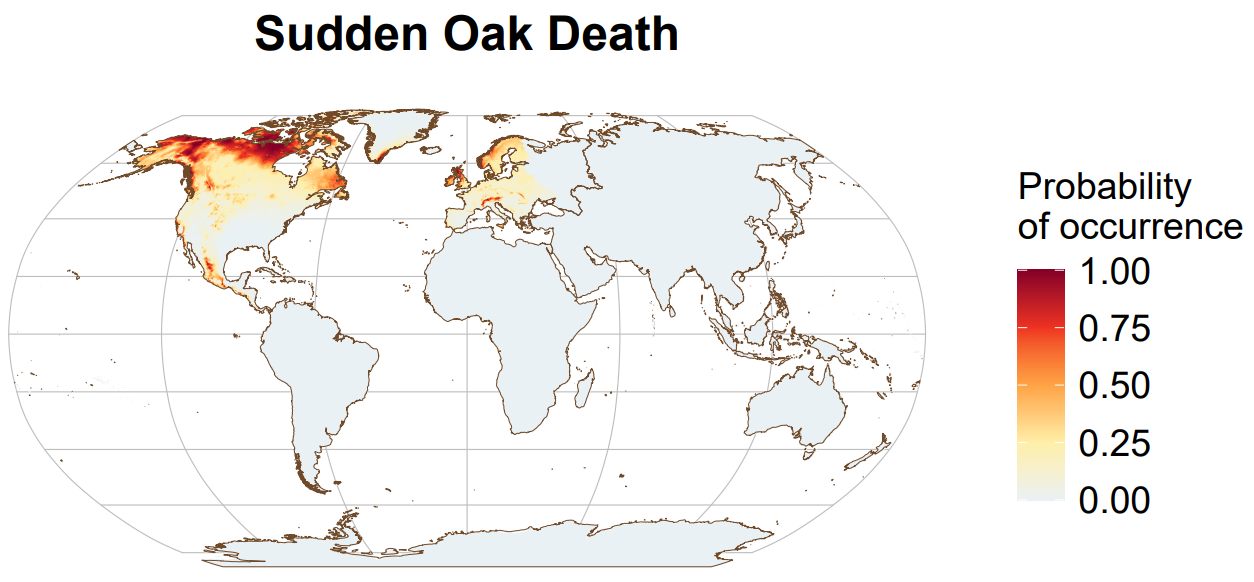

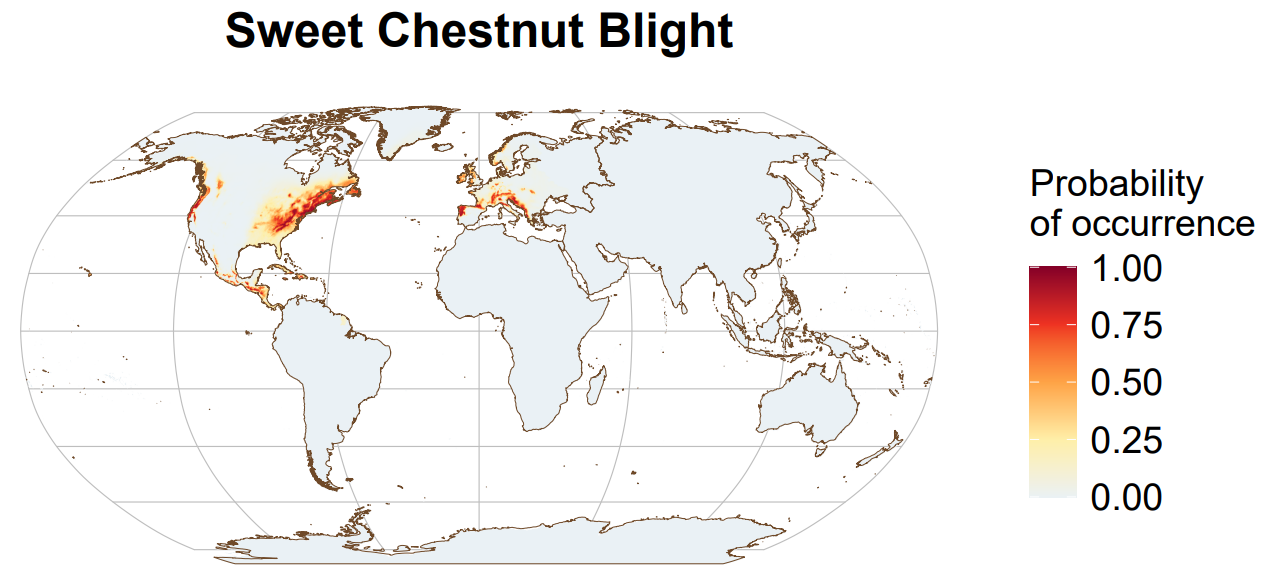

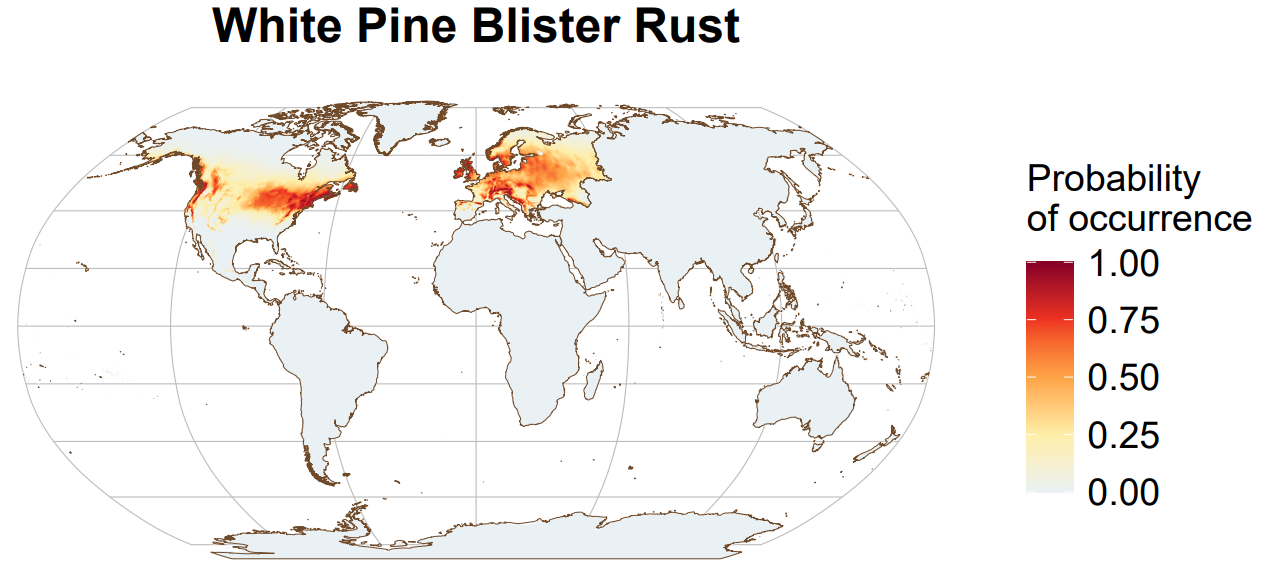

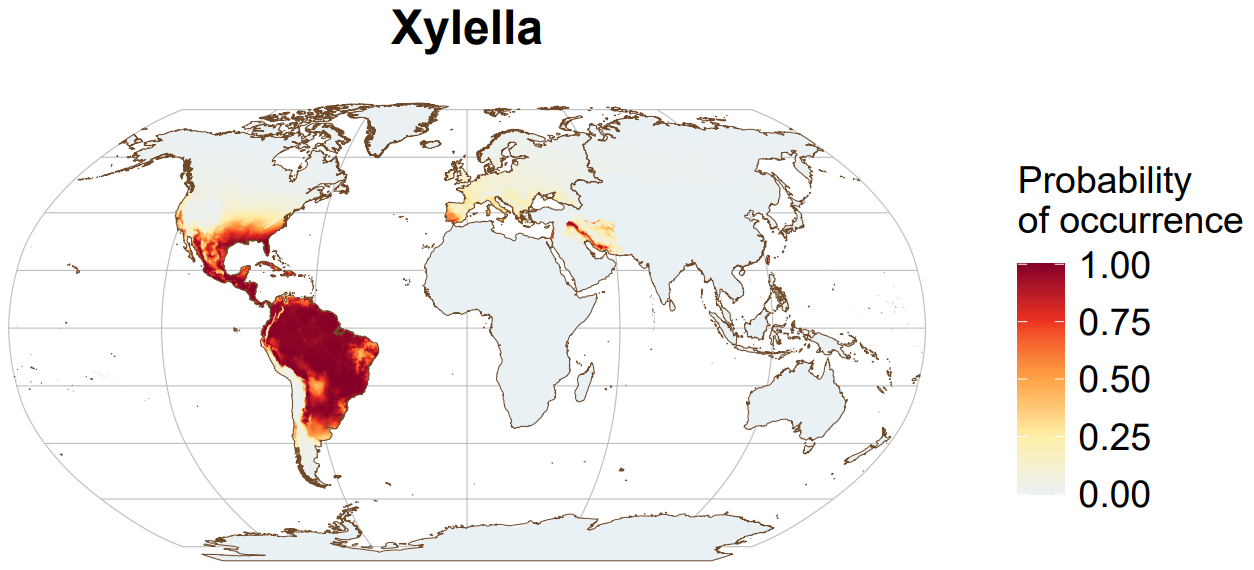

**Supplementary Figure S2.** Modelled spatial distribution for 31 major forest pests and pathogens under current conditions.

### Figure S2. Standard Deviation of Future Pest or Pathogen Richness

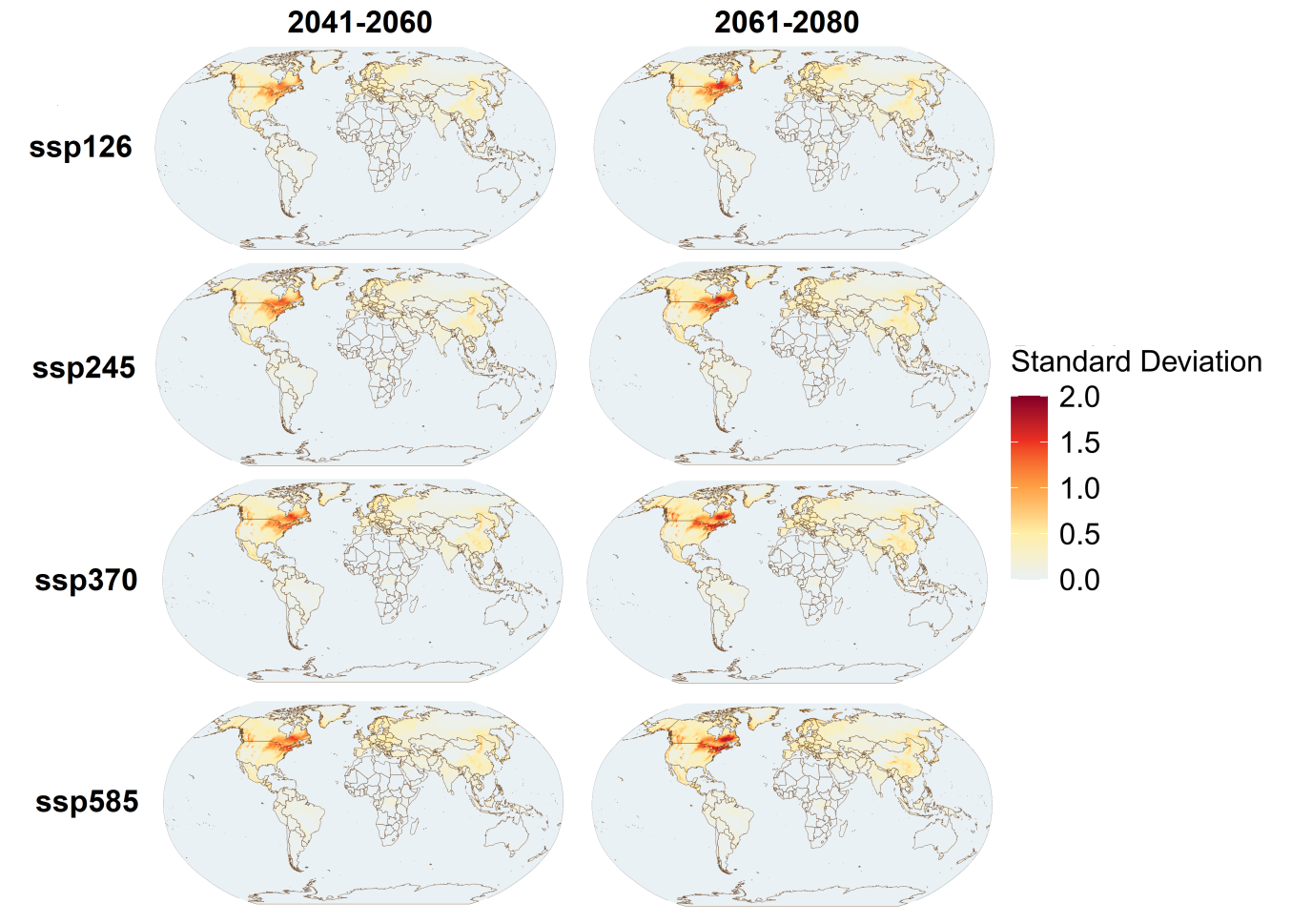

**Supplementary Figure S2.** Standard deviation of model projections of tree pest and pathogen richness across 11 global climate models (CGMs): ACCESS-CM2, BCC-CSM2-MR, CMCC-ESM2, EC-Earth3-Veg, GISS-E2-1-G, INM-CM5-0, IPSL-CM6A-LR, MIROC6, MPI-ESM1-2-HR, MRI-ESM2-0, UKESM1-0-LL. Projections are made under four Shared Socio-economic Pathways (SSP126, SSP245, SSP370, SSP585) for two future periods (2041–2060 and 2061–2080).

**Figure S3. Future Pest and Pathogen Richness**

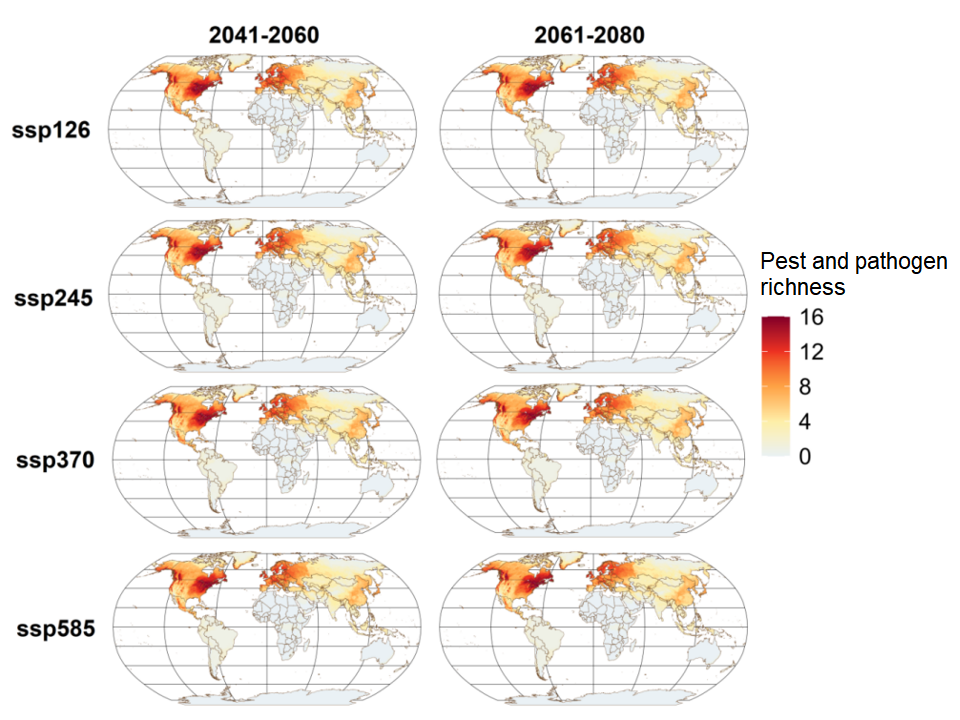

**Supplementary Figure S3.** Projections of tree pest and pathogen richness under four Shared Socio-economic Pathways (SSP126, SSP245, SSP370, SSP585) for two future periods (2041–2060 and 2061–2080). Color intensity represents pest and pathogen richness.

### Figure S4. Change of Future Pest or Pathogen Range

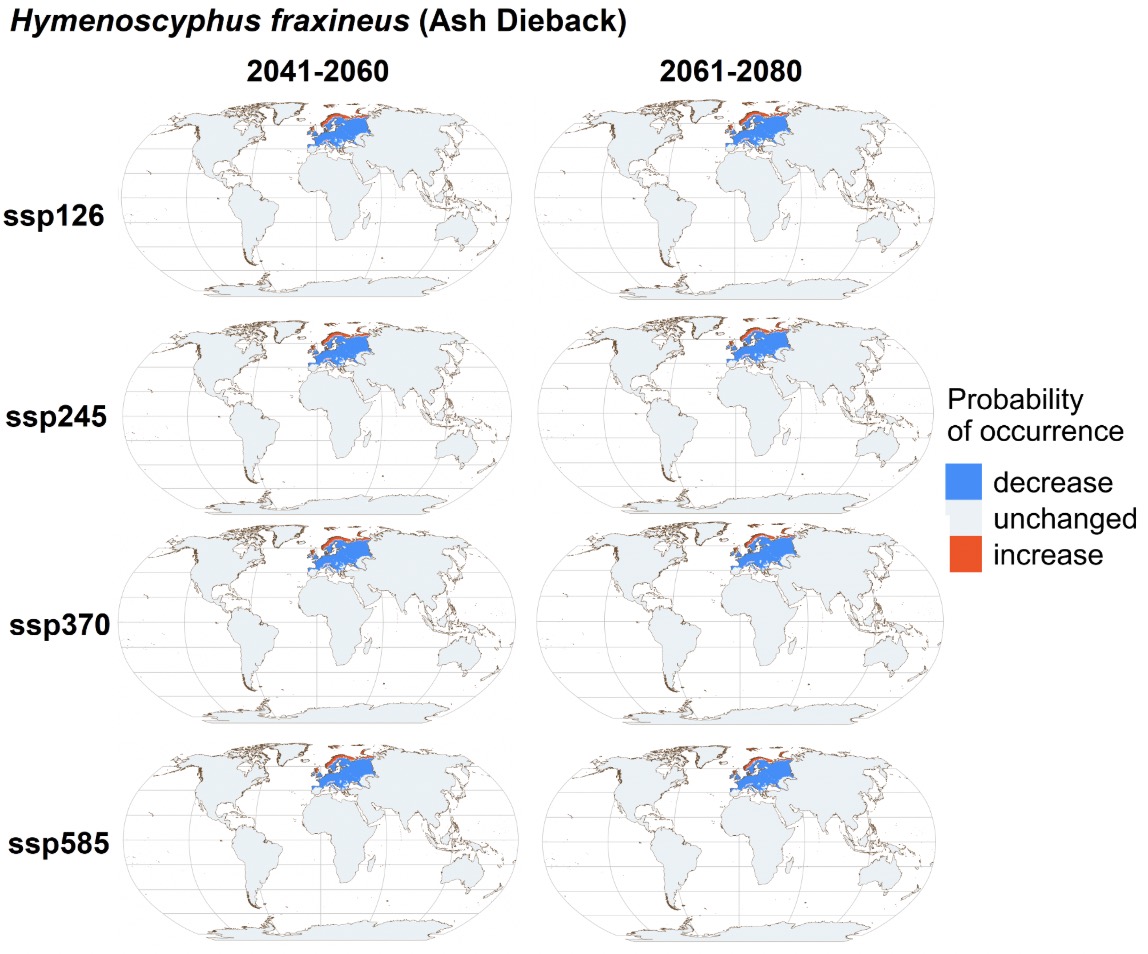

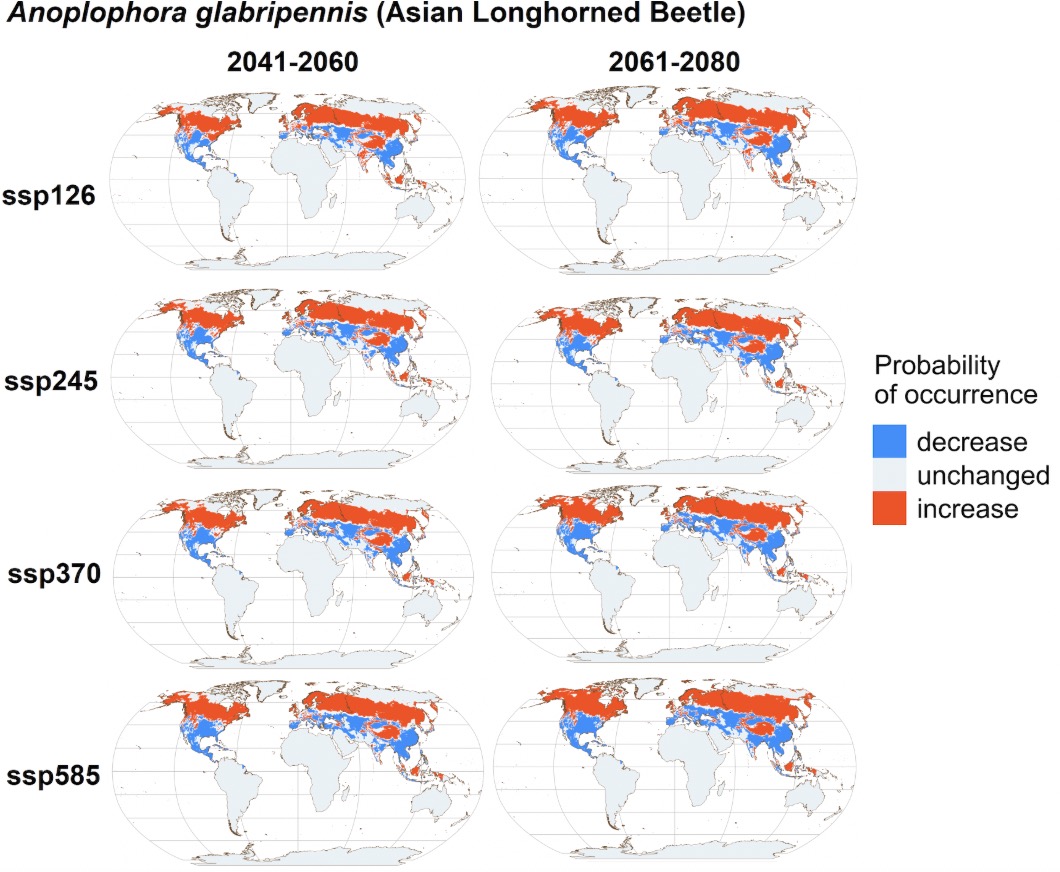

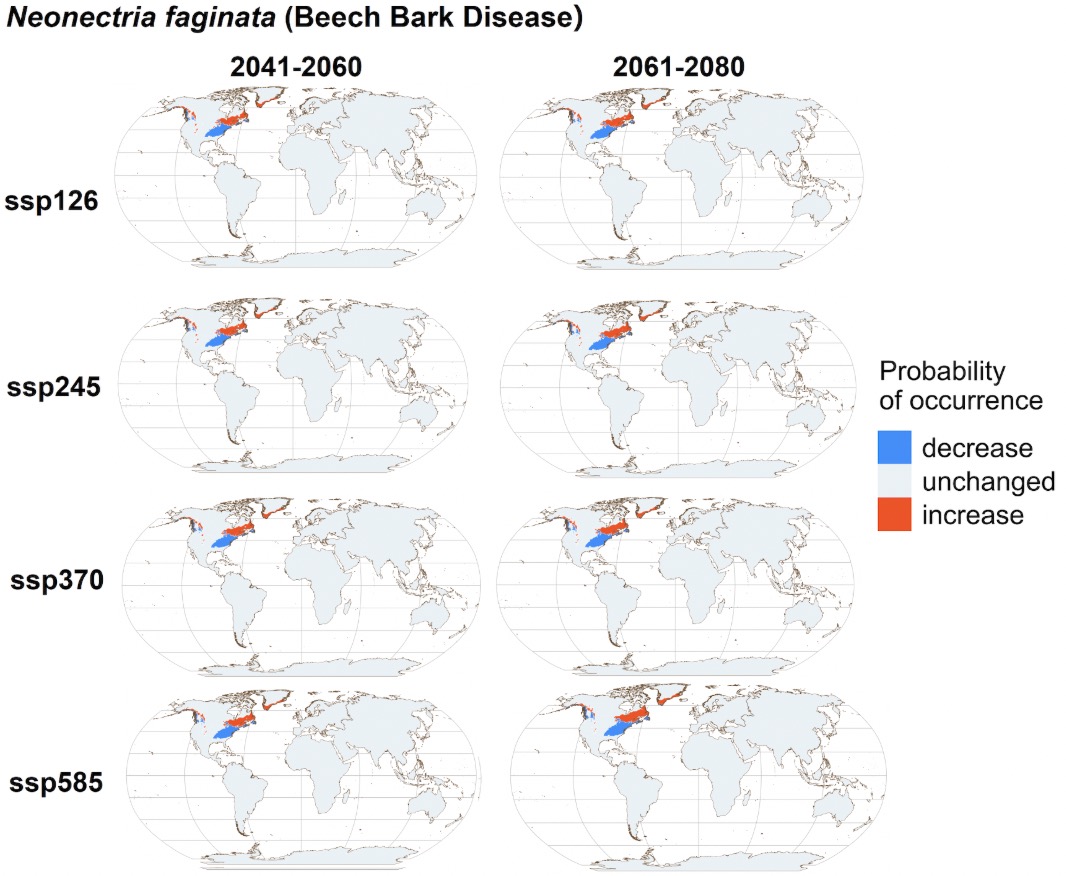

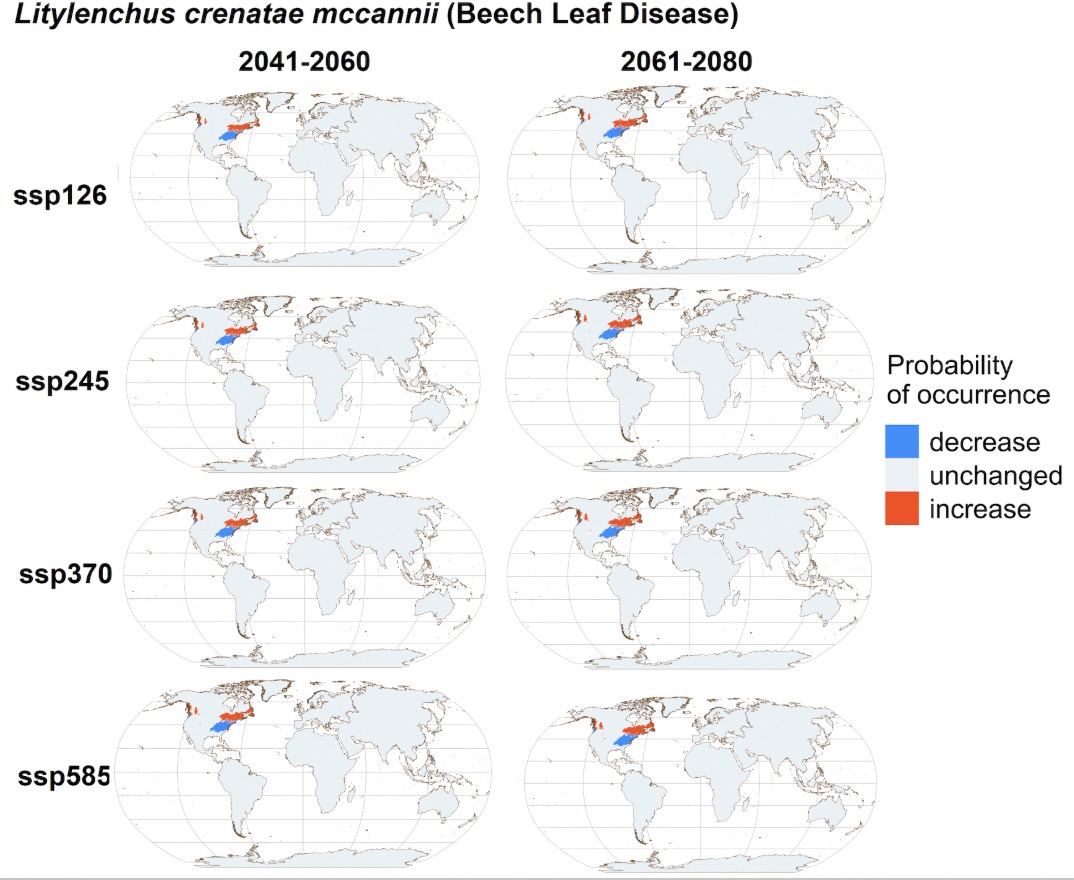

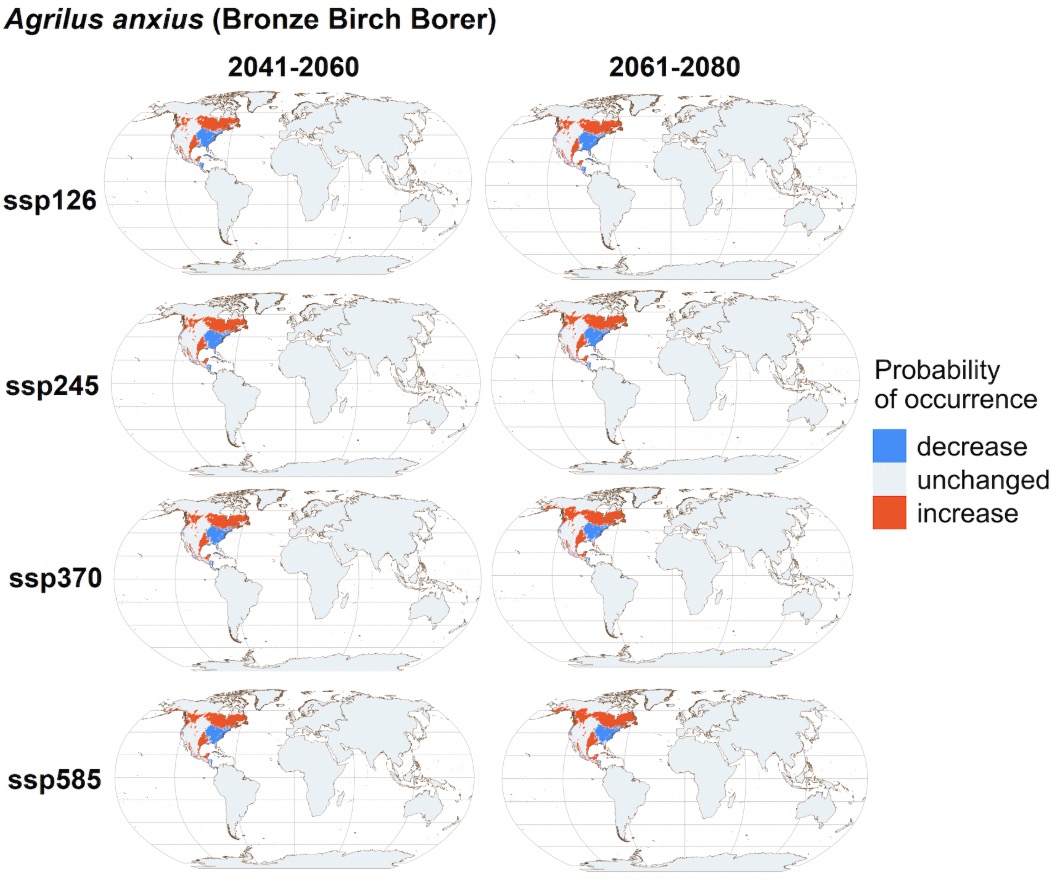

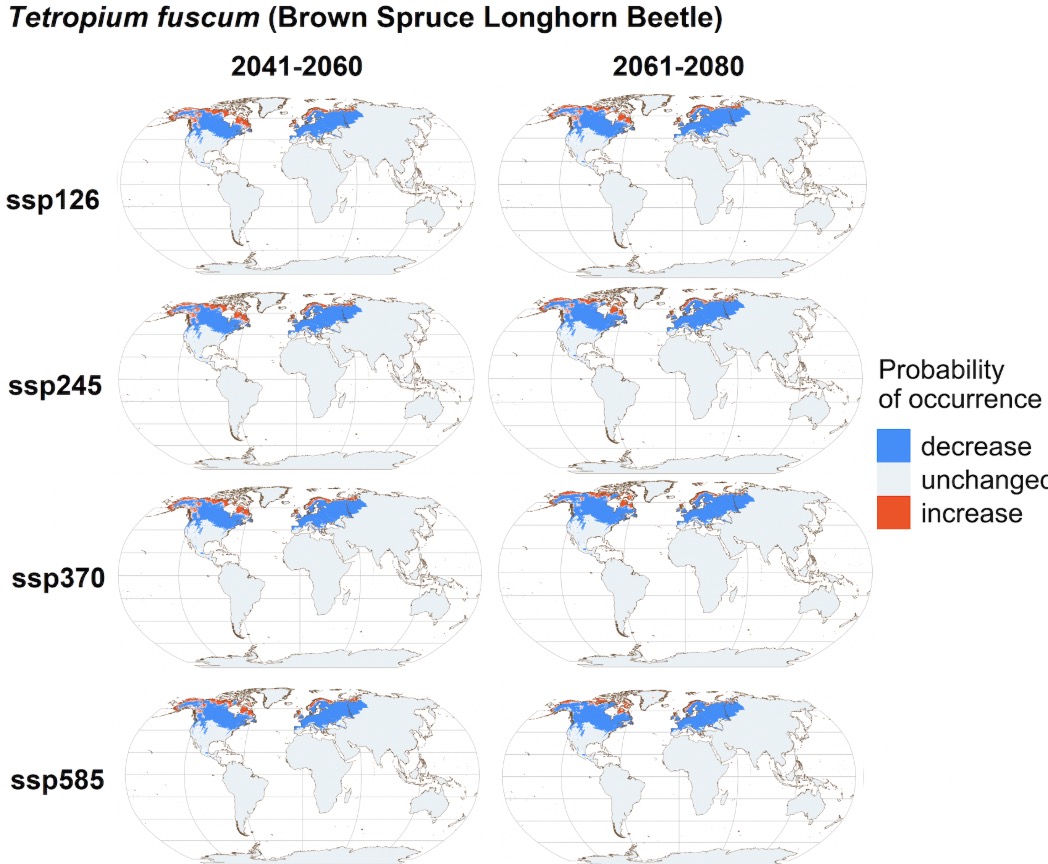

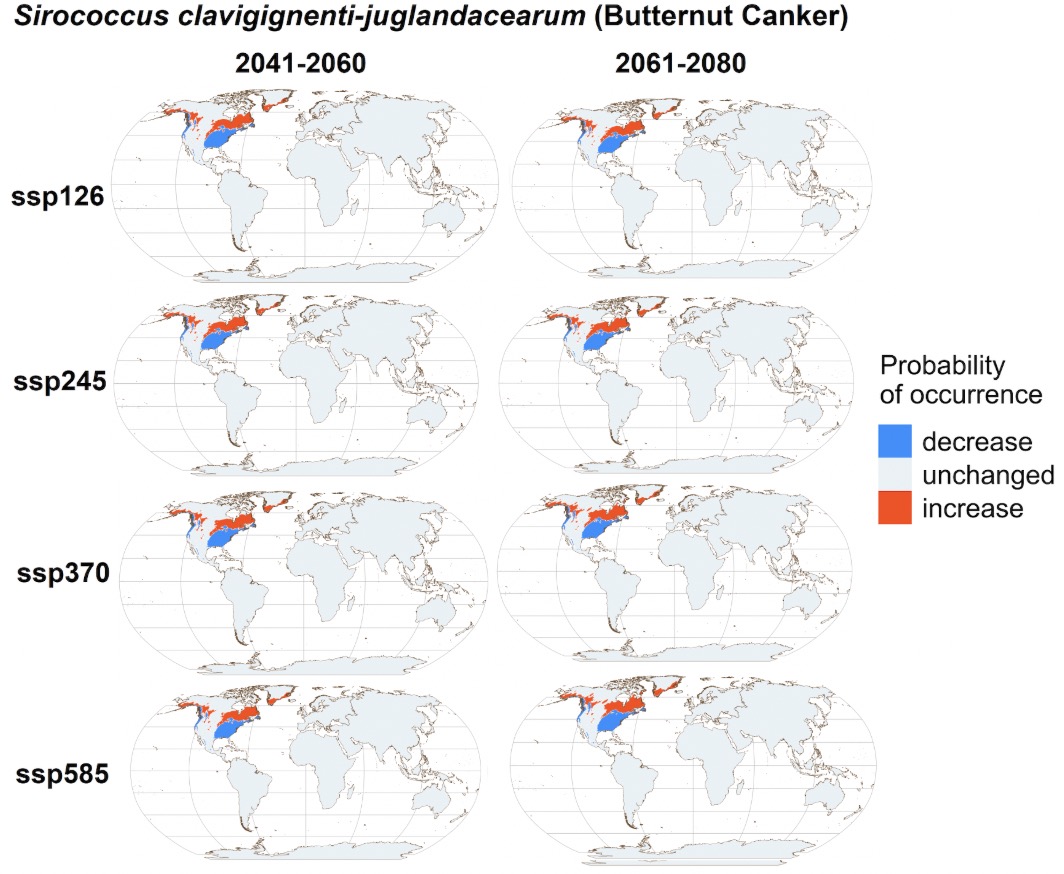

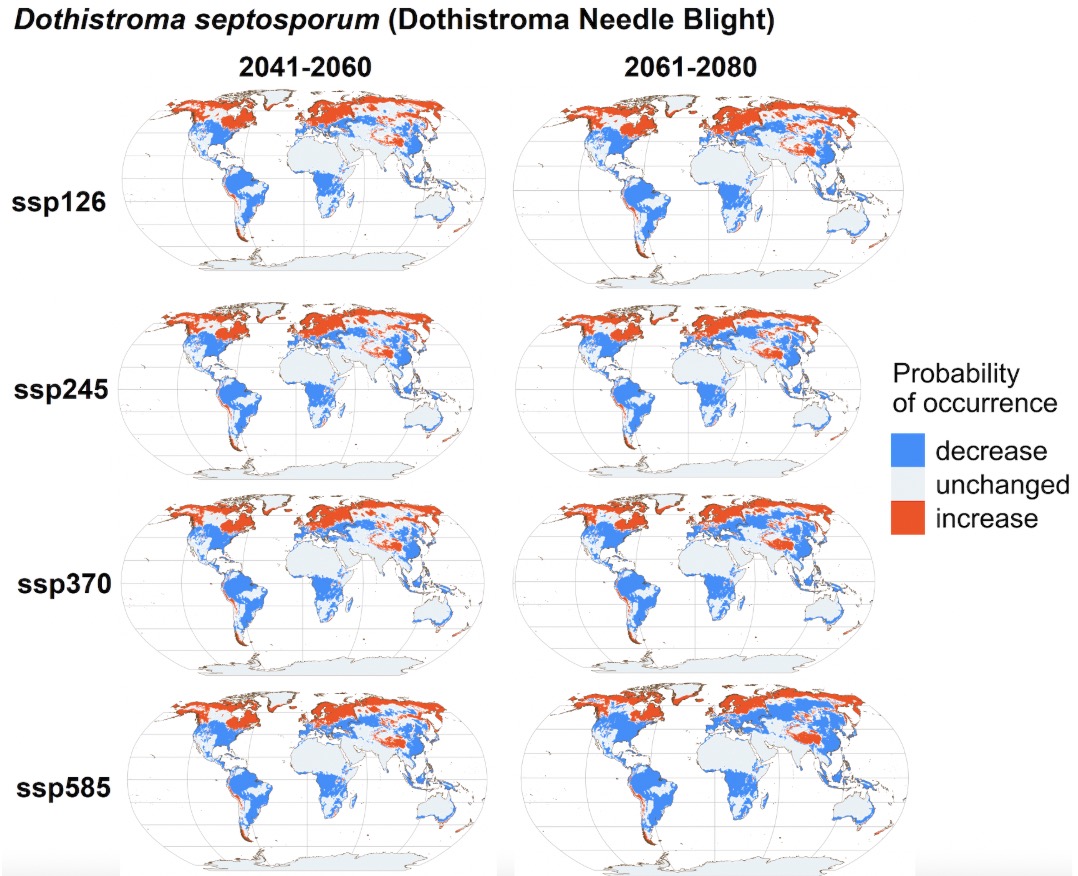

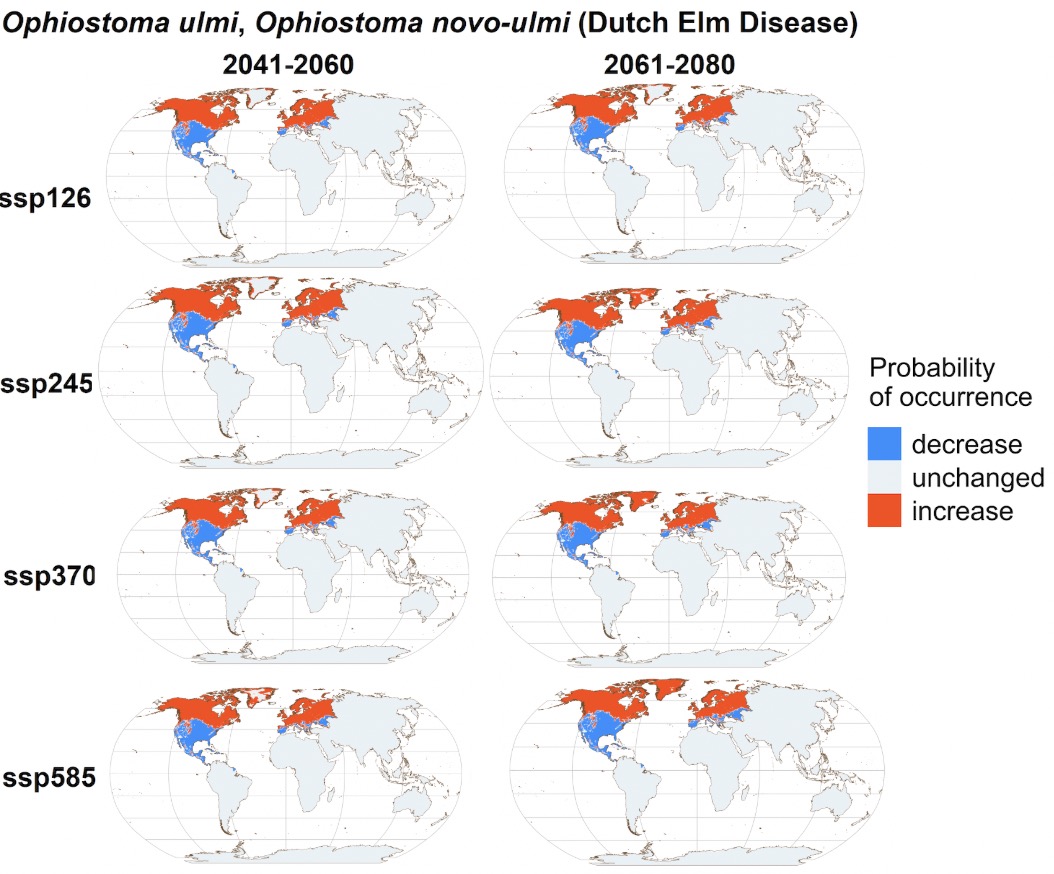

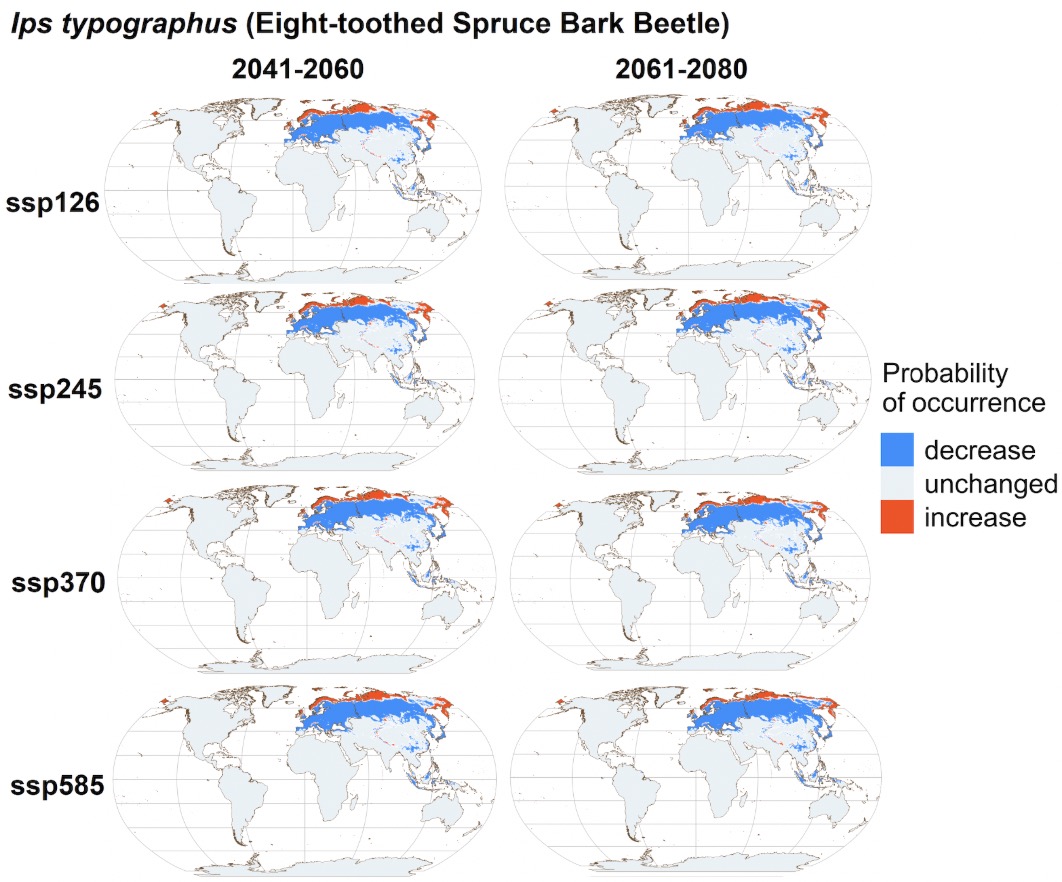

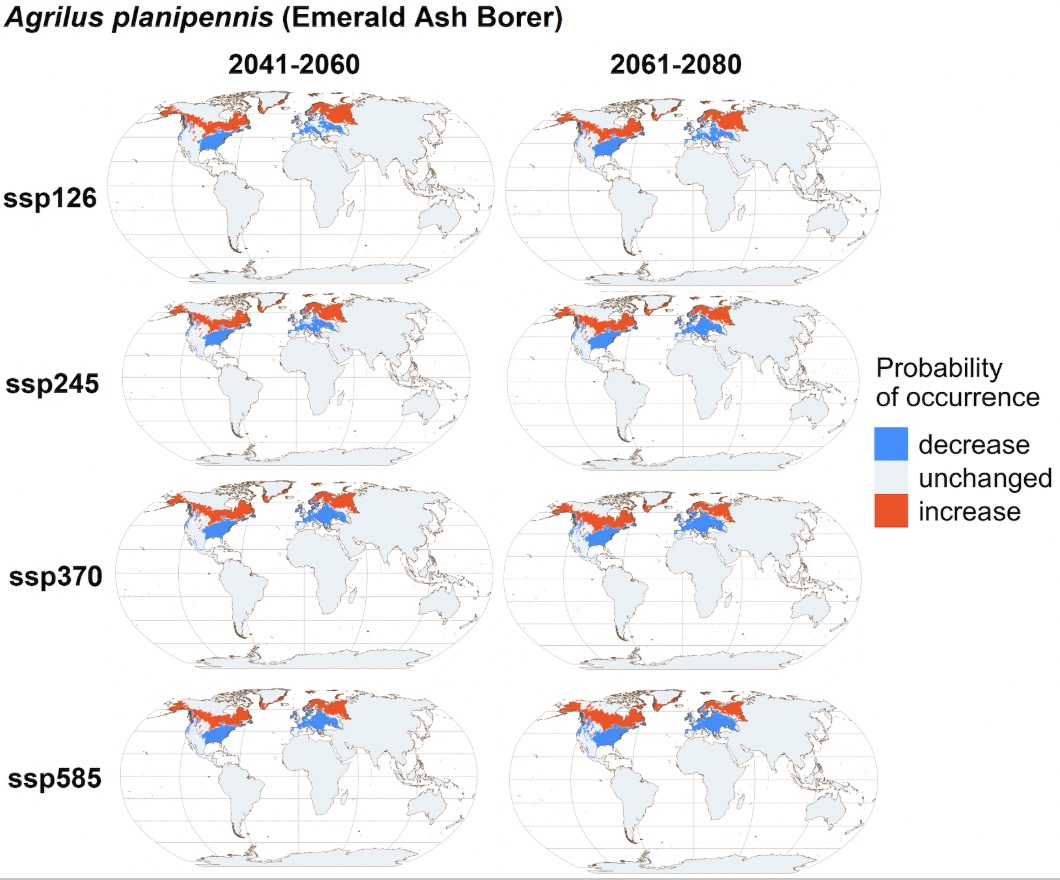

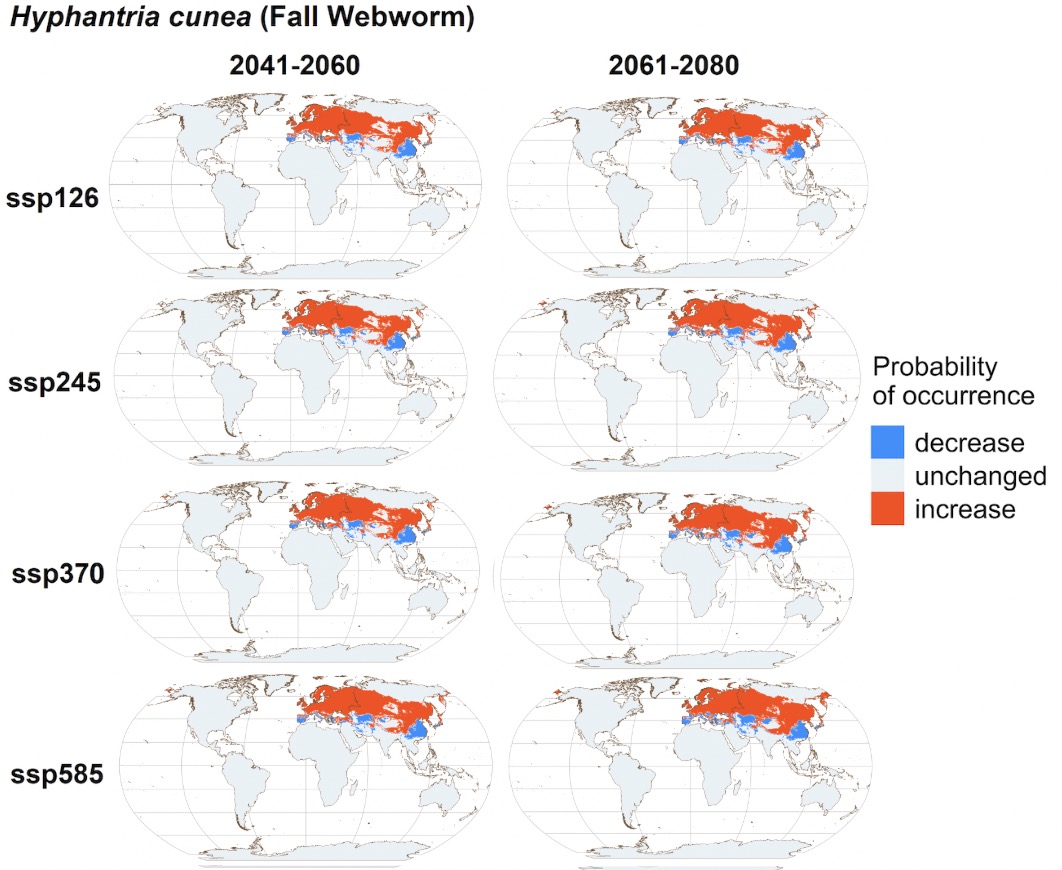

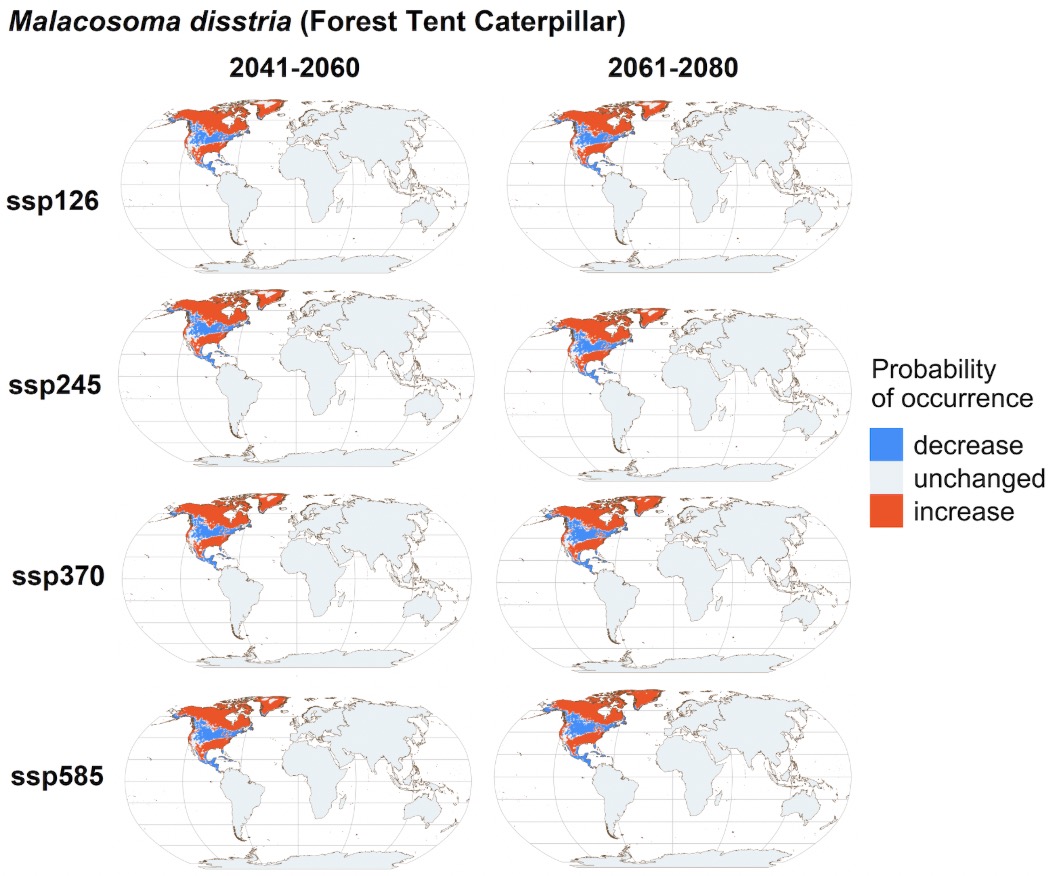

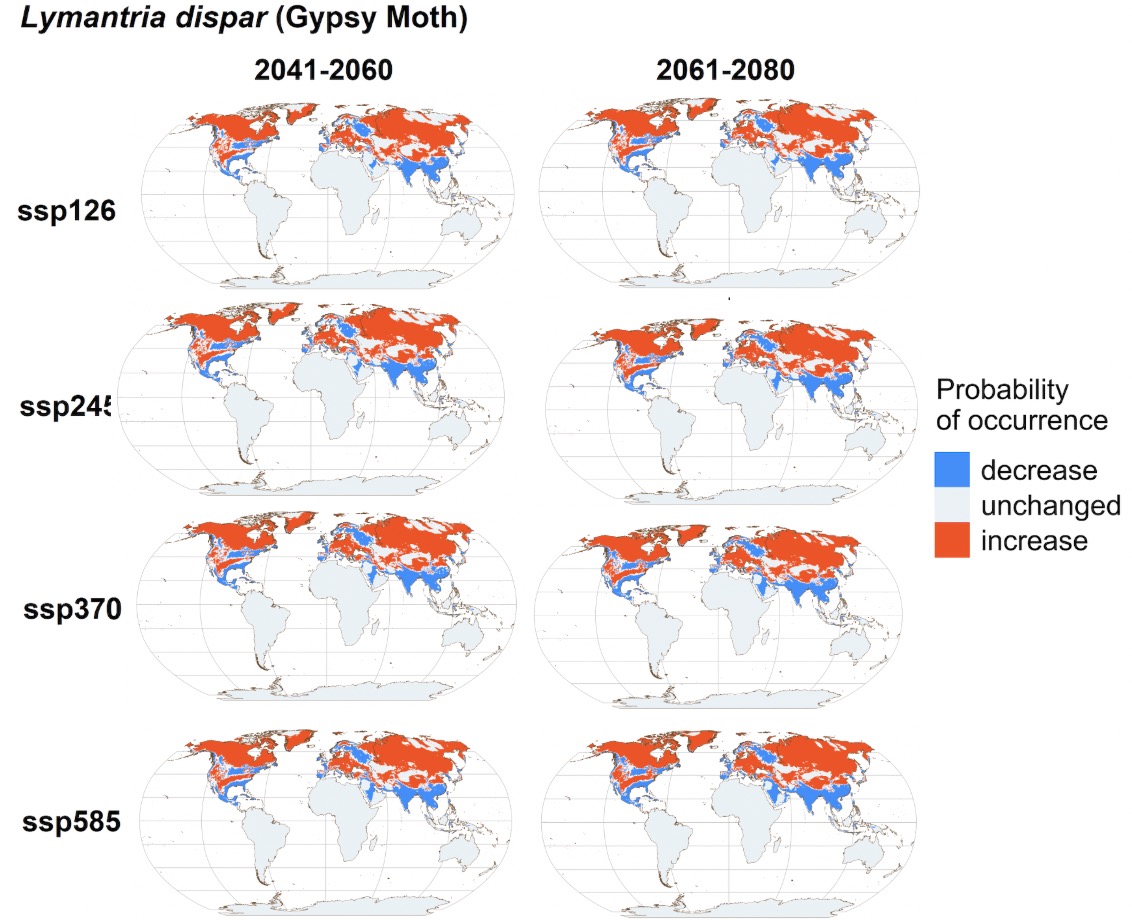

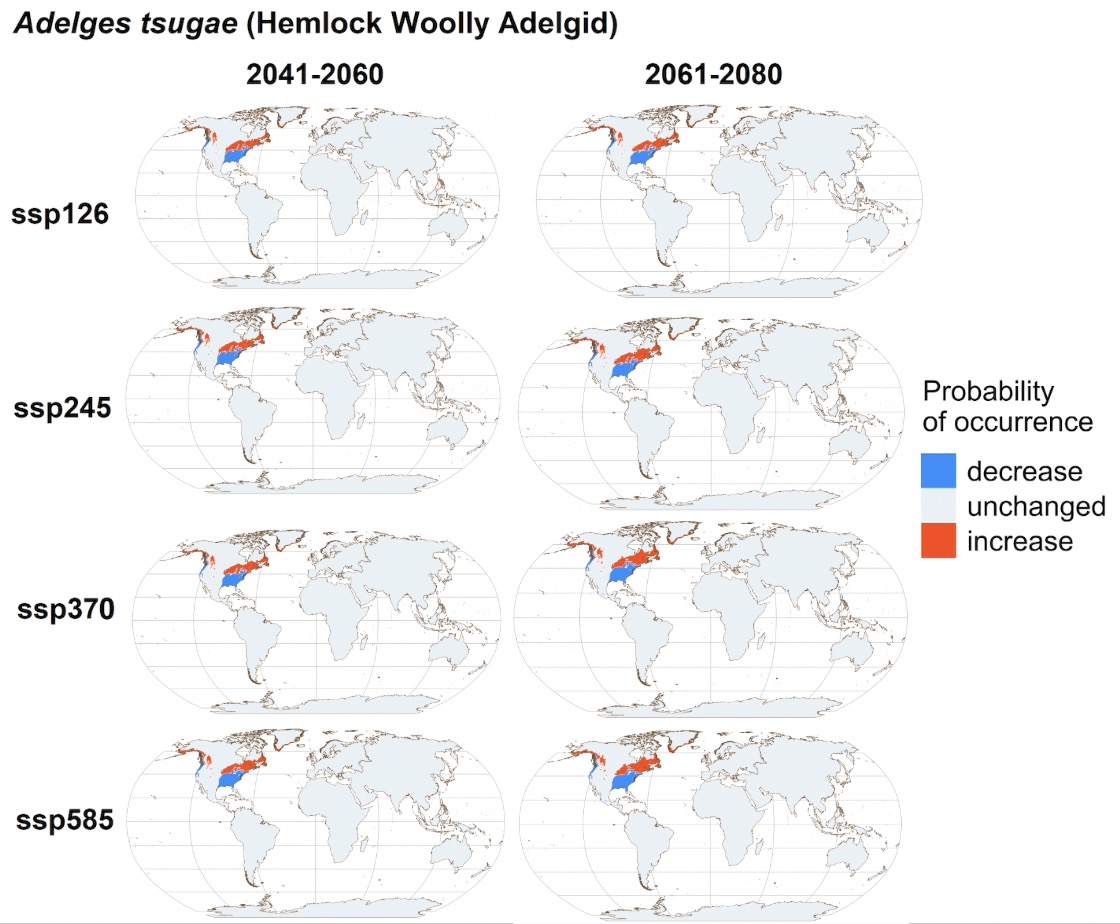

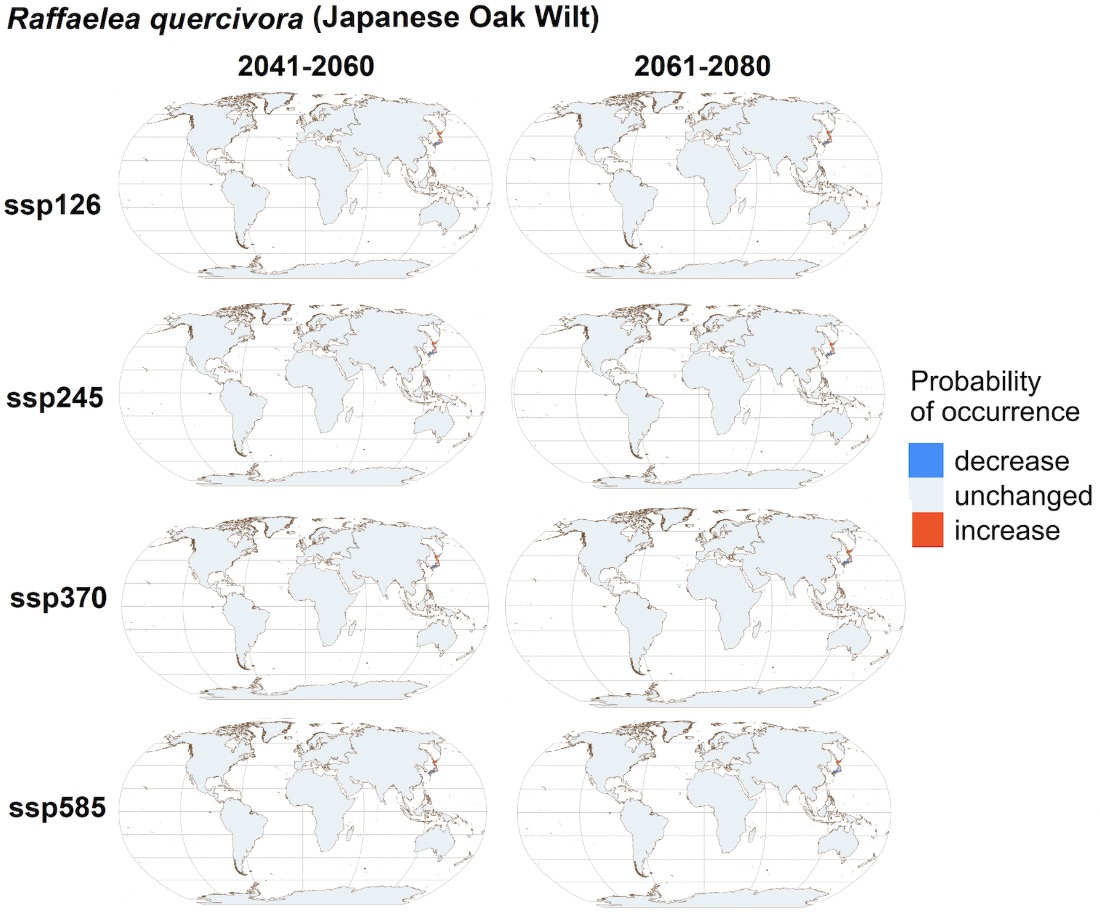

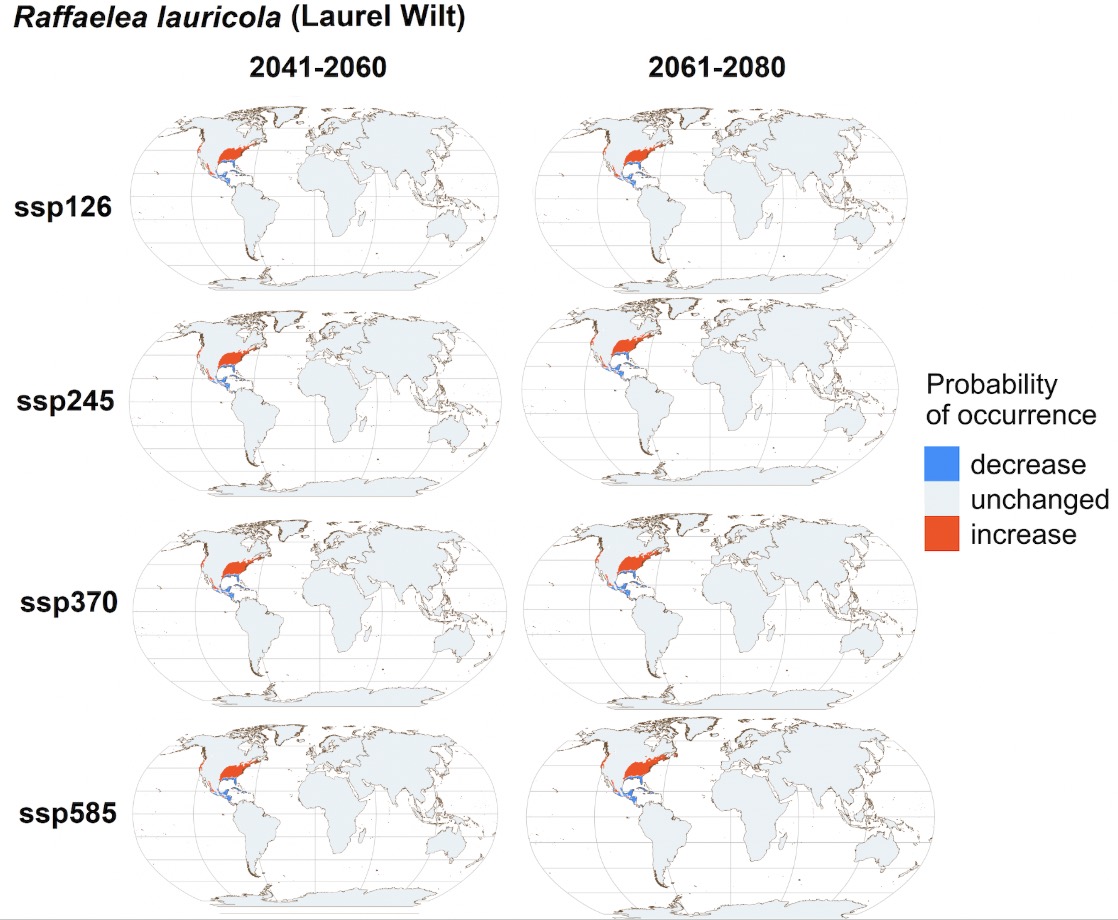

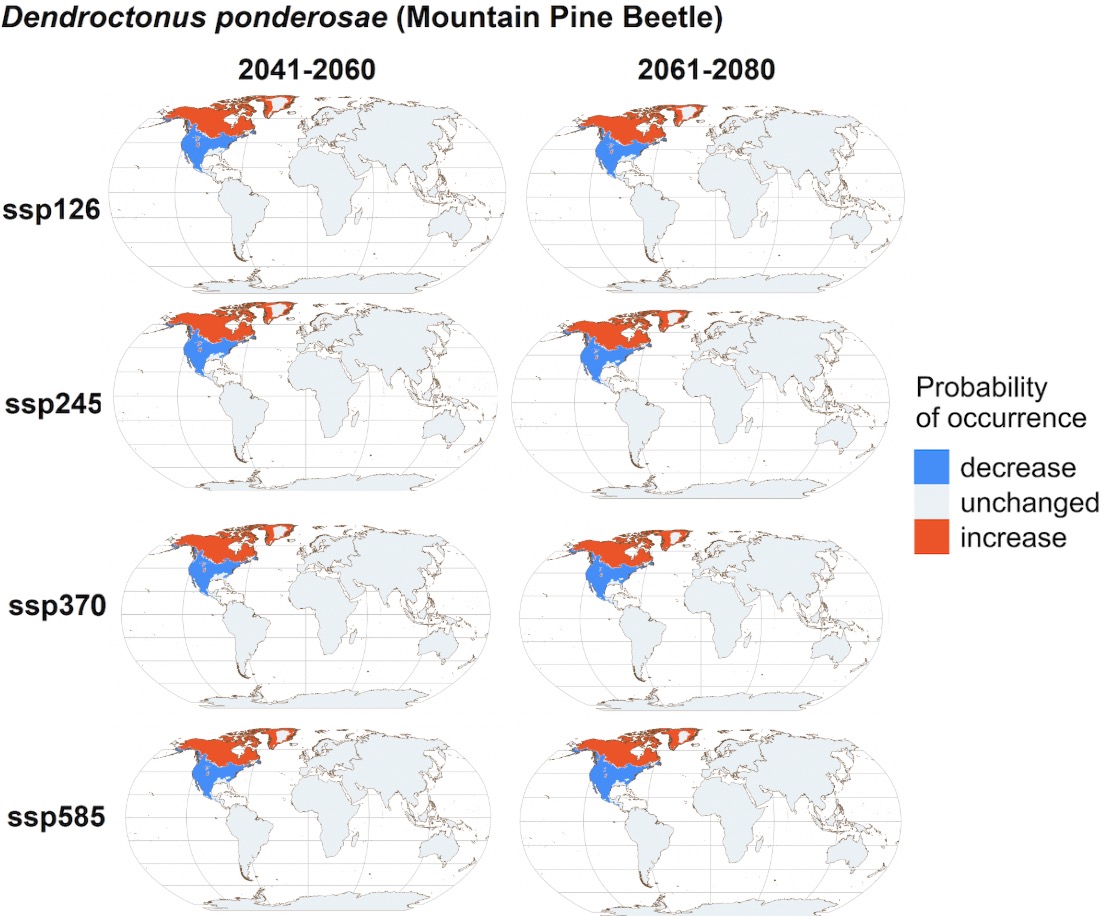

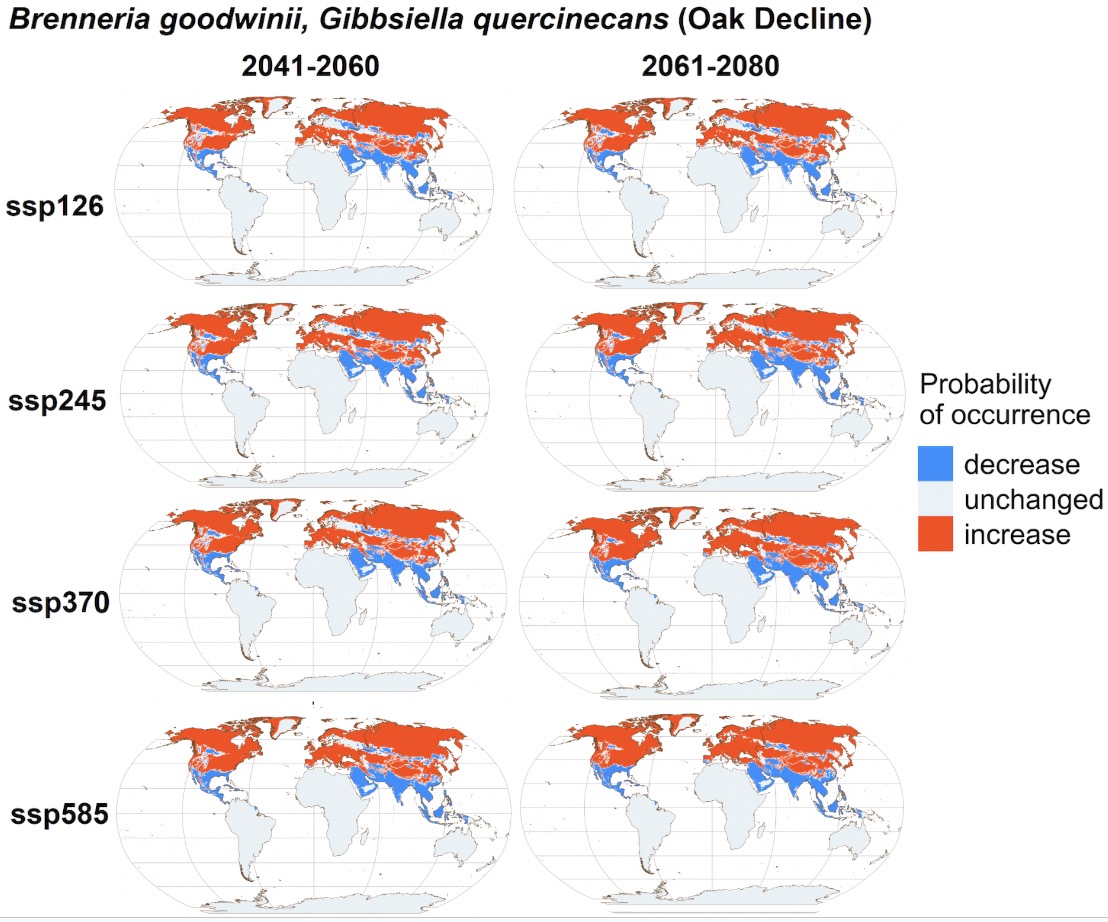

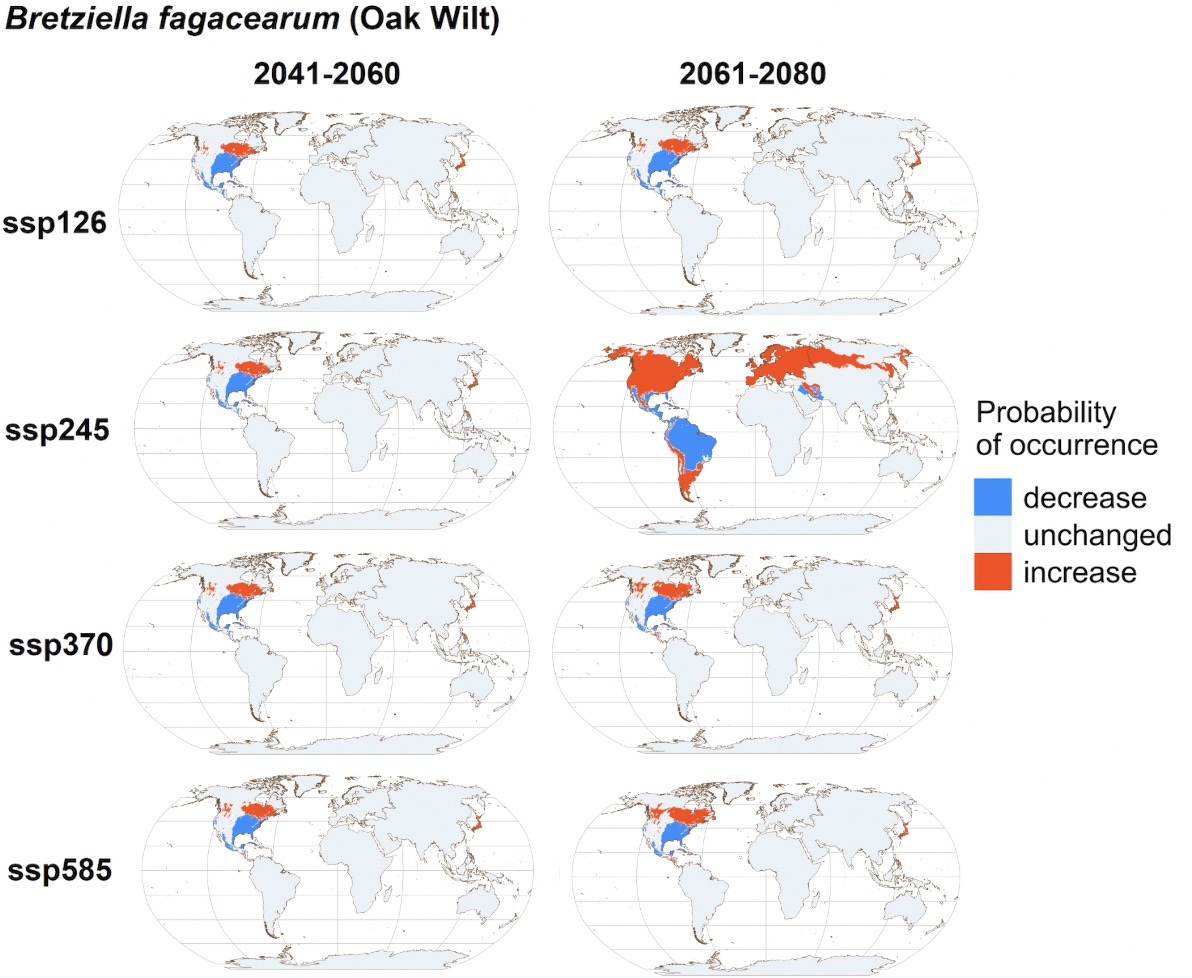

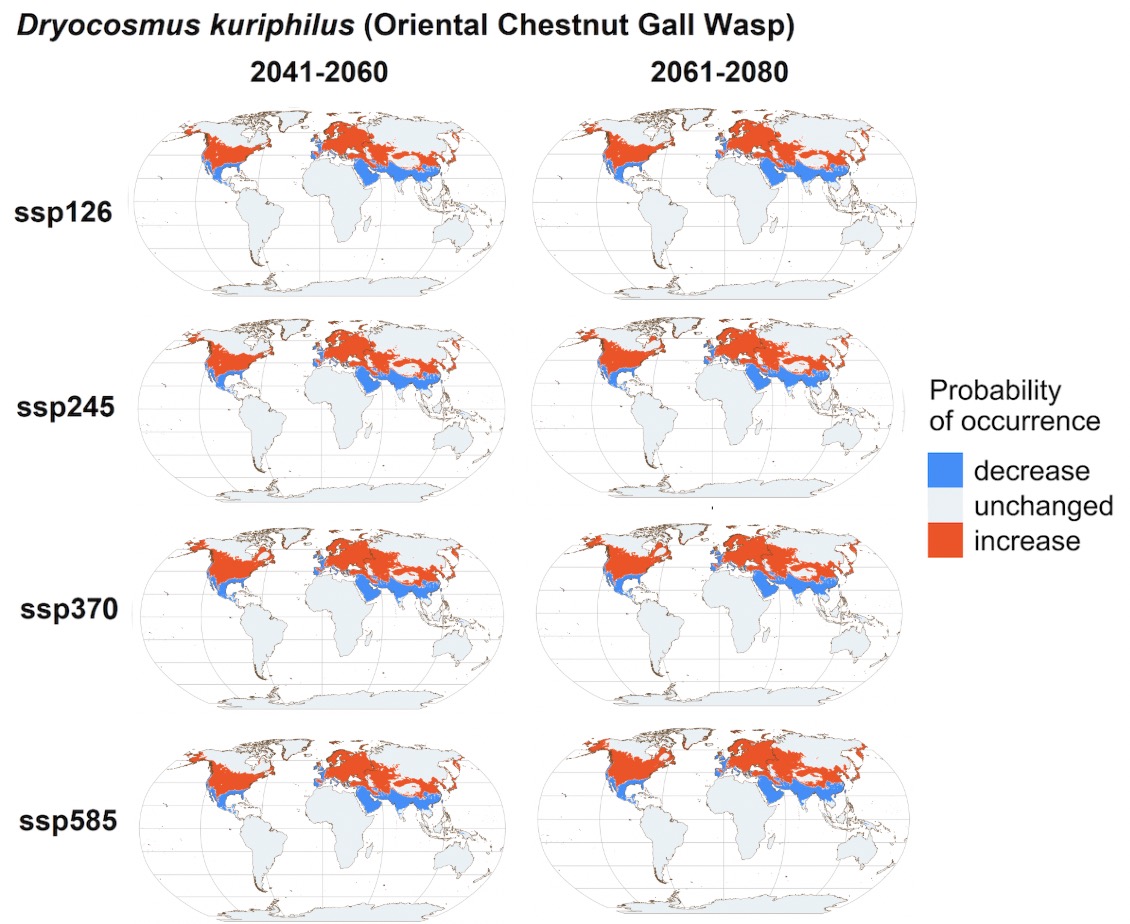

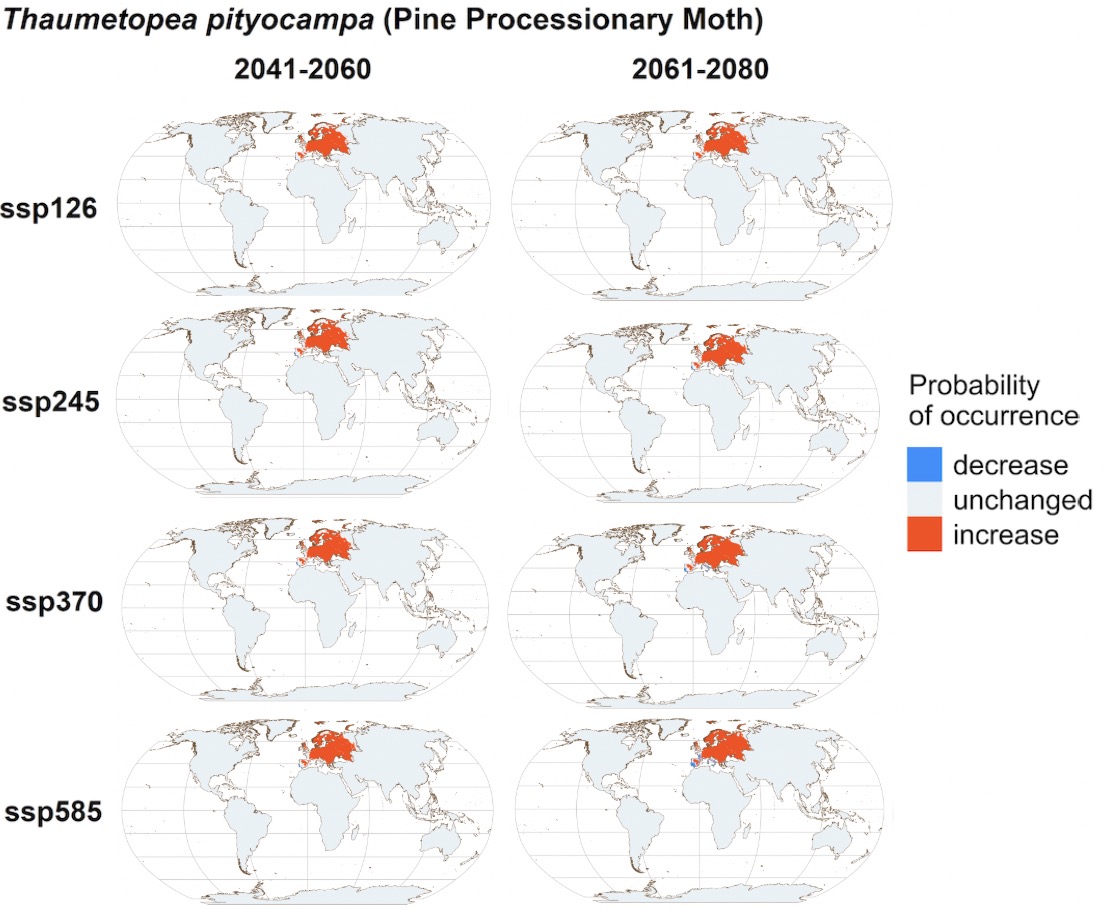

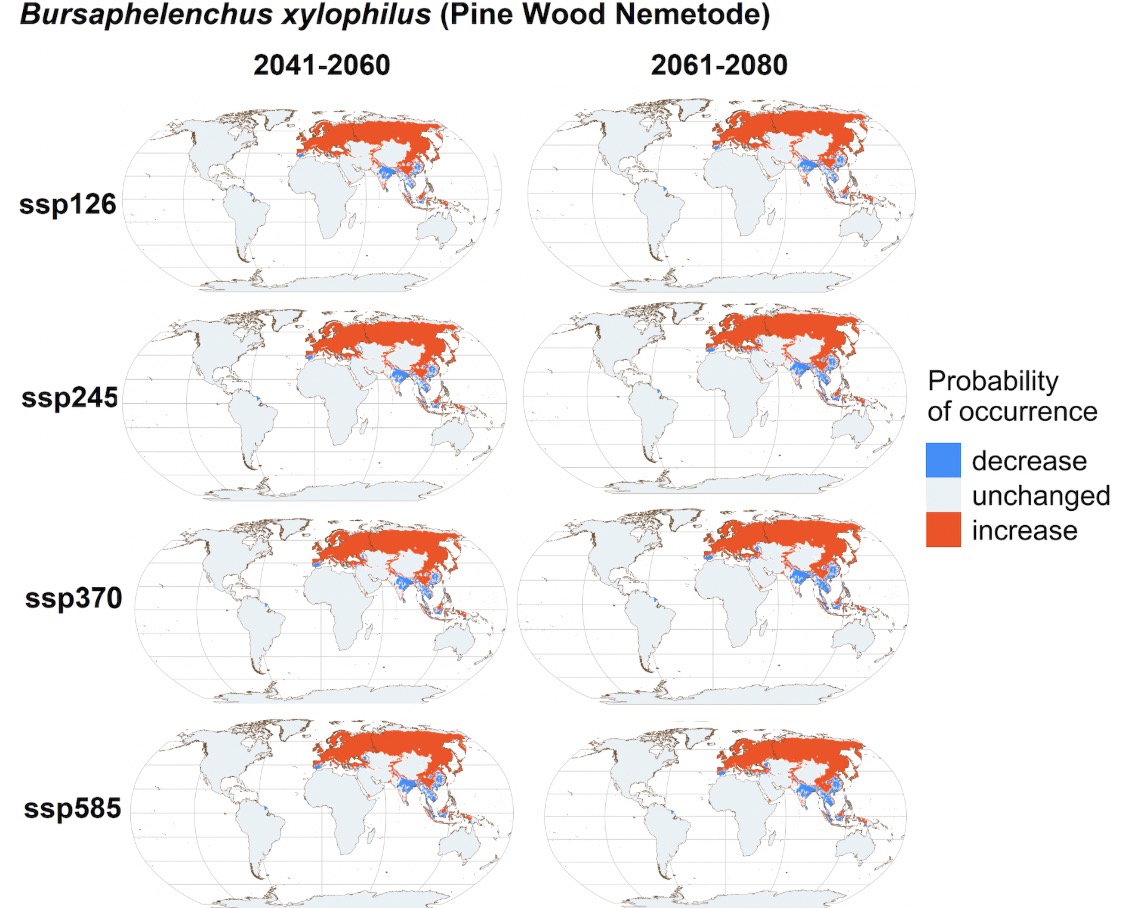

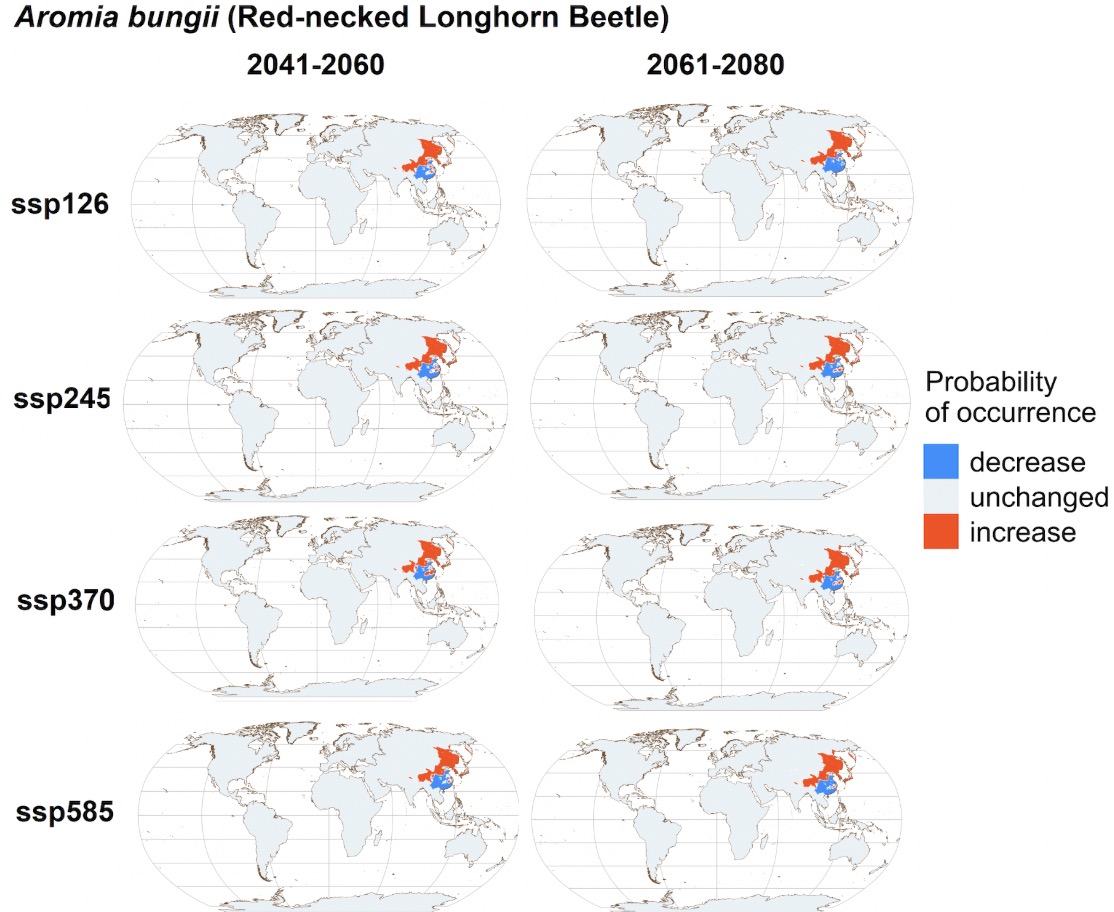

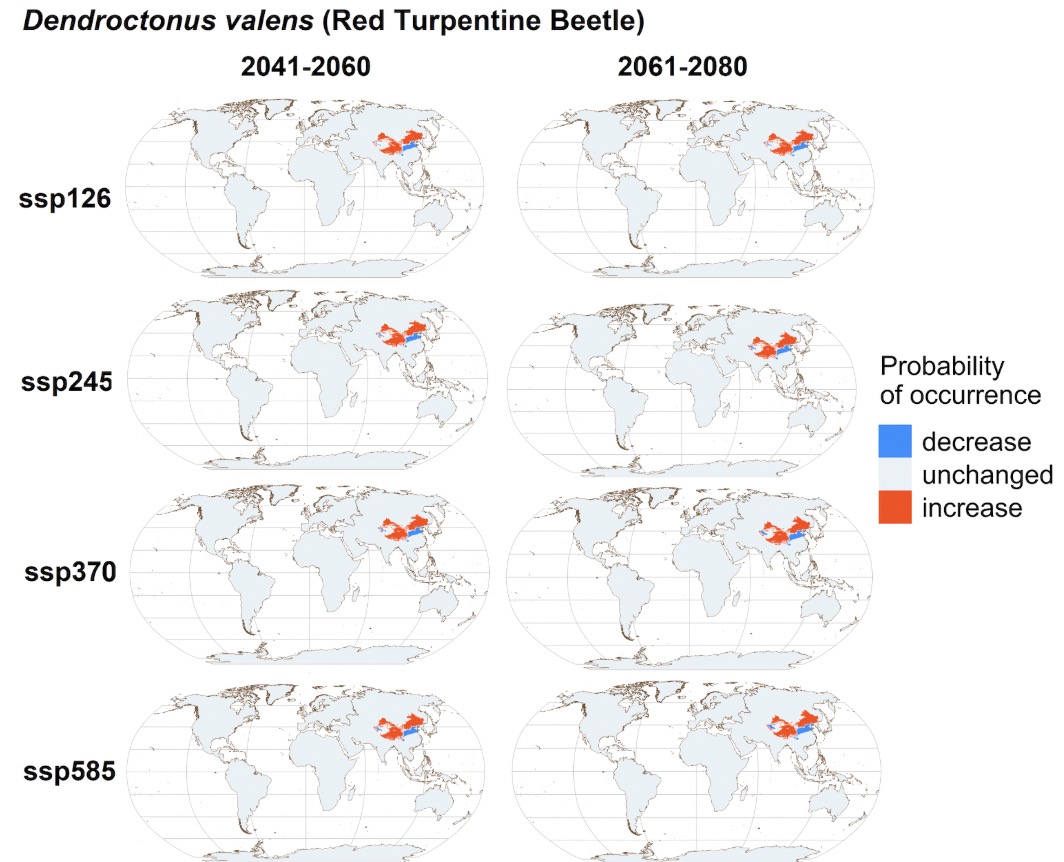

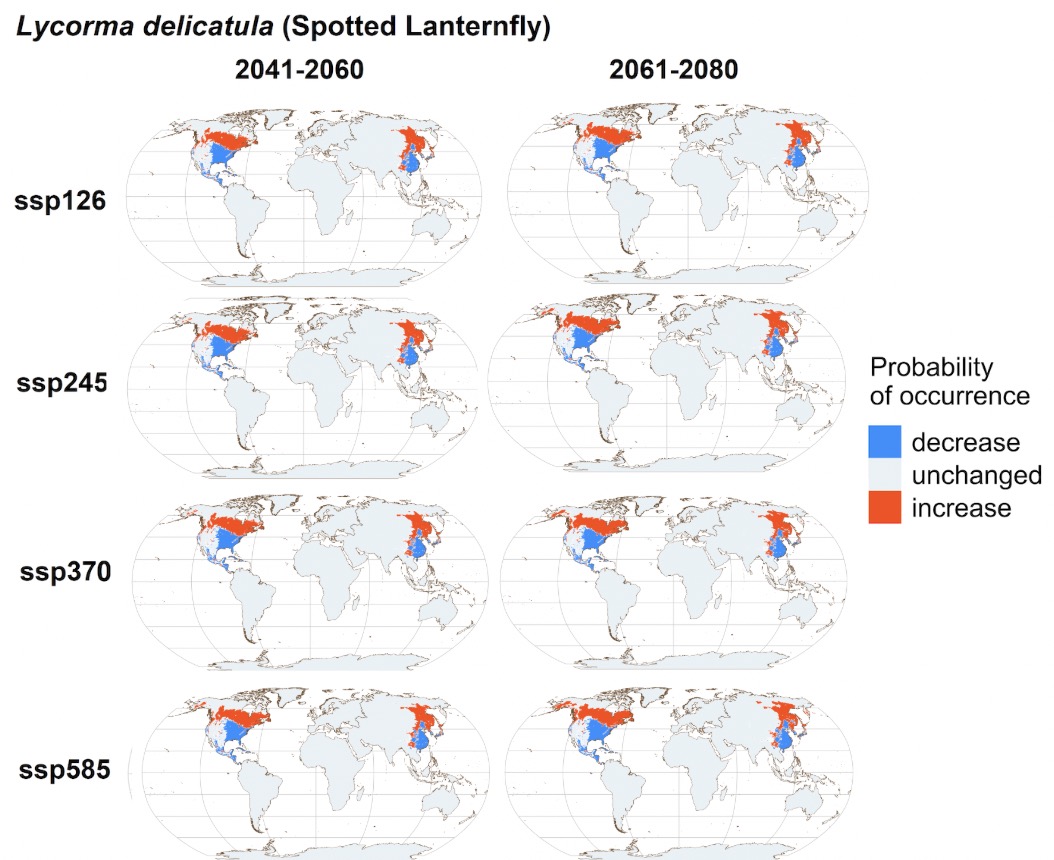

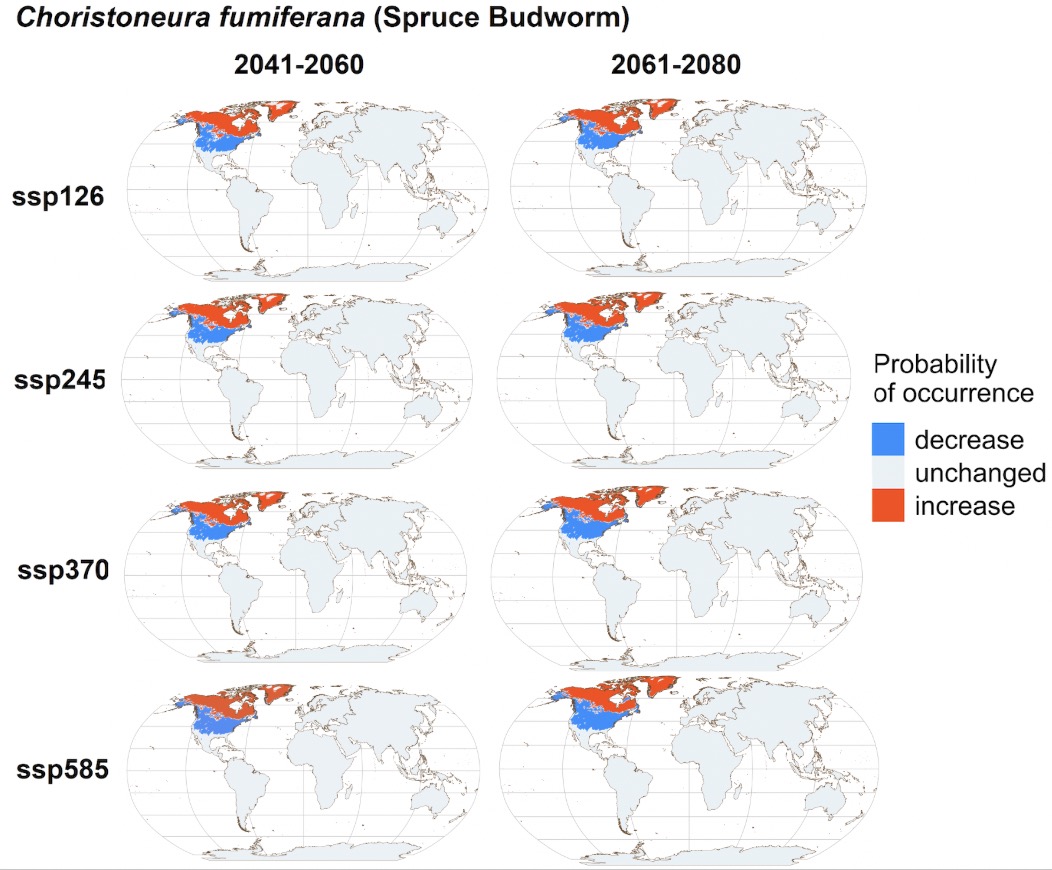

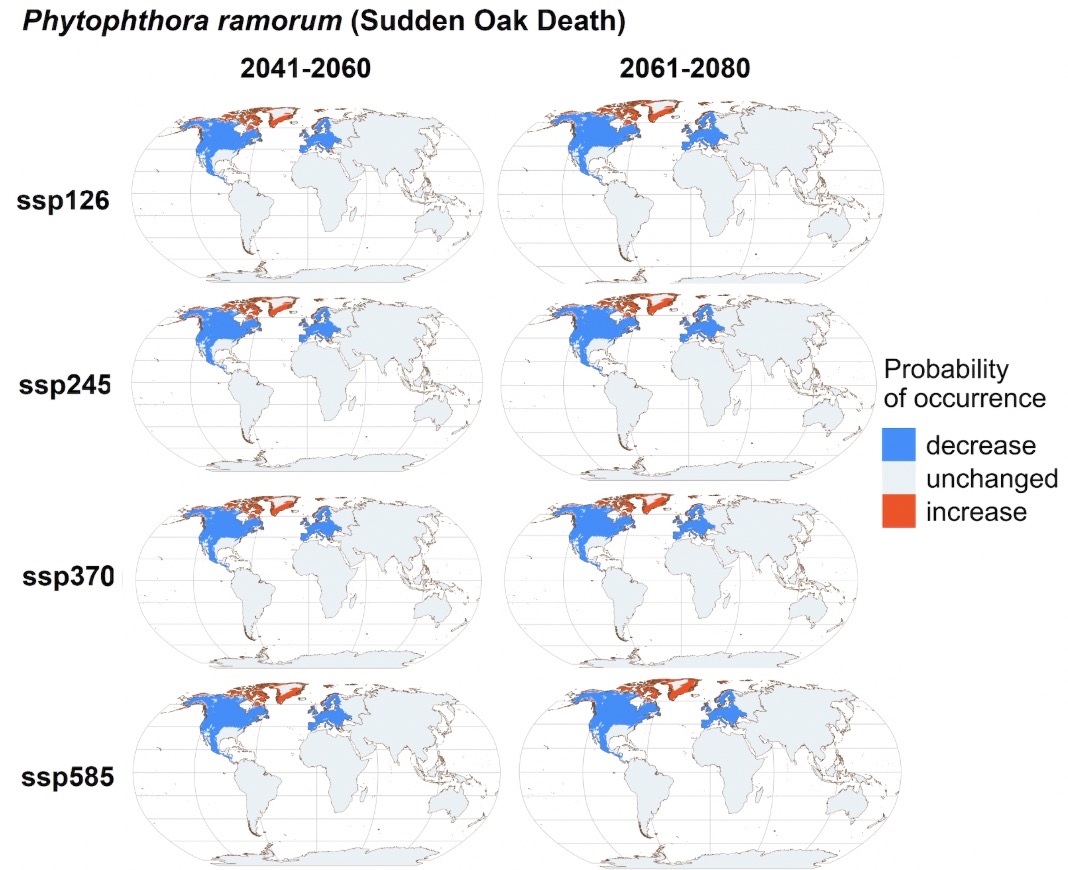

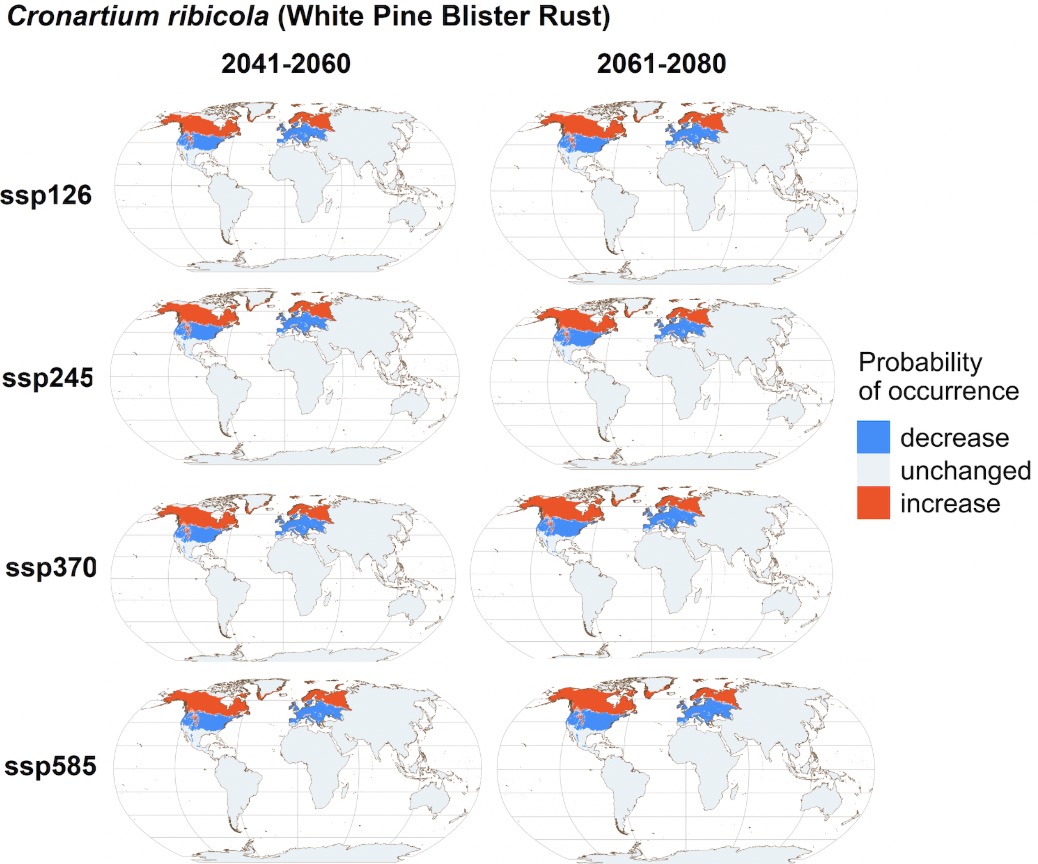

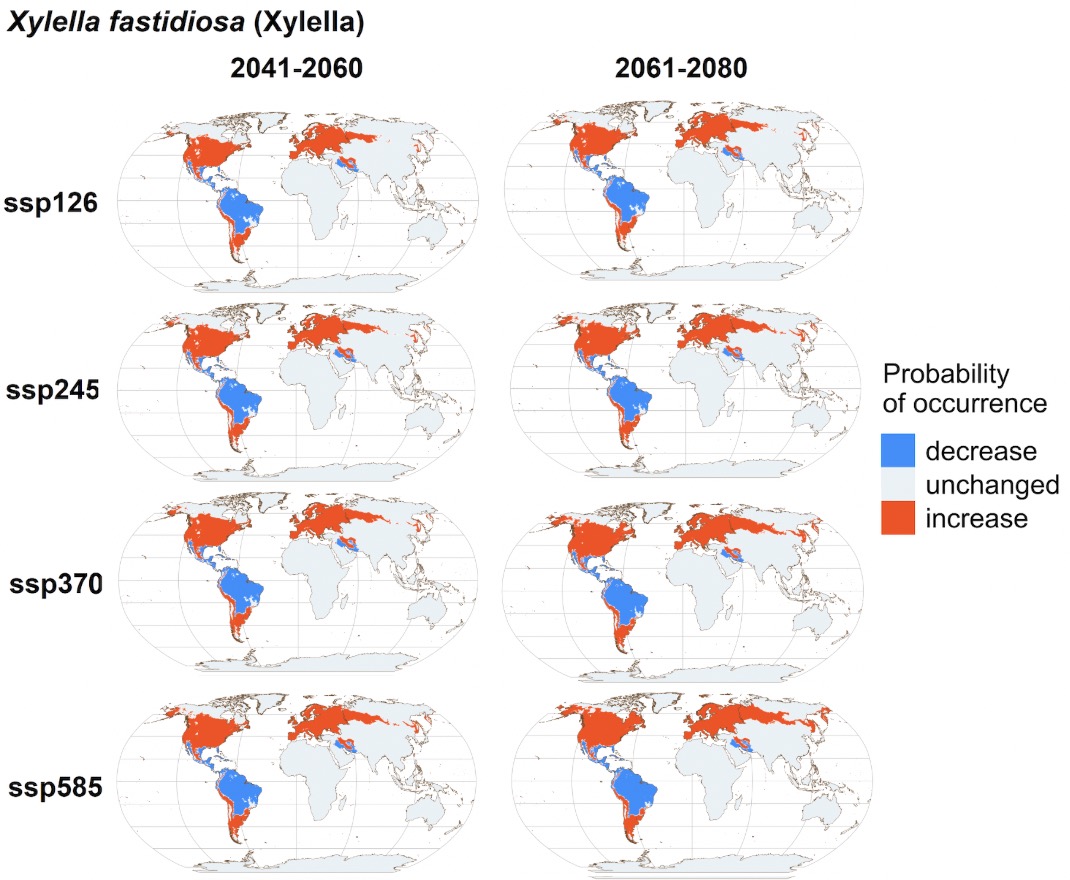

**Supplementary Figure S4.** Projections of changes in the probability of occurrence for 24 major forest pests and pathogens under four Shared Socio-economic Pathways (SSPs): SSP126, SSP245, SSP370, and SSP585. Regions where the change in pest or pathogen occurrence probability exceeds 0.01 are designated as “increase” (red-shaded areas), while those with a change of less than -0.01 are designated as “decrease” (blue-shaded areas). Note that the distribution maps for six major pests or pathogens (Ash Dieback, Asian Longhorned Beetle, Dutch Elm Disease, Emerald Ash Borer, Pine Wood Nematode, and Sweet Chestnut Blight) have been given in Figure 3.

### Figure S5. Changes in Pest and Pathogen Richness

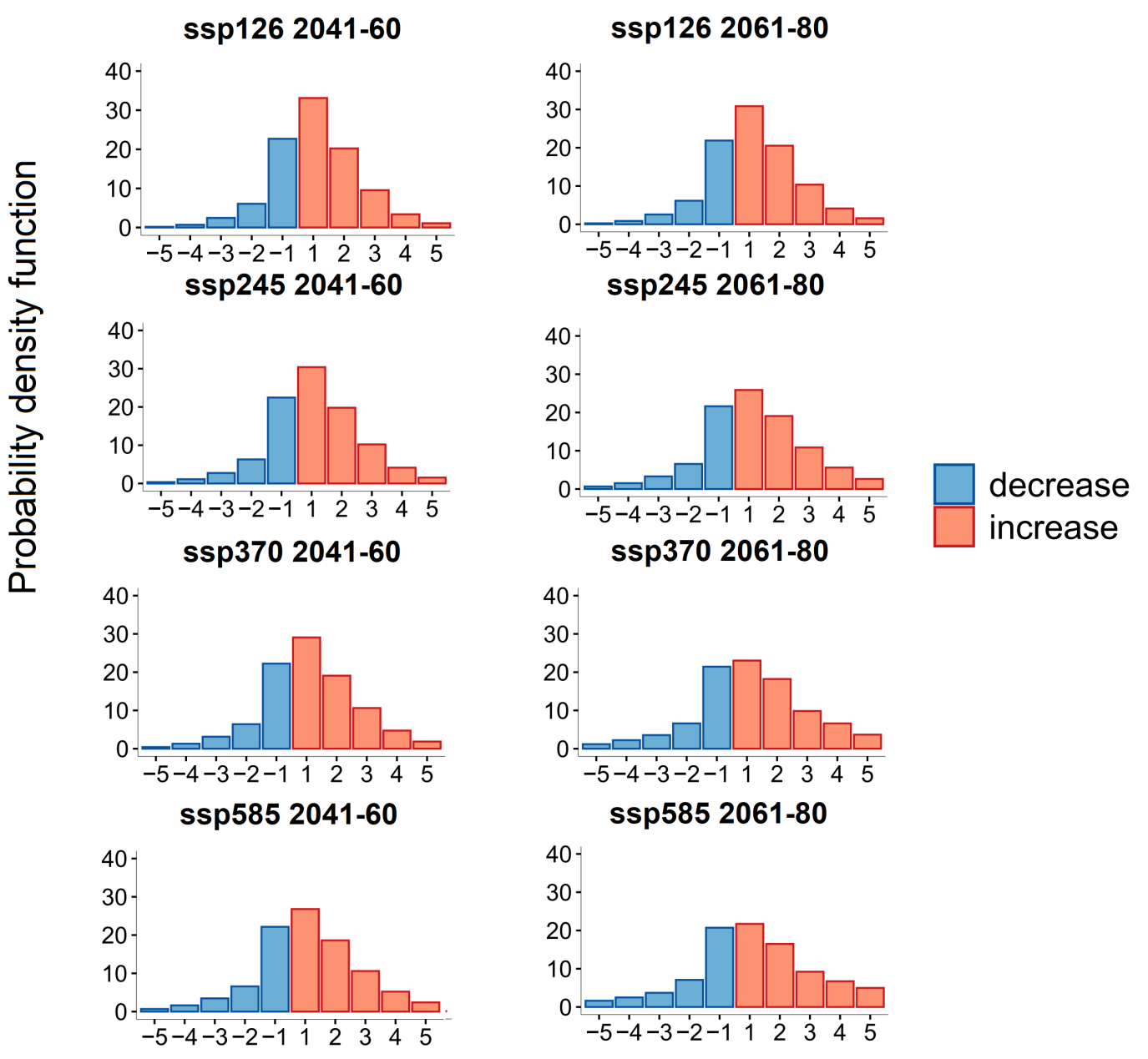

**Supplementary Figure S5.** Probability density functions of changes in pest and pathogen richness (number of pests and pathogens) for the years 2041–2060 and 2061–2080 under four Shared Socio-economic Pathways (SSPs): SSP126, SSP245, SSP370, and SSP585. The *y*-axis is the probability density (in percentage), indicating the relative likelihood of observing a certain change in pest and pathogen richness in an arbitrary location. Blue and red areas represent geographic regions with a decrease and increase in pest and pathogen richness, respectively. The sum of the areas of blue and red shaded regions equals to 1, the total probability, representing the complete range of possible changes in pest and pathogen richness.
